# Paternal genome elimination, monogenic reproduction, and the evolutionary genetics of atypical sex chromosome systems

**DOI:** 10.1101/2024.10.23.619029

**Authors:** Thomas J. Hitchcock, Robert B. Baird, Andy Gardner, Laura Ross

## Abstract

Sex chromosomes differ from autosomes in both their ploidy and transmission genetics. Consequently, selection, mutation, and drift may act differently upon them, driving distinct patterns in genetic divergence, diversity, and gene content. Recently, researchers have begun to consider a wider set of organisms with non-standard inheritance and sex-determination systems, however in many cases we lack theory which extends to such cases. One such example is paternal genome elimination (PGE), an unusual reproductive system which has independently evolved in two fly families, the fungus gnats (Sciaridae) and gall midges (Cecidomyiidae), and one order of springtails (Symphypleona). Under PGE, males receive but do not transmit a paternal genome, such that the autosomes and X chromosomes exhibit the same transmission genetics, but with different somatic ploidy. This makes them uniquely suited to test hypotheses about the role of haploid selection in males. Additionally, repeatedly throughout these groups a novel sex determination system – monogeny – has evolved, whereby females produce broods of exclusively one sex. The genetic basis of monogeny partitions the X chromosome into three segments, all displaying distinct inheritance patterns. Here we develop a series of theoretical models adapted to the genetics of these groups, generating testable predictions as to the relative genetic diversity within populations, and divergence between populations. Our results suggest that these species are excellent systems with which to test many fundamental principles in evolutionary genetics.

## Introduction

Natural selection, mutation, and random drift collectively shape the genomes and traits of all living organisms. For the better part of a century, researchers have been interested in how the balance between these forces may differ under different life cycles and inheritance systems (Gerstein and Otto 2009; Immler 2019). Possibly the best studied example of this – both theoretically and empirically – has been comparisons between sex chromosomes and autosomes. With their asymmetric genetic flow between the sexes, and sex-specific ploidy, sex chromosomes often experience selection coefficients, mutation rates, and effective population sizes that are distinct from those of the autosomes. This can generate a range of differences between the autosomes and sex chromosomes in terms of genetic diversity, rates of adaptation, and gene content (Haldane 1924; Avery 1984; Rice 1984; Charlesworth et al. 1987; Hedrick and Parker 1997; Vicoso and Charlesworth 2006; Meisel and Connallon 2013; Charlesworth et al. 2018).

In many well studied sex chromosome systems, the action of selection, mutation, and drift may simultaneously differ between autosomes and sex chromosomes. This confounding can make it challenging to identify the specific causes of differences (or the absence of differences) between portions of the genome, thus limiting the power of such comparisons (Jaquie ry et al. 2012; Charlesworth et al. 2018). However, in recent years new genomic technologies have allowed for the investigation of a much wider set of sex chromosome and inheritance systems across the tree of life (Bachtrog et al. 2011; Ross et al. 2022). Amongst this diversity, certain exceptional systems and ecologies may allow for more refined and powerful comparative tests of theory (Jaquie ry et al. 2012; Jaquie ry et al. 2013; Coelho et al. 2018).

From this perspective, three groups of interest are the dark winged fungus gnats (Sciaridae), the gall midges (Cecidomyiidae), and the globular springtails (Symphyleona). These groups all have X chromosomes, and all share an autosomal inheritance system - paternal genome elimination (PGE) - in which males receive but do not transmit genes from their fathers (Box 1; Herbette and Ross 2023). This leads autosomes and X chromosomes to exhibit the same transmission genetics but with distinct somatic genetics. Secondly, various species within these groups have also evolved a novel sex ratio strategy – monogenic reproduction – whereby females produce single sex broods (Box 2; Baird *et al*. 2023b). The genetic basis of this monogeny differs amongst groups. In fungus gnats, it is known to be caused by an X-linked daughter-producing supergene, thus resembling a nested sex chromosome system upon the X chromosome (Baird, Urban, et al. 2023). Conversely, in gall midges, a similar supergene system appears to exist, albeit located on the autosomes (Stuart and Hatchett 1991; Benatti et al. 2010). Both these phenomena may be expected to result in autosome/sex chromosome comparisons which are distinct from – and potentially clearer than – classic systems. But whilst there has been verbal discussion of these points (De La Filia et al. 2015), we currently lack concrete models with which to generate predictions about, and interpret data from, these systems.

To address this shortfall in understanding, here we develop theory applicable to X/PGE systems. Firstly, we describe how different portions of the genome may be expected to flow between different states across evolutionary time, which we frame in terms of the concept of class reproductive value, and we use this to calculate the effective marginal fitness effects, population sizes, and mutation rates under these different inheritance systems. We then generate results concerning the genetic diversity within populations and genetic divergence between populations, exploring how such results are expected to differ between PGE and “eumendelian” systems (Normark 2006), and across different portions of the X chromosome (and autosomes) under monogenic sex determination. Collectively, these results not only help to guide future empirical work on these species but also highlight these groups as being exceptional systems with which to test evolutionary theory.

### Class reproductive value modulates effective fitness effects, mutation rates, and population size

Over generations, gene lineages move between different classes of individual. Examples include sexes, habitats, and ages. The specific pattern of flow ultimately shapes the amount of time that a gene lineage spends in a particular class and thus the average mutational, selective, and demographic environment it experiences. The proportion of time a gene lineage spends in a specific class defines the reproductive value of that class (Fisher 1930; Price and Smith 1972; Taylor 1990; Grafen 2006; Lehmann 2014). For example, in a classic autosomal system, with no overlapping of generations, gene lineages spend an equal fraction of their time in males and females (*c*_f_ = *c*_m_ = 1/2) (Taylor 1996). In contrast, for a standard X chromosome (or haplodiploid) system, gene lineages will, on average, spend twice as much time in females as they will in males (*c*_f_ = 2/3, *c*_m_ = 1/3)(Taylor 1996). This is also the same for paternal genome elimination (PGE, Box 1), provided there is strictly no male transmission of paternal-origin gene copies (Hitchcock et al. 2022).

We can determine class reproductive values for the more complex genetic systems of interest here. We consider three states: female-producing females (gynogenic females, f_𝕘_), male-producing females (androgenic females, f_𝕒_), and males (*m*) (see Box 2 for an illustration). We describe the different ways that a gene lineage may flow between these states using a transition matrix *T*. The left-eigenvector associated with *T* provides us with the class-reproductive values of these various states (Taylor 1990). We notate the class reproductive value of class *i* as *c*_i_. These reproductive values can be further decomposed to consider the frequency of movement between states *c*_i→j_ (e.g. androgenic females to males, or males to gynogenic females), which are also known as “elasticities” in demography (Caswell 2018; Giaimo 2022). The class reproductive values for different portions of the genome are given in Table 2, and a full description of methods is given in SM§1.

**Table 1:** Verbal summary of results for different types of X/autosome comparisons. Mathematical expressions can be found in the main text and supplementary material.

| | | X vs A | X vs PGE | $X_a$ vs PGE |
| --- | --- | --- | --- | --- |
| Genetic Diversity | Neutral | Autosomes generally higher, but X may be higher when there is high male reproductive skew. | Equal | Higher under PGE, except when there is much higher skew in androgenic females. |
|  | Deleterious | Autosomes generally higher, but may be lower for dominant, male biased genes. | Always higher under PGE | Always higher under PGE |
| | Sexually antagonistic | Higher on X if equal dominance, higher on autosomes if reversals of dominance. | Higher on X if equal dominance, higher under PGE if reversals of dominance | Higher on $X_a$ if equal dominance, higher under PGE if reversals of dominance |
| Genetic Divergence | Beneficial – Hard | Higher on X when mutations are recessive, higher on autosomes when dominant. | Always higher on the X. | Higher on $X_a$, except for strongly dominant mutations. |
|  | Beneficial – Soft | Higher on autosomes under equal dominance. May be higher on X for recessive beneficial mutations with dominance reversals. | Equal under equal dominance. With dominance reversals, higher under PGE for dominant beneficial mutations, higher under X for recessive beneficial mutations. | Higher under PGE, except for recessive beneficial mutations under dominance reversals. |
|  | Neutral | Equal, unless sex biases in the mutation rate | Equal | Equal, unless sex biases in mutation rate. |
|  | Deleterious | Higher on autosomes if recessive, higher on X if dominant. Sensitive to sex-biases in demography. | Higher under PGE. | Higher under PGE. |

**Table 2:** Class reproductive values under different modes of inheritance, where λ is the probability that a gene copy in a haploid descends from a male. These are presented for different genetic systems under digeny (D) and monogeny (M).

| | | Gynogenic females<br>$c_{f_g}$ | Androgenic females<br>$c_{f_a}$ | Males<br>$c_m$ |
| --- | --- | --- | --- | --- |
| Diploid | D | 1/2 |  | 1/2 |
|  | M | 1/4 | 1/4 | 1/2 |
| X-linked/<br>PGE | D | 2/3 |  | 1/3 |
|  | M | 1/3 | 1/3 | 1/3 |
| Cytoplasmic | D | $1 - \lambda$ | | $\lambda$ |
| | M | $(1 - \lambda)^2$ | $(1 - \lambda)\lambda$ | $\lambda$ |
| $X_a$ | M | 1/5 | 2/5 | 2/5 |
| $X_g$ | M | 1 | 0 | 0 |

Provided that the forces driving allele-frequency change are relatively weak, and the process of mixing between states is relatively fast, then we can make the approximation that the allele frequency is the same in all classes (*p*_*i*_ *≈ p*). We can then describe the forces changing overall allele frequency by averaging across forces acting within each of the different classes, weighting them by the corresponding reproductive values (Nagylaki 1979; Laporte and Charlesworth 2002; Mullon et al. 2012; Lehmann 2014). For example, the (coalescent) effective population size is 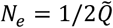, where

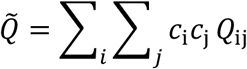

is the reproductive-value weighted rate of coalescence per generation, and *Q*_ij_ is the probability that a gene copy in state *i* and a gene copy in state *j* coalesce in the previous generation.

Similarly, we can calculate the effective mutation rate 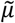 as

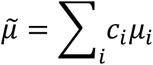

where *µ*_*i*_ is the mutation rate in class *i* (Lehmann 2014). And, finally, the effective marginal fitness effect is

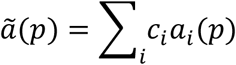

where *a*_*i*_ *(p)* is the marginal relative fitness effect of the allele in the context of class *i*, expressed as a function of the allele’s frequency, *p*. A fuller explanation of these effective population statistics is given in SM§1. Expressions for these weighted forms across different portions of the genome are given in Table 3.

**Table 3:**
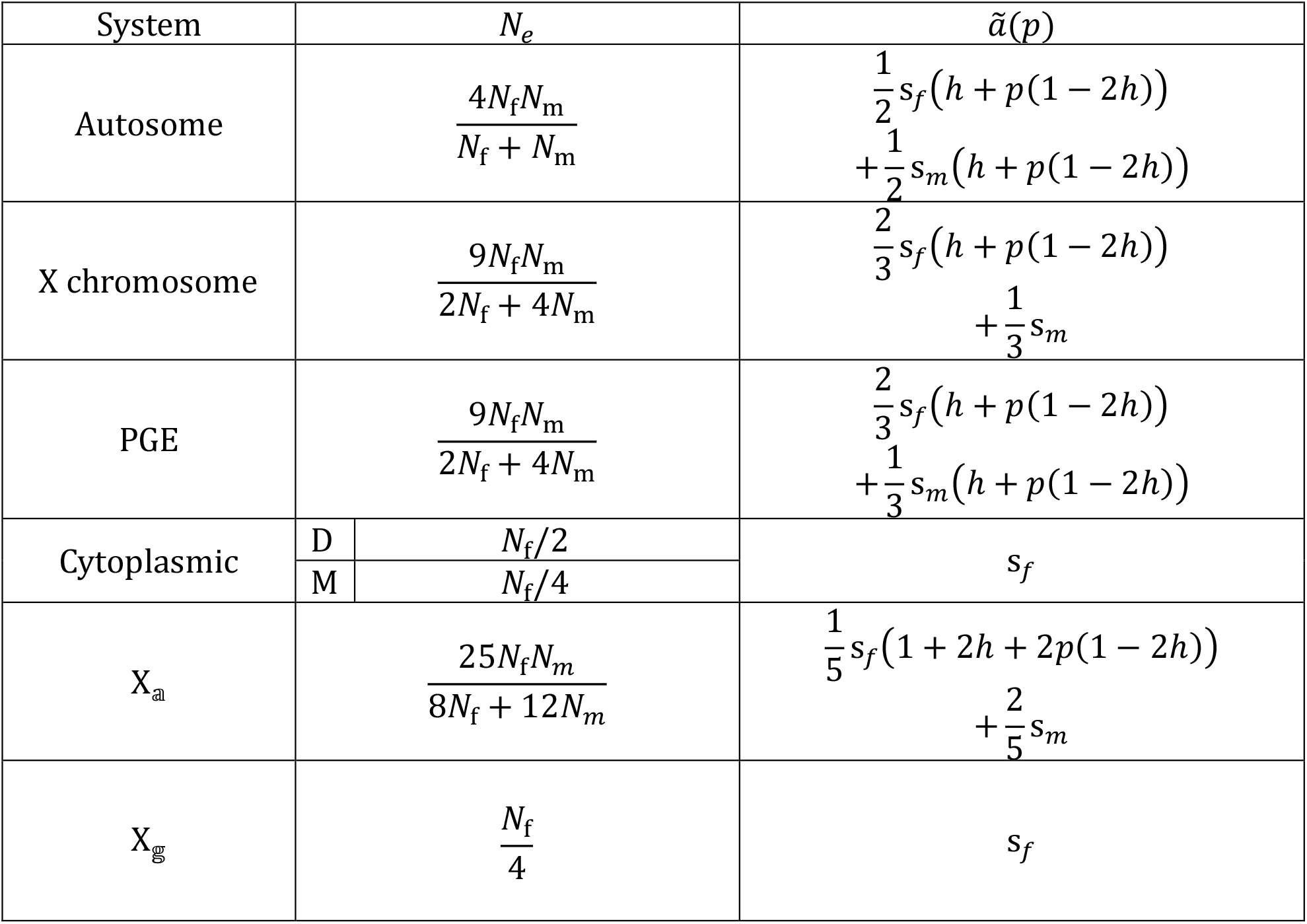
Effective population size and marginal fitness effects of different portions of the genome under different inheritance systems. We assume that there are an equal number of gynogenic and androgenic females (2*N*_f G_ = 2*N*_f *A*_ = *N*_f_), and a Poisson distributed number of offspring. For cytoplasmic elements strict matrilineal inheritance is assumed, we present results for both monogenic (M) and digenic (D) cases. Fuller expressions with unequal morph numbers and reproductive skew can be seen in SM§1.

These results enable us to identify some key patterns. The first is that both the effective population size, *N*_*e*_, and the mutation rate, 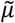, will be equal for the autosomes and X chromosomes under PGE, due to their identical transmission genetics. The second is that the marginal fitness effect, 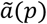, will typically be lower for autosomes and higher for X chromosomes, due to their different male somatic ploidy. Monogenic-linked portions of the genome will typically have a lower effective population size than either the X chromosomes or the autosomes. They will also experience relatively more efficient selection due to their effective haploidy in the gynogenic females. These latter comparisons are sensitive to sex-biases in occurring in demographic, mutational, and processes, as these portions of the genome spend a different fraction of time in males and females than the X chromosome and autosomes (Table 3, SM§1).

### Genetic diversity

We now consider processes that might maintain genetic diversity within populations, and how the balance of these forces may work differently for the different portions of the genome.

#### Neutral genetic diversity

For loci not under selection, the genetic diversity will be governed by a balance between mutation and drift. For such loci, under the infinite sites model, the pairwise genetic diversity may be written out as

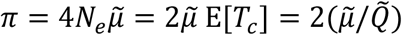

such that we can alternatively view this in terms of the effective population size, the expected time to coalescence, or the average coalescent rate (Hudson 1990). Based on the balance of these two forces, we can see that, under PGE, the autosomes and X chromosomes should harbour the same neutral genetic diversity (and distribution of variant frequencies), and that this will typically be lower than that for the autosomes in a comparable “eumendelian” system. For monogeny-associated portions of the X chromosome, X_𝕒_ or X_𝕘_, we expect a lower genetic diversity, due to its lower effective population size. Additionally, as the effective population size of the mitochondria is halved under monogenic – as opposed to digenic – reproduction, their genetic diversity is expected to be halved as well (SM§1-2), a result that applies to other species with split sex ratios (e.g. Meunier *et al*. 2008).

Sex-specific processes may alter the relative effective population sizes and mutation rates of different portions of the genome, and hence also the relative neutral genetic diversity. Under PGE, sex differences in mutation and demography will affect the autosomes and X chromosomes in the same manner, leaving their relative neutral genetic diversity unaffected (Fig1A-B). In contrast, monogenic associated portions of the genome will be affected by either sex differences, or differences between gynogenic and androgenic females. For example, whilst the X_𝕒_ will typically have a lower effective population size (and lower neutral diversity) than the autosomes, they will be close to equal with a sufficiently male biased sex ratio and can even exceed the autosomes if there are also more androgenic than gynogenic females (see SM§1-2).

**Figure 1:**
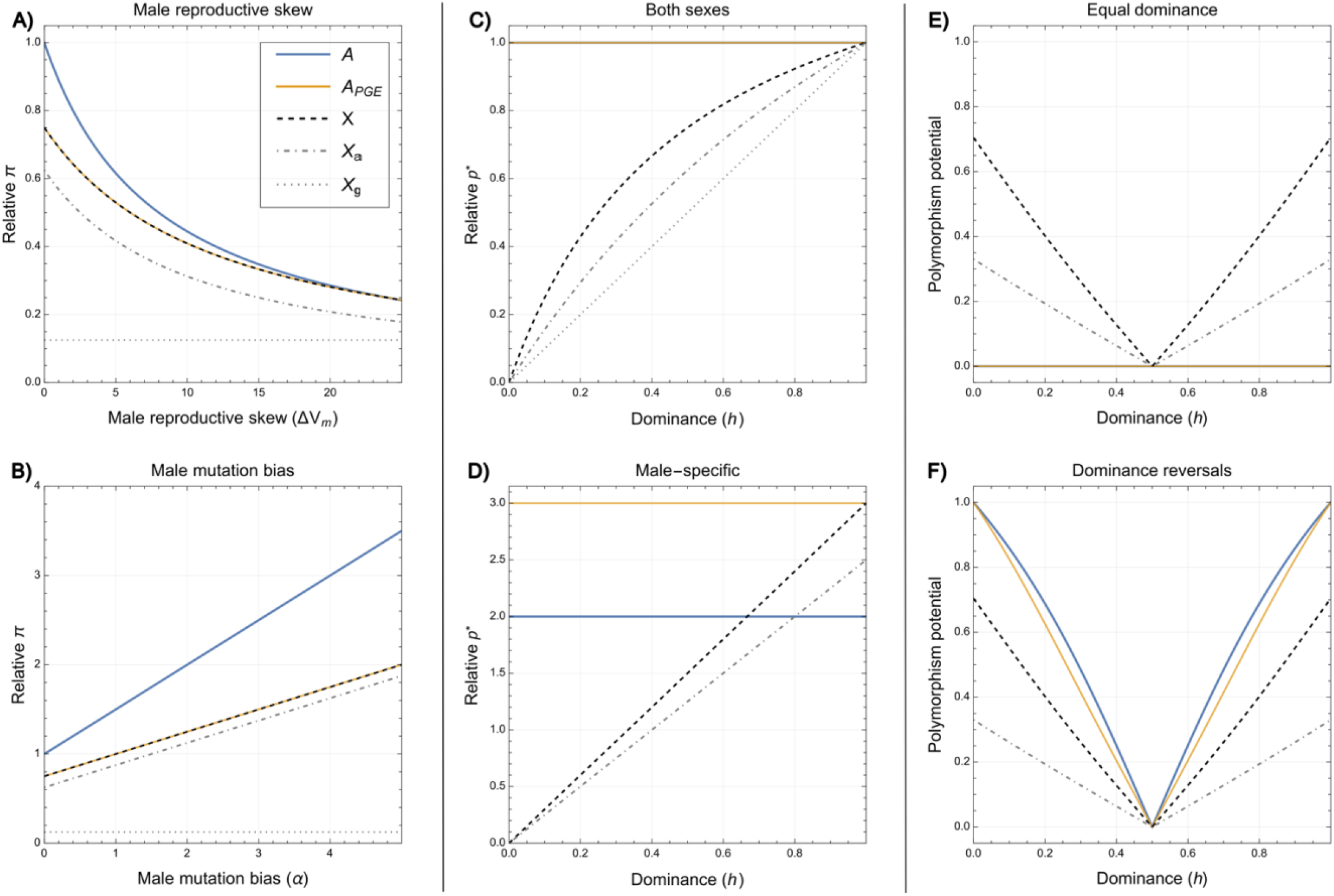
Patterns of relative genetic diversity differ under X/PGE systems as compared to standard X/autosome systems. Relative neutral genetic diversity (π_*i*_ /4*N*_*e*_*µ*) is always expected to be equivalent on PGE autosomes and X chromosomes, regardless of (A) sex-biases in demography, such as higher male reproductive skew, or (B) sex-differences in mutation rate. Δ*V*_*m*_ is excess over the Poisson variance in male offspring number (Laporte and Charlesworth 2002) and *α* is the degree of male mutation bias, where *µ*_*m*_ =(1+*α)µ*_f_ (Kirkpatrick and Hall 2004). (C) The relative frequency of a deleterious mutation (*p*_∗_ /*(µ*/*hs*)) is expected to be higher on a PGE autosome as compared to the X chromosome, regardless of the assumption about dominance. (D) This is especially true for loci only expressed in males. (E) The space for sexually antagonistic polymorphisms is smaller on PGE autosomes under equal dominance (*h*_f_ =*h*_*m*_), (F) but is larger under dominance reversals (*h*_f_ =1−*h*_*m*_). The polymorphism potential is calculated at the angle between the two invasion conditions, divided by π/2 radians – the maximum possible angle (SM§4).

#### Mutation selection balance

A second source of genetic diversity within populations is the balance between deleterious mutation and selection. Once again, if we consider that selection and mutation are relatively weak (relative to the rate of mixing), then we can consider the balance between these two forces in determining the equilibrium allele frequency (i.e. such that Δ*p*_*s*_ + Δ*p*_*µ*_ = 0). The greater the efficacy of selection, the lower will be the frequency of these deleterious alleles. We can approximate the equilibrium allele frequency as

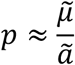

which, for an additive, deleterious mutation with fitness cost *s* in the hemi/homozygous state, gives *p ≈* 2*µ*/*s* for both eumendelian and PGE autosomes, and *p ≈* 3*µ*/2*s* for X chromosomes (Haldane 1935; Avery 1984; SM§2; SM§4). Consequently, under PGE, the autosomes are expected to harbour more deleterious mutations than the X chromosomes, due to the equivalent mutation rate, but smaller effective fitness effects (as deleterious mutations are partially “shielded” from selection in males). They also generate a sex-symmetric genetic load, unlike for X chromosomes (Werren 1993). In this regard, X/A comparisons under PGE will be identical to those in a classic X/A system. The androgenic and gynogenic-associated portions of the X will show different results. Their effective haploidy in the gynogenic females (as well as haploidy in the males), means that they are expected to harbour a lower frequency of deleterious alleles (Fig 1C-D). This also results in a lower genetic load although, like X chromosomes, this load is not equally imposed upon all classes (SM§4).

For genes that experience selection in only one sex, the frequency of deleterious mutations will be higher. This is because, by being sex-specific, they are less frequently exposed to selection. For both eumendelian autosomes and X chromosomes, sex-specific selection (under additivity) doubles the frequency of deleterious alleles (Avery 1984; Dapper et al. 2022), as in both cases the selection acting on the locus has effectively halved. For the PGE autosomes, as less of the ancestry is bound up in males, the male-specific loci are expected to harbour a much higher frequency of deleterious alleles compared to unbiased genes (3 times higher). Moreover, this also results in a much higher genetic load upon males under PGE than either typical autosomal or X-linked systems, and one that is sensitive to both dominance and the strength of selection (SM§4). Female-specific loci will also show an increased frequency of deleterious alleles compared to unbiased genes, but to a lesser extent (1.5 times higher). Thus, whether one is looking at constitutive expression, female-specific, or male-specific loci, one expects distinct results with X/A comparisons under PGE (Fig 1C-D). We can also extend this to consider the androgenic and gynogenic specific portions of the genome. Such loci will experience more haploid expression, and thus in general a lower frequency of deleterious mutations would be expected (Fig 1C). The allele frequency will be elevated with sex-specific gene expression, and even more so with female morph-specific gene expression (Fig 1D, SM§2, SM§4).

#### Balancing selection

As well as the balance between mutation versus drift, and mutation versus selection, selection itself may be capable of maintaining polymorphisms within populations. Perhaps the best investigated aspect of selectively maintained polymorphism has been the study of sexual antagonism (Owen 1953; Parsons 1961; Kidwell et al. 1977; Rice 1984; Connallon and Clark 2012; Mullon et al. 2012; Hitchcock and Gardner 2020; Klein et al. 2021; Flintham et al. 2023). If selection is weak, then we can make the approximations regarding the action of selection in the same way as before. Importantly, for a polymorphism to be stably maintained the following condition must be met:

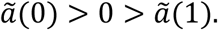

That is, the allele can invade from rarity (the first inequality holds) but cannot fix (the second inequality holds). If these conditions are satisfied, then an approximation for the polymorphic equilibrium point *p*^∗^ satisfies 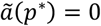. For sexually antagonistic alleles in PGE systems, we can identify two clear differences between the autosomes and X chromosome. The first is that – due to less-effective selection in males – female-beneficial alleles more readily invade and fix on the autosomes than the X chromosome, and this pattern holds for all dominance coefficients (Klein et al. 2021; Hitchcock et al. 2022). Secondly, the parameter space under which stable polymorphism obtains is very sensitive to assumptions about dominance. If there is equal dominance in the two sexes (*h*_f_ = *h*_*m*_), then the X chromosome has a much larger scope for stable polymorphisms. In contrast, if there are dominance reversals (*h*_f_ = 1 − *h*_*m*_), then the autosomes have much greater scope for stable polymorphisms. This is qualitatively similar to standard X/A systems (Fry 2009; Patten 2019). The key difference, however, is that the polymorphisms that will be maintained under PGE are typically more male-biased. One explanation is that the space for female fixation has expanded, and the space for male fixation has retreated, and so the polymorphic space is enriched for male-beneficial alleles.

Alongside sexual antagonism are the possibilities for other forms of trade-off that may maintain polymorphism. One novel example that may be found in monogenic species is the trade-off between gynogenic and androgenic females, which we will refer to here as maternal antagonism. This can be thought of as a subset of parental antagonism, i.e. where parents exhibit trade-offs between the production of sons and daughters (Miller et al. 2006; Rice et al. 2008; Patten and Haig 2009; Friberg and Rice 2015; Hitchcock et al. 2022). For maternal antagonism on the androgenic portion of the X chromosome, the evolutionary dynamics are very similar to sexual antagonism on an X chromosome. Gene copies here are expressed more frequently in androgenic females but will be haploid when expressed in gynogenic females. Thus, whether we anticipate androgenic-beneficial or gynogenic-beneficial alleles to accumulate on this portion of the genome will be sensitive to assumptions about dosage and dominance, similarly to the invasion of female and male-beneficial alleles on X chromosomes (Patten 2019; Frank and Patten 2020; Hitchcock and Gardner 2020). A very similar dynamic will also occur for the autosomally linked monogenic regions (SM§2, SM§4).

### Genetic divergence

Fixation of distinct alleles causes populations to diverge genetically. First, we analyse the relative substitution rates of beneficial mutations – for both single copies and standing genetic variation – and then those of neutral and deleterious alleles.

#### Adaptive substitutions: single copies

For new, beneficial mutations, we may use a branching-process approximation to determine the fixation probability (Haldane 1927; Patwa and Wahl 2008). When there is class structure, as here, then one must consider a multitype-branching process (Allen 2011). Assuming that the growth factor of the mutant lineage on average is close to 1 (i.e. selection is weak) then the probability that a mutation which arises in class *i* fixes in the population can be written as:

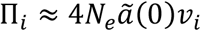

Where ν_*i*_ is the reproductive value of a single gene copy in class *i*, normalised such that the total reproductive value sums to 1. This represents a generalisation of Haldane’s approximation ∏ *≈* 2 *s* for haploid genetics (Haldane 1927; Hoppe 1992; Pollak 1992; Pollak 2000). A fuller derivation, and numerical results investigating the consequences of relaxing this assumption, are provided in SM§3. Inserting the values from Table 3, we can see that the probability of fixation is larger for mutant alleles that spend a greater fraction of their time in haploid setting (i.e. the basis for the faster X effect).

The rate of adaptive substitution *K*can be expressed as a product of the flux of new mutations, and their probability of fixation (Kimura and Ohta 1971a; Kimura and Ohta 1971b; Charlesworth et al. 1987; McCandlish and Stoltzfus 2014). If the number new mutations entering a population is given by the physical number of gene copies and their respective mutation rates, then the adaptive substitution rate will be

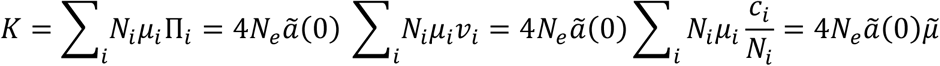

where 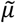 is the class reproductive value weighted mutation rate described earlier. Assuming no sex-biases in the mutation rate, then for a standard X chromosome system (with full dosage compensation), the classic result given by Charlesworth et al. (1987) is

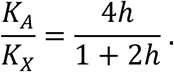

In contrast, for an organism with PGE we have

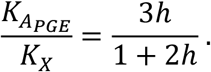

Thus, regardless of the degree of dominance, we still expect a faster rate of accumulation of beneficial mutations on the X chromosome than on the autosomes in a system with PGE (Figure 2A). For the PGE case, this holds even if there are sex-differences in the mutation rate, as – unlike in classic X autosome comparisons – the two portions of the genome will experience the same fraction of time in males and females (Kirkpatrick and Hall 2004). We can further partition the adaptive substitution rate according to the relative fitness effects experienced in the two sexes, and thus the rate of adaptive substitution for sex-specific genes. As the differences between the autosomes and X chromosome are driven purely by males, then we expect no differences at female-specific loci, and more-exaggerated differences at male-specific loci (Figure 2B).

**Figure 2:**
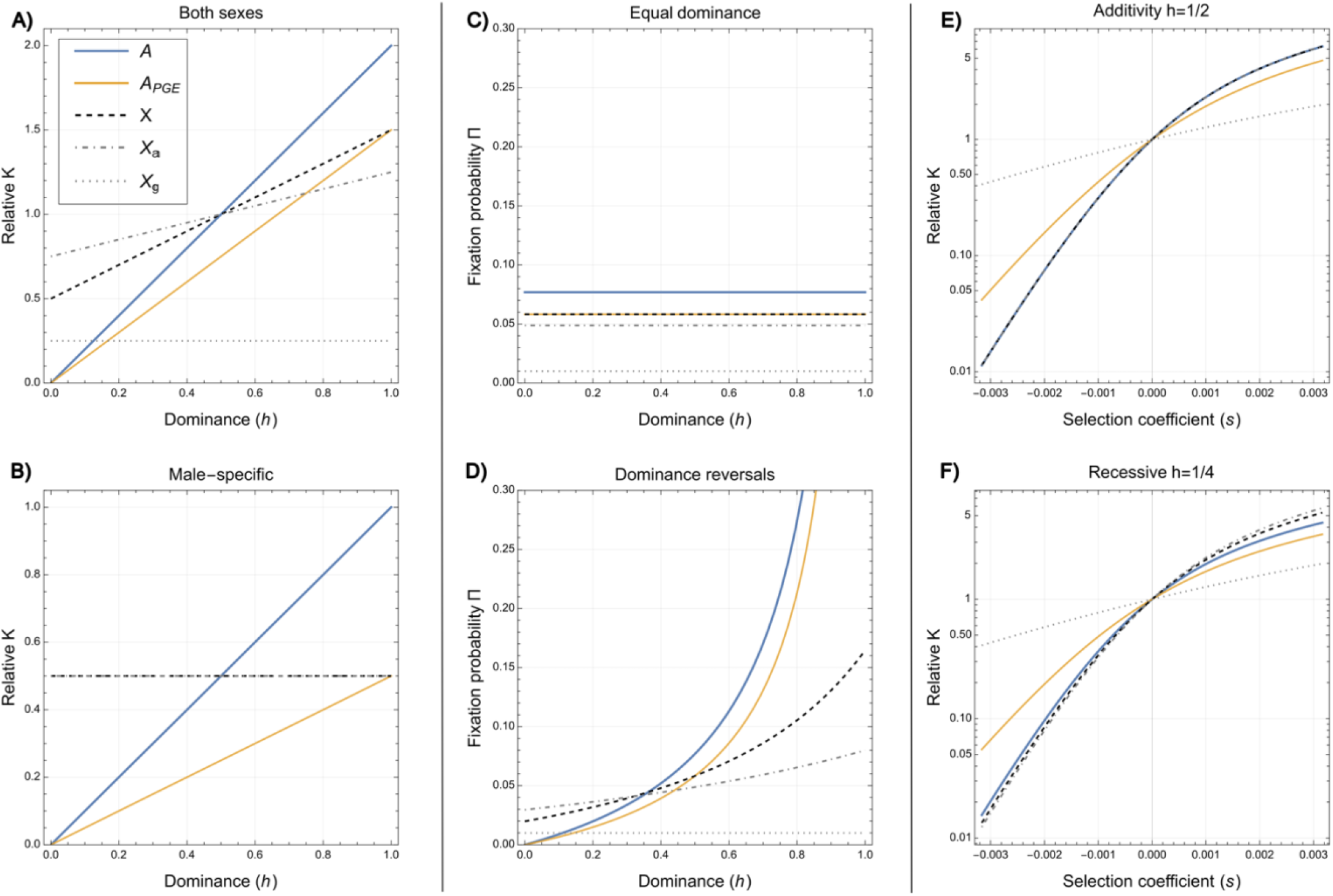
Rates of substitution differ under X/PGE systems as compared to standard X/autosome systems. (A) Relative rates of adaptive substitution (*K*_*i*_ /[2*Nµs*]) are always expected to be lower on PGE autosomes than X chromosomes, and (B) this pattern is strongest for recessive male-specific mutations. (C) Adaptive mutations arising from previously standing genetic variation are equally likely to fix for PGE autosomes and X chromosomes when the dominance during the deleterious and beneficial phase are assumed to be equal (*h*_*b*_ =*h*_*d*_). (D) When there are reversals of dominance (*h*_*b*_ =1−*h*_*d*_), then dominance beneficial mutations have a higher fixation probability under PGE autosomes than X chromosomes, but a lower fixation probability when recessive. For (C) and (D) *N*=10^4^, *µ*=10^−5^, *s*_*d*_ =0.2, and *s*_*d*_ =0.01. (E) For weakly deleterious mutations, the PGE autosomes have a higher relative substitution rate (*K*_*i*_ /*µ*) than X chromosomes. For weakly beneficial mutations, the PGE autosomes have a lower relative substitution rate than X chromosomes. (F) These effects are stronger for recessive mutations. In both (E) and (F) *N*=10^3^, and an equal sex and morph ratio are assumed.

We can similarly consider the androgenic-specific portions of the sciarid chromosome, to obtain

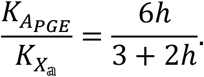

That is, compared with the autosomes in the PGE system, we also expect a faster rate of adaptation for the *X*_𝕒_ region, because not only is this portion of the genome haploid in males (unlike the autosomes), but it is also haploid/pseudohaploid in gynogenic females. A similar pattern will apply to the androgenic portion of the gall midge supergene, although less extreme due to its diploidy in males (SM§3).

#### Adaptive substitutions: standing genetic variation

In addition to drawing upon new mutations, adaptive substitutions may also be fuelled by standing genetic variation. Using the approach of Orr and Betancourt (2001), and assuming that a previously deleterious mutation with marginal fitness effect 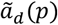 held at mutation-selection balance becomes positively selected with fitness effect 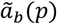, then we can again use a branching-process approximation. However, rather than starting with a single copy of the allele, we now begin with *k* copies in the population. We also assume that the reproductive value of a single gene copy is equivalent in all classes (ν_*i*_ = 1/*N*_*h*_). Considering *k* independent branching processes, and calculating the probability that at least one does not become extinct, then the fixation probability can be expressed as

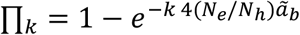

where for brevity we will write 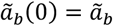 and 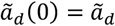. If, following Orr and Betancourt (2001), we assume that the initial number of copies of the allele is given by the mutation-selection balance expressions 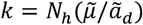, then the fixation probability is

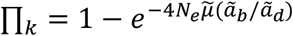

which, if selection against the deleterious mutation is stronger than selection for the beneficial mutation 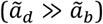 approximates to

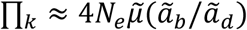

Consider the contrast, under PGE, between an autosomal and X-linked locus. In both cases, the effective population size will be equivalent. However, the marginal fitness effects will not be. If dominance is assumed to be the same between the deleterious and beneficial phase, then this ratio between the marginal fitness effects – and hence the probability of fixation - will be the same for the autosomes and X chromosomes.

The equal-dominance assumption may be questioned. Orr and Betancourt (2001) initially argued that if dominance emerges from a physiological source (e.g. non-linearities of enzyme function (Kacser and Burns 1981)), then there is little reason to suppose it might differ between the deleterious and beneficial phase. Recent work has instead argued that reversals of dominance might be common if dominance emerges from non-linearities in fitness landscapes (Fry 2009; Manna et al. 2011; Connallon and Chenoweth 2019; Muralidhar and Veller 2022). Exploring this latter suggestion, if the allele is recessive when deleterious, but dominant when beneficial, then this results in a higher fixation probability on the autosomes than on the X chromosome (Fig 2D). In contrast, if the allele is dominant when deleterious and recessive when beneficial, there will be a higher fixation probability on the X chromosome than on the autosomes. Thus, in the case of adaptation from standing genetic variation – unlike the previously analysed new mutation scenario – the relative rate of adaptation is qualitatively sensitive to assumptions about dominance (Fig 2C-D). A richer understanding of expected distributions of dominance for certain genes, pathways, and selective regimes is important for generating more detailed predictions about these patterns of adaptation (Billiard et al. 2021). Similar qualitative conclusions are reached by instead using the stationary distribution of allele frequencies, rather than the deterministic mutation selection balance (Orr and Betancourt 2001; Hermisson and Pennings 2005; Hermisson and Pennings 2017; Charlesworth et al. 2018).

#### Neutral and weakly deleterious mutations

The above results focus on adaptive substitutions. However, neutral and deleterious alleles may also become fixed in finite populations as a consequence of random drift. To analyse non-adaptive substitutions, we instead adopt a diffusion approximation, whereby we consider a mutant allele that is initially present at frequency *p*_0_ in the population (Kimura 1962; Avery 1984; Charlesworth et al. 1987; Vicoso and Charlesworth 2009; Mrnjavac et al. 2023). We can then write out the probability that this new mutation fixes as:

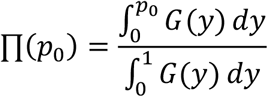

where

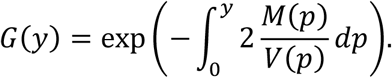

Here, *M(p)* is the deterministic advection term which we can further decompose into the average effect 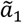, and the second order terms due to the deviation from additivity 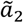 (Mrnjavac et al. 2023), to obtain

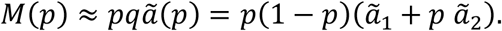

The diffusion term is given by:

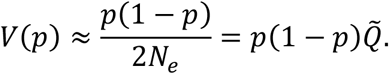

Accordingly, we can express the fixation probability in terms of error functions (see SM§2). Focusing on the additive case 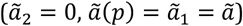, the fixation probability is given by:

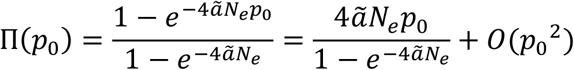

which can be further simplified by assuming that the initial allele frequency is small (*p*_0_ ≪ 1) - typically giving a very good approximation (Vicoso and Charlesworth 2009; Mrnjavac et al. 2023). In the neutral limit, the fixation probability is simply the allele’s starting frequency 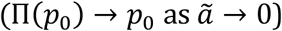. Under PGE, new mutations on both autosomes and X chromosomes will start at the same (reproductive value-weighted) allele frequencies, and thus we would expect the neutral substitution rate to be equivalent for these portions of the genome, given by the effective mutation rate, 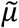 (Lehmann 2014). This will still obtain even if there are sex-differences in the number of breeders and/or sex-differences in the mutation rate, unlike in eumendelian systems (Kirkpatrick and Hall 2004; Vicoso and Charlesworth 2009).

Secondly, we can see that for deleterious variants, the fixation probability will be substantially higher on the autosomes than on the X chromosome. This is especially true for those variants that are recessive and male-specific (Figure 2E-F). Thus, we may expect that the autosomal loci under PGE should accumulate deleterious mutations at a faster rate than the X chromosome. Whilst the quantitative extent of this difference is sensitive to the dominance coefficient and sex-differences in expression, the qualitative pattern is unaffected by these factors. This is not true of eumendelian systems, wherein – in the standard additive, constitutively expressed case – the faster accumulation of deleterious mutations on X chromosomes is a pattern that is sensitive to assumptions not only about the expression of such loci but also about sex-differences in demography (Vicoso and Charlesworth 2009).

Finally, returning to beneficial mutations, we can see that if *p*_0_ = 1/*N*_*H*_ (i.e. the allele is initially present in a single copy) and 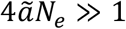, then the fixation probability is given by

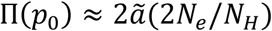

which recovers the branching-process approximation when the number of offspring is Poisson distributed (*N*_*e*_ = *N*_*H*_ /2).

## Discussion

Here we have developed a series of population genetic models tailored to the atypical genetics of fungus gnats, gall midges, and globular springtails, focusing on the effects of two features: paternal genome elimination and monogenic reproduction. We found that, under paternal genome elimination, the autosomes and sex chromosomes are likely to show distinct patterns of genetic diversity and divergence to those of classic systems, with neutral processes being identical between the autosomes and sex-chromosomes, whilst selective processes are more effective on X chromosomes. Additionally, we found that in species with monogenic reproduction, monogeny associated portions of the X (or A) show a distinct inheritance pattern, with typically lower effective population sizes, but more effective selection, and therefore different predictions about the diversity and divergence in these genomic regions. Moreover, under monogeny, the effective population size of mitochondria is halved, leading to reduced genetic diversity and efficacy of selection on this section of the genome.

Our results point to species that combine PGE and X chromosomes as providing excellent opportunities for investigating the faster X effect. The equal effective population size of the X chromosome and the autosomes – independent of sex-biases in demography – means that these genetic systems isolate the effects of haploid vs diploid selection in males. However, this picture may be complicated by the sources of adaptive variation. Whilst classic work has modelled the process of adaptive substitution as drawing upon new mutations (Charlesworth et al. 1987), more recent models consider that it may be fueled by standing genetic variation (Orr and Betancourt 2001; Hermisson and Pennings 2005; Muralidhar and Veller 2022). The latter situation may result in a higher initial frequency of beneficial mutations on the autosomes and, under some scenarios, may instead generate a faster autosomal effect. Charlesworth and colleagues (2018) – making use of parameter estimates from *Drosophila* – suggested that the rate of adaptive substitution will likely be dominated by the contribution from segregating variants. Additionally, it has also been suggested that in standard X/autosome comparisons we should expect proportionally more hard sweeps on the X chromosome (Charlesworth et al. 2018), with some recent evidence for this pattern in *Drosophila* (Harris and Garud 2023; Harris et al. 2024). Putting these together, whilst the overall rate of adaptative substitution on X vs A will vary depend upon patterns of dominance, we nonetheless would expect a greater fraction of sweeps on the X to be “hard”. Although in practice it may be challenging to disentangle these scenarios as even if there are initially multiple copies of a beneficial mutation in many scenarios only one copy will go to fixation, eliminating the signature of the sweep (Orr and Betancourt 2001; Jensen 2014; Harris et al. 2018).

A second complication in testing the faster-X effect concerns neutral and deleterious variants. In conventional systems, under certain demographic regimes, these non-beneficial alleles may fix more readily on X chromosomes, also driving a faster-X effect. For example, this is likely the main force driving the observed faster X effect in aphids (Jaquie ry et al. 2018). Under PGE, neutral or nearly neutral variants should behave almost identically on autosomes and X chromosomes, and thus neutral regions – such as pseudogenes – should evolve at a similar rate whilst deleterious variants have a higher fixation probability on the autosomes than on the X chromosome. Methods based on the McDonald-Kreitman test have been developed that allow beneficial and deleterious variants to be discerned (McDonald and Kreitman 1991; Eyre-Walker 2006; Stoletzki and Eyre-Walker 2011). A better understanding of the distribution of fitness effects, and estimations of the relative fraction of fixations that are beneficial vs neutral vs deleterious variants is crucial for interpreting substitution patterns in these groups.

Alongside patterns of substitutions, PGE species are expected to show distinct patterns of within-population variation to conventional systems. Specifically, we expect equivalent neutral variation across the X chromosome and autosomes in these systems, regardless of demography. Such patterns may be assessed by surveying regions of the genome that are assumed to be neutral, e.g. fourfold degenerate sites. Complications to this picture may emerge when there are interactions between neutral loci and those under selection. For example, selection both against strongly deleterious alleles (background selection) and for beneficial alleles (selective sweeps) may reduce genetic variation at linked sites (Charlesworth and Jensen 2021). Given the increased efficacy of selection on the X chromosome, then we may also expect these effects to be strongest on this portion of the genome, and thus reduced genetic variation on the X chromosome. More specifically, this pattern is expected to be strongest in low-recombination regions of the genome, such as those with inversions associated with monogeny. In addition, recent work has shown scenarios where weakly beneficial variants may actually increase the genetic variation of linked sites (Zhao and Charlesworth 2016; Charlesworth 2022). Given the generally weaker selection on the autosomes, then this may provide a further mechanism generating higher levels of neutral variation on the autosomes in these groups.

The present analysis has assumed that alleles expressed in hemizygous and homozygous males behave equivalently. This assumption is a common feature of models of X/autosome comparisons and is typically justified by the presence of dosage compensation mechanisms, which normalize the relative expression despite the reduced physical number of gene copies. However, there are many documented instances in which this is not the case (Furman et al. 2020). If at an X-linked locus there is a lack of dosage compensation, then we would expect it to behave more similarly to the autosomal case (Charlesworth et al. 1987; Mank et al. 2010). A similar effect will occur under PGE if gene expression is not equivalent between the maternal-origin and paternal-origin genomes. In PGE species, there is documented variation in the balance of expression due to silencing or elimination of the paternal genome (Herbette and Ross 2023). With an increasing fraction of expression emerging from the maternal copy, then the X/autosome differences are expected to disappear. Another further reason for breakdown in the hemizygous/homozygous equivalence is the presence of interacting gene copies (Mrnjavac et al. 2023). Whilst this is not likely to be applicable to the X chromosomes of these species (as there is no Y chromosome), it will apply to interactions between the gynogenic and androgenic associated portions of the X chromosome (Box 2., SM§1). Selection acting solely during the haploid phase would also eliminate differences between the X chromosome and the autosomes, as it does in eumendelian systems (Immler et al. 2012).

We have here focused predominantly on the genetic aspects of these species, assuming panmixia, constant population size, and no population structure. Yet departures from these assumptions are likely to be important aspects of these groups’ ecology (Yukawa and Tokuda 2021). For example, in gnats and midges, only adults disperse and are typically both short-lived and weak fliers, likely leading to limited dispersal and population structure (Briggs and Latto 2000; Cloyd 2015). Although there is great variation across species (Sutou et al. 2011; Yukawa et al. 2019). This may have some important effects on the results outlined here. Firstly, increased inbredness (*sensu* Frank 1986, *F*_IS_) will reduce differences in the efficacy of selection between the sex chromosomes and autosomes (Hitchcock and Gardner 2020; Flintham et al. 2021; Hitchcock et al. 2022). In the limit of full inbredness, the differences we have outlined between the X chromosome and the autosomes will be eliminated. Secondly, the mating structure may generate systemic sex-differences in the magnitude of selection. For example, with local mate competition, males are disproportionately likely to compete with related males, but this may not occur (or will do to a lesser extent) for females (Hamilton 1967). For example, in the lentil gall midge, adults tend to mate close to their site of emergence, before gravid females disperse to new fields (Kolesik 1993). In such cases, this will effectively weaken selection on males, and thus reduce the magnitude of the faster-X effect (Flintham et al. 2021; Hitchcock et al. 2022). More generally, population structure and fluctuations in population size will alter fixation probabilities, patterns of genetic variation, and the balance between hard and soft sweeps, and complicate signatures of selection in the genome (Soni et al. 2023; Sudbrack and Mullon 2024 Mar 25). A better empirical understanding of the ecology of these groups, and theoretical models of these scenarios, should prove a fruitful area for future investigation.

In conclusion, our results suggest that X/PGE systems can provide exceptional opportunities for testing evolutionary theory. And whilst some these groups have long been the focus of genetic study (Metz 1925; Gerbi 2022), only with new developments in sequencing technology have we started to acquire the whole genome data with which we can put such theory to empirical test (Zhao et al. 2015; Urban et al. 2021; Jaron et al. 2022; Baird, Urban, et al. 2023). Moreover, the diversity and species richness of these groups – spanning the arctic to the tropics and pests to pollinators - may also make them very suitable for comparative analyses (Menzel et al. 2020). This fusion of genetics, natural history, and evolutionary theory offer new avenues for testing old hypotheses and enriching our understanding of evolution in these groups and across the tree of life.

### Box 1.

Paternal Genome Elimination

is a genetic system found in groups of flies (fungus gnats, gall midges), springtails, mites, coccids, lice, and beetles (Herbette and Ross 2023). Under this system both males and females arise from fertilized eggs, and thus receive genes from both fathers and mothers. However, during their life, males eliminate their paternal genome, such that they only transmit their maternal-origin genes, and thus their transmission genetics are functionally haplodiploid. Both the timing of the elimination and the degree of paternal expression vary greatly, and so the “effective ploidy” of the males depends on the species, tissue, and genes of interest. Additionally, in some groups, such as the fungus gnats, gall midges, and springtails, post-fertilization elimination of the paternal X chromosome in males is also the mechanism of sex determination. Whilst embryos initially contain both maternal and paternal-origin X chromosomes (X_*m*_, X_p_), a round of X chromosome elimination occurs such that males only retain their maternal X chromosomes, generating X0 males and XX females.

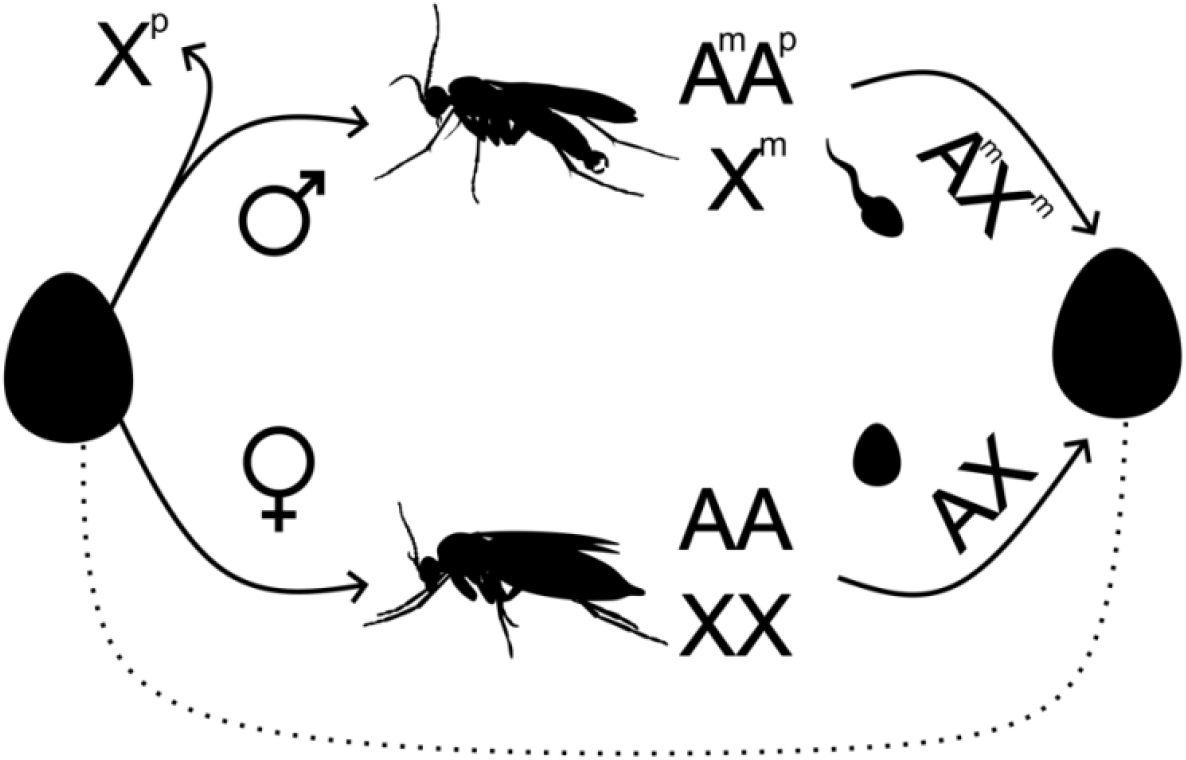

### Box 2.

Monogenic reproduction

is a sex ratio strategy found in a range of species including flies, wasps, and bees (Stone et al. 2002; Meunier et al. 2008; Baird, Mongue, et al. 2023). In *Bradysia coprophila*, the model fungus gnat, this process is governed by a novel form of the X chromosome - the X’. Females who are heterozygous for this chromosome (X’X) are gynogenic (producing only daughters) whilst those that do not possess the X’ (XX) are androgenic (producing only sons). Males contain a single copy of the X. The X’ and the X are distinguished by an inversion present on the X’ (Crouse 1979; Baird, Urban, et al. 2023). This portion of the genome, X_𝕘_, is exclusively restricted to gynogenic females. Due to this restriction, and its lack of recombination, the inversion has a much-reduced effective population size, and its gene content has partially degenerated. In contrast, the non-inverted portion, X_𝕒_, spends time in both types of female, as well as males. This scenario is illustrated below. In gall midges, a similar system appears to exist, but with the inversion located on an autosome (Benatti et al. 2010). Thus, whist the transmission genetics are the same, the gall midge system is distinguished by the males being somatically diploid in the _𝕒_ region. Finally, blowflies, a more distantly related group which do not possess PGE, also have monogeny controlled in a similar fashion by an autosomal locus (Ullerich 1996). Thus, whilst this is conceptually similar, due to the lack of PGE, the transmission genetics are quantitatively distinct (SM§1). A fuller, recent review can be found in Baird et al. (2023b).

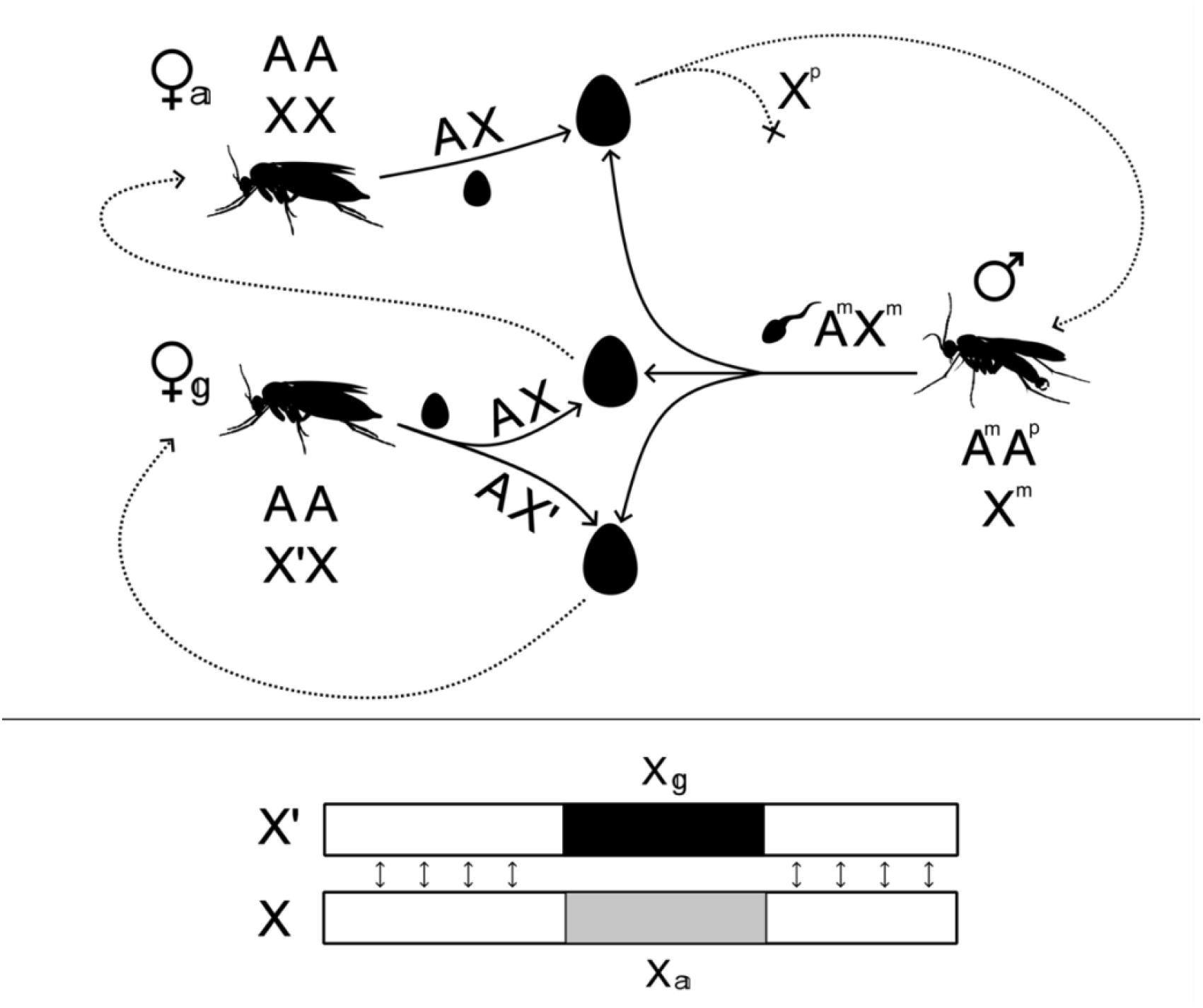

