## Supplementary Material for "Paternal genome elimination, monogenic reproduction, and the evolutionary genetics of atypical sex chromosome systems"

#### Summary

Here we outline further methods and results. In Section 1, we outline the modeled genetic systems and calculate the class reproductive values for different portions of the genome. We then use these reproductive values to calculate the effective population size, marginal fitness effects, and mutation rate. In Section 2, we then use diffusion equations to approximate the fixation probabilities and stationary distributions of allele frequencies. In Section 3, we use a branching process approximation to calculate the fixation probability for a beneficial mutation. Finally, in Section 4, we consider some deterministic results for the invasion of a sexually antagonistic or maternally antagonistic allele, and the region of polymorphism, and also the mutation-selection balance of a deleterious allele, and the genetic load that the deleterious allele imposes.

#### 1 Marginal fitness effects, mutation rates, and effective population size

##### 1.1 Genetic systems and class reproductive values

Here we consider results for eumendelian autosomes ($A$), paternal genome elimination ($PGE$), X chromosomes ($X$), and mitochondria ($M$), under monogeny ($\mathbb{m}$) and digeny ($\mathbb{d}$). Additionally, we consider the transmission genetics of monogeny associated supergenes. We consider three possible supergene systems corresponding to those known (or supposed) in fungus gnats, gall midges, and blow-flies. In all of these systems the daughter-producing female (gynogenic) is thought to be heterozygous, whilst the son-producing female is thought to be homozygous, males are also homo/hemizygous. We notate the gynogenic specific portion $\mathbb{g}$ and the other (androgenic associated) portion $\mathbb{a}$.

To describe these different genetic systems we consider three possible states that a gene copy may occupy: gynogenic females $f_{\mathbb{g}}$, androgenic females $f_{\mathbb{a}}$, and males $m$. We then describe the ways that genes flow between these states using the following transition matrix $T$, where the probability that a gene in state $i$ descended from a gene copy in state $j$ in the previous time-step is $t_{\mathrm{ij}}$, and so $T = \left(t_{\mathrm{ij}}\right)_{i,j\in G}$, where $G$ is the set of possible gene positions or classes. In our case of interest we can write out:

$$T = \begin{pmatrix} t_{f_{\mathbb{g}}f_{\mathbb{g}}} & t_{f_{\mathbb{g}}f_{\mathbb{a}}} & t_{f_{\mathbb{g}}m} \\ t_{f_{\mathbb{a}}f_{\mathbb{g}}} & t_{f_{\mathbb{a}}f_{\mathbb{a}}} & t_{f_{\mathbb{a}}m} \\ t_{mf_{\mathbb{g}}} & t_{mf_{\mathbb{a}}} & t_{mm} \end{pmatrix} = \begin{pmatrix} \left(\frac{1+\gamma}{2}\right)\alpha_{\mathbb{g}} & \left(\frac{1-\gamma}{2}\right)\alpha_{\mathbb{g}} & (1-\alpha_{\mathbb{g}}) \\ \left(\frac{1+\gamma}{2}\right)\alpha_{\mathbb{a}} & \left(\frac{1-\gamma}{2}\right)\alpha_{\mathbb{a}} & (1-\alpha_{\mathbb{a}}) \\ \left(\frac{1-\gamma}{2}\right)(1-\beta) & \left(\frac{1+\gamma}{2}\right)(1-\beta) & \beta \end{pmatrix} \tag{S1}$$

Where $\alpha_{\mathbb{g}}$ is the probability a gene copy residing in a gynogenic female descended from a female in the previous generation, $\alpha_{\mathbb{a}}$ is the probability a gene copy in an androgenic female descended from a female in the previous generation, $\beta$ is the probability a gene copy in a male descended from a male, and $\gamma$ scales the degree of monogenic reproduction, such that $\gamma = 0$ is digenic reproduction and $\gamma = 1$ being pure monogenic reproduction.

Class reproductive values describe the asymptotic fraction of the ancestry of the population that descends from a particular class today (Taylor, 1990; Grafen, 2006). The reproductive values can be calculated by taking the dominant left-eigenvector of our matrix $T$.

$$\vec{v} = \vec{v}T \tag{S2}$$

We can normalise these reproductive values by multiplying by the right eigenvector:

$$\vec{u} = T\vec{u} \tag{S3}$$

Such that:

$$\vec{c} = \frac{\vec{v}\times\vec{u}}{\vec{v}\cdot\vec{u}} \tag{S4}$$

We can also use this information, alongside $T$ to get the matrix of elasticities $E$. This provides the description of the reproductive value of particular types of transition through our life cycle, where $E = \left(e_{\mathrm{ij}}\right)_{i,j\in G}$, is the portion of time that a gene spends passing backwards in time from $i$ to $j$, or alternatively the time spend passing from $j$ to $i$:

$$e_{\mathrm{ij}} = t_{\mathrm{ij}}\frac{v_{\mathrm{i}}\times u_{\mathrm{i}}}{\vec{v}\cdot\vec{u}} \tag{S5}$$

Each element in $e$, describes the value of the flow of reproductive value between class $i$ and class $j$. And where:

$$\sum_j e_{\mathrm{ij}} = c_{\mathrm{i}} = \sum_j e_{\mathrm{ji}} \tag{S6}$$

Which is the row-column symmetry noted previously in demographic studies (Van Groenendael et al., 1994; Giaimo and Traulsen, 2021). Here we can see that one interpretation of this result is that because elasticities are equivalent to class reproductive values, and at equilibrium the reproductive value into a class must equal the reproductive value leaving that class, then the sum of the $i$th row and column must be equal. The reproductive values for our different genetic systems of interest can be seen in Table 1.

Table S1: Class reproductive values for different portions of the genome and under different inheritance systems, where $\lambda$ is the probability that a cytoplasmic gene copy descends from the mother, and $\gamma$ scales the degree of monogenic reproduction.

| | Females: gynogenic | Females: androgenic | Males |
|---|---|---|---|
| Autosomes | $\frac{1}{4}$ | $\frac{1}{4}$ | $\frac{1}{2}$ |
| PGE/$X$ | $\frac{1}{3}$ | $\frac{1}{3}$ | $\frac{1}{3}$ |
| Mitochondria | $\frac{1}{2}(1-\lambda)(1+\gamma(1-2\lambda))$ | $\frac{1}{2}(1-\lambda)(1-\gamma(1-2\lambda))$ | $\lambda$ |
| $A_{\mathbb{a}}$ | $\frac{1}{7}$ | $\frac{2}{7}$ | $\frac{4}{7}$ |
| PGE$_{\mathbb{a}}$/$X_{\mathbb{a}}$ | $\frac{1}{5}$ | $\frac{2}{5}$ | $\frac{2}{5}$ |
| $A_{\mathbb{g}}$/PGE$_{\mathbb{g}}$/$X_{\mathbb{g}}$ | 1 | 0 | 0 |

#### 1.2 Marginal fitness effects

First, let us consider the fitness scheme outlined in Table 1, where we consider two alleles {0,1}. The fitness differences between these two alleles are assumed to be small, such that our selection coefficients are of $O(\delta)$, and where we will ignore terms of $O(\delta^2)$. In that case, we can make the approximation that the allele frequency $p$ is equivalent in all classes $p_i \approx \tilde{p}$. We can then write the marginal fitness effect in this case as so:

$$\tilde{a} = \sum_i c_i a_i \tag{S7}$$

Where:

$$a_i = \omega_{i,1} - \omega_{i,0} \tag{S8}$$

Where $\omega_{i,1}$ and $\omega_{i,0}$ are the average relative fitness of the alleles in state $i$, and hence $a_i$ is the marginal fitness effect of our mutant allele (1). For a haploid state:

$$a_{hap} = \frac{1}{\bar{w}}\left(p(w_{11} - w_{01}) + (1-p)(w_{01} - w_{00})\right) \tag{S9}$$

And for a diploid state:

$$a_{dip} = \frac{1}{\bar{w}}\left(p(w_{11} - w_{01}) + (1-p)(w_{01} - w_{00})\right) \tag{S10}$$

We may further partition the difference between these fitness effects (Fisher's average excess) to consider the additive and non-additive contributions to this. Alternatively, similar to Mullon et al. (2012), we can write as a scaling of strength of selection $\alpha$ and the difference between the allele frequency and the equilibrium allele frequency $p^* - p$.

$$a(p) = a_1 + pa_2 = \alpha\left(p^* - p\right) \tag{S11a}$$

$$\alpha = -a_2 \tag{S11b}$$

$$p^* = -a_1/a_2 \tag{S11c}$$

If the condition for the 1 allele to invade is:

$$I_1 = \lim_{p\to 0} \tilde{a}(p) = a_1 > 0 \tag{S12}$$

Table S2: Fitness scheme used for the population genetic analysis. $\eta$ scales the fitness effect experienced in the hemizygous haploid state as compared to the homozygous diploid state, and $h$ is the dominance coefficient.

| | 00 | 01/10 | 11 | 0 | 1 |
|---|---|---|---|---|---|
| $w_{f_{\mathrm{g}}}$ | 1 | $1+h_{f_{\mathrm{g}}}s_{f_{\mathrm{g}}}$ | $1+s_{f_{\mathrm{g}}}$ | 1 | $1+\eta_{f_{\mathrm{g}}}s_{f_{\mathrm{g}}}$ |
| $w_{f_{\mathrm{a}}}$ | 1 | $1+h_{f_{\mathrm{a}}}s_{f_{\mathrm{a}}}$ | $1+s_{f_{\mathrm{a}}}$ | 1 | $1+\eta_{f_{\mathrm{a}}}s_{f_{\mathrm{a}}}$ |
| $w_m$ | 1 | $1+h_m s_m$ | $1+s_m$ | 1 | $1+\eta_m s_m$ |

Table S3: Marginal fitness effects under different inheritance systems partitioned into the additive $a_1$ and non-additive $a_2$ components.

| | $a_1$ | $a_2$ |
|---|---|---|
| Autosomes | $\frac{1}{4}\left(h_{f_{\mathrm{g}}}s_{f_{\mathrm{g}}}+h_{f_{\mathrm{a}}}s_{f_{\mathrm{a}}}+2h_m s_m\right)$ | $\frac{1}{4}\left((1-2h_{f_{\mathrm{g}}})s_{f_{\mathrm{g}}}+(1-2h_{f_{\mathrm{a}}})s_{f_{\mathrm{a}}}+2(1-2h_m)s_m\right)$ |
| X | $\frac{1}{3}\left(h_{f_{\mathrm{g}}}s_{f_{\mathrm{g}}}+h_{f_{\mathrm{a}}}s_{f_{\mathrm{a}}}+s_m\right)$ | $\frac{1}{3}\left((1-2h_{f_{\mathrm{g}}})s_{f_{\mathrm{g}}}+(1-2h_{f_{\mathrm{a}}})s_{f_{\mathrm{a}}}\right)$ |
| PGE | $\frac{1}{3}\left(h_{f_{\mathrm{g}}}s_{f_{\mathrm{g}}}+h_{f_{\mathrm{a}}}s_{f_{\mathrm{a}}}+h_m s_m\right)$ | $\frac{1}{3}\left((1-2h_{f_{\mathrm{g}}})s_{f_{\mathrm{g}}}+(1-2h_{f_{\mathrm{a}}})s_{f_{\mathrm{a}}}+(1-2h_m)s_m\right)$ |
| $A_{\mathrm{a}}$ | $\frac{1}{7}\left(s_{f_{\mathrm{g}}}+2h_{f_{\mathrm{a}}}s_{f_{\mathrm{a}}}+4h_m s_m\right)$ | $\frac{1}{7}\left(2(1-2h_{f_{\mathrm{a}}})s_{f_{\mathrm{a}}}+4(1-2h_m)s_m\right)$ |
| $X_{\mathrm{a}}$ | $\frac{1}{5}\left(s_{f_{\mathrm{g}}}+2h_{f_{\mathrm{a}}}s_{f_{\mathrm{a}}}+2s_m\right)$ | $\frac{1}{5}\left(2(1-2h_{f_{\mathrm{a}}})s_{f_{\mathrm{a}}}\right)$ |
| $PGE_{\mathrm{a}}$ | $\frac{1}{5}\left(s_{f_{\mathrm{g}}}+2h_{f_{\mathrm{a}}}s_{f_{\mathrm{a}}}+2h_m s_m\right)$ | $\frac{1}{5}\left(2(1-2h_{f_{\mathrm{a}}})s_{f_{\mathrm{a}}}+2(1-2h_m)s_m\right)$ |

And the condition for the 0 allele to invade is:

$$I_0 = -\left(\lim_{p\to 1}\tilde{a}(p)\right) = -(a_1+a_2) > 0 \tag{S13}$$

Then we can see that our equilibria can be written as:

$$p^* = \frac{I_1}{I_0+I_1} \tag{S14}$$

As previously shown by Scott M. (personal communication).

#### 1.3 Effective mutation rate and the number of new mutations

The effective mutation rate experienced by a population is the mutation rate felt in different classes, multiplied by the fraction of evolutionary time spent in that particular state (Lehmann 2014):

$$\tilde{\mu} = \sum_i c_i \mu_i \tag{S15}$$

If, for example, we assume that males have a higher mutation rate than females, and describe this as so $\mu_{\mathrm{m}} = \mu_{\mathrm{f}}(1+\alpha)$ (Kirkpatrick and Hall, 2004). Then the relative difference in mutation rates between different portions of genome as a function of $\alpha$, will be determined by the relative portions of the time those genomes spend in males.

$$\frac{d}{d\alpha}\left(\frac{\tilde{\mu}_x}{\tilde{\mu}_y}\right) = \frac{c_{m,x}}{c_{m,y}} \tag{S16}$$

For some processes, the absolute number of new mutations that enter the population may matter. In this case, we can write the absolute number of new mutations that emerge in class $i$ as:

$$U_i = N_{\mathrm{i}}\gamma_i\mu_{\mathrm{j}} \tag{S17}$$

Where $\gamma_i$ is the ploidy of class $i$.

#### 1.4 Effective population size

We now calculate the effective population sizes under our different inheritance systems. If we assume that the transitions between classes occur at a rate much faster than the rate of coalescence, i.e a fast-time scale approximation of the coalescent process (Nordborg, 1997; Laporte and Charlesworth, 2002; Wakeley, 2009), then we may write the probability that two lineages coalesce at each time step $\tilde{Q}$ as a weighted average of the probability of coalescence between lineages in different classes:

$$\tilde{Q} = \sum_i \sum_j \left(u_i v_i \times u_j v_j\right) Q_{ij} = \sum_i \sum_j c_i c_j Q_{ij} \tag{S18}$$

Where $Q_{ij}$ is the probability that a gene copy in class $i$ and a gene copy in class $j$ descended from the same gene copy (i.e. coalesced) in the previous generation, and $c_i$ and $c_j$ are the reproductive values of these classes. These emerge because the probability of sampling an individual (or gene copy) of a particular class today is given by it's abundance $u_i$, and then the expected contribution to the future ancestry of that gene copy is given by it's reproductive value $v_i$, with the product of these being the reproductive value of the class $c_i$. These class reproductive values are equivalent to the $\alpha$'s in Laporte and Charlesworth (2002). We may then further write $Q_{ij}$ as:

$$Q_{ij} = \sum_k \frac{c_{k\to i}}{c_i} \frac{c_{k\to j}}{c_j} Q_{ijk} \tag{S19}$$

Where $c_{k\to i}$ is the reproductive value from class $k$ to class $i$, and $Q_{ijk}$ is the conditional probability that $i$ and $j$ coalesce, given that they both descend from class $k$. The product $\frac{c_{k\to i}}{c_i}\frac{c_{k\to j}}{c_j}$ equates to the $\beta$'s in Laporte and Charlesworth (2002). Combining these, allows the rate of coalescence to be expressed straightforwardly in terms of the elasticities:

$$\tilde{Q} = \sum_i \sum_j \sum_k c_{k\to i} c_{k\to j} Q_{ijk} = \sum_i \sum_j \sum_k e_{ik} e_{jk} Q_{ijk} \tag{S20}$$

Given that our two gene copies descended from the same class, the probability that they coalesce can be written as the probability that they come from the same individual $A_{ijk}$ (i.e. the probability of sibship (Gardner, 2010)), multiplied by the probability that, given $i$ and $j$ descend from the same individual in class $k$, they came from the same gene copy, $B_{ijk}$. This assumes that there is no covariance between these quantities:

$$Q_{ijk} = A_{ijk} \times B_{ijk} \tag{S21}$$

The $A$'s capture the demographic component, and are equivalent to the $Q$'s in Laporte and Charlesworth (2002), whilst the $B$'s capture the genetic component, the $\gamma$'s in Laporte and Charlesworth (2002). Thus the combinations of these can be seen as equivalent to their equation (13).

Focusing on the demographic component, we can calculate the probability of sibship by calculating the probability that two gene copies from class $i$ and $j$ came from the same individual in class $k$. If an individual $\mathbb{i}$ in class $k$ produces $b_{i_{\mathbb{i}}}$ offspring of class $i$ to adult age and $b_{j_{\mathbb{i}}}$ offspring of class $j$ to adult age, and there are in total $N_k$ individuals of class $k$, then the probability of sibship is:

$$A_{ijk} = \sum_{\mathbb{i}} \left(\frac{b_{ik_{\mathbb{i}}}}{N_k \bar{b}_{ik}}\right)\left(\frac{b_{jk_{\mathbb{i}}} - \delta_{ij}}{N_k \bar{b}_{jk} - \delta_{ij}}\right) \approx \sum_{\mathbb{i}} \left(\frac{b_{ik_{\mathbb{i}}}\left(b_{jk_{\mathbb{i}}} - \delta_{ij}\right)}{N_k^2 \bar{b}_{ik} \bar{b}_{jk}}\right) \tag{S22}$$

Where if $i = j$ then $\delta_{ij} = 1$ and if $i \neq j$ then $\delta_{ij} = 0$. We can write these in terms of variances and covariances (Caballero and Hill, 1992; Caballero, 1995), and can further simplify these by expressing these as deviations from the Poisson variance (Laporte and Charlesworth, 2002). When $i = j$, then:

$$\Delta V_{\mathrm{i}}^{\mathrm{k}} = V_{\mathrm{i}}^{\mathrm{k}} - \overline{b}_{\mathrm{ik}} \tag{S23a}$$

$$A_{\mathrm{iik}} \approx \frac{\sum_{\mathbb{i}} b_{\mathrm{ik}_{\mathbb{i}}}^2 - \sum_{\mathbb{i}} b_{\mathrm{ik}_{\mathbb{i}}}}{N_{\mathrm{k}}^2 \overline{b}_{\mathrm{ik}}^2} = \frac{\left(V_{\mathrm{i}}^{k} + \overline{b}_{\mathrm{ik}}^2\right) - \overline{b}_{\mathrm{ik}}}{N_{\mathrm{k}} \overline{b}_{\mathrm{ik}}^2} = \frac{1}{N_{\mathrm{k}}}\left(1 + \frac{\Delta V_{\mathrm{i}}^{\mathrm{k}}}{\overline{b}_{\mathrm{ik}}^2}\right) \tag{S23b}$$

Where $V_{\mathrm{i}}^{\mathrm{k}}$ is the variance in the number of class $i$ offspring produced by class $j$. If $i \neq j$, then:

$$A_{\mathrm{ijk}} = \frac{\sum_{\mathbb{i}} b_{\mathrm{ik}_{\mathbb{i}}} b_{\mathrm{jk}_{\mathbb{i}}}}{N_{\mathrm{k}}^2 \overline{b}_{\mathrm{ik}} \overline{b}_{\mathrm{jk}}} = \frac{C_{\mathrm{ij}}^{\mathrm{k}} + \overline{b}_{\mathrm{ik}} \overline{b}_{\mathrm{jk}}}{N_{\mathrm{k}} \overline{b}_{\mathrm{ik}} \overline{b}_{\mathrm{jk}}} = \frac{1}{N_{\mathrm{k}}}\left(1 + \frac{C_{\mathrm{ij}}^{\mathrm{k}}}{\overline{b}_{\mathrm{ik}} \overline{b}_{\mathrm{jk}}}\right) \tag{S24}$$

Where $C_{\mathrm{ij}}^{\mathrm{k}}$ is the covariance in the number of offspring in classes $i$ and $j$ produced by class $k$. Now, we can consider the probability of descending from the same particular gene copy within that individual. If the ploidy is 1, then $B_{\mathrm{ijk}} = 1$ as well. Else:

$$B_{\mathrm{ijk}} = \sum_{\mathbb{l}} \pi_{\mathrm{ik}_{\mathbb{l}}} \pi_{\mathrm{jk}_{\mathbb{l}}} \tag{S25}$$

Where $\pi_{ik_{\mathbb{l}}}$ is the probability of descending from $\mathbb{l}$th gene copy in the individual. If all gene copies are equally likely to be transmitted, then $B_{\mathrm{ijk}} = 1/\mathbb{P}_{\mathrm{k}}$ where $\mathbb{P}_{\mathrm{k}}$ is the ploidy of class $k$. Although exceptions may be possible.

Finally, we may note that, once we have computed our appropriately weighted probability of single generation coalescence $\tilde{Q}$, then we may use this to calculate our effective population size.

$$N_{\mathrm{e}} = \frac{1}{2\tilde{Q}} \tag{S26}$$

We now calculate explicit expressions for the effective population size for a digenic situation where there is a binomial distribution of sons vs daughters produced by a mother (Laporte and Charlesworth, 2002). We then consider a similar monogenic situation where there is a binomial distribution of gynogenic and androgenic daughters. In both cases we assume there are $N_{\mathrm{f}_{\mathbb{g}}}$ gynogenic females, $N_{\mathrm{f}_{\mathbb{a}}}$ androgenic females, and $N_{\mathrm{m}}$ males. In the digenic situation there are $N_{\mathrm{f}}$ females.

##### 1.4.1 Digenic scenario with binomial distribution of sons vs daughters

Following Laporte & Charlesworth (2002), we can more explicitly model a scenario where there is arbitrary variance in total offspring number, but there is a binomial distribution of sons and daughters, with $c$ being the probability of a son. We can calculate the variance in the number of sons a female produces as so:

$$\mathrm{Var}[b_{\mathrm{mf}}|b_{\mathrm{f}}] = b_{\mathrm{f}} c(1-c) \tag{S27a}$$

$$\mathrm{E}[\mathrm{Var}[b_{\mathrm{mf}}|b_{\mathrm{f}}]] = \frac{1}{N_{\mathrm{f}}} \sum_{\mathbb{i}} b_{\mathrm{f}_{\mathbb{i}}} c(1-c) = \overline{b}_{\mathrm{f}} c(1-c) \tag{S27b}$$

$$\mathrm{E}[b_{\mathrm{mf}}|b_{\mathrm{f}}] = c b_{\mathrm{f}} \tag{S27c}$$

$$\mathrm{Var}[\mathrm{E}[b_{\mathrm{mf}}|b_{\mathrm{f}}]] = \frac{1}{N_{\mathrm{f}}} \sum_{\mathbb{i}} \left(c b_{\mathrm{f}_{\mathbb{i}}}\right)^2 - \left(\frac{1}{N_{\mathrm{f}}} \sum_{\mathbb{i}} c b_{\mathrm{f}_{\mathbb{i}}}\right)^2 = c^2 \left(V_{\mathrm{f}} + \overline{b}_{\mathrm{f}}^2\right) - c^2 \overline{b}_{\mathrm{f}}^2 = c^2 V_{\mathrm{f}} \tag{S27d}$$

$$V_{\mathrm{m}}^{\mathrm{f}} = \mathrm{E}\left[\mathrm{Var}[b_{\mathrm{mf}}|b_{\mathrm{f}}]\right] + \mathrm{Var}\left[E[b_{\mathrm{mf}}|b_{\mathrm{f}}]\right] = c^2 V_{\mathrm{f}} + \overline{b}_{\mathrm{f}} c(1-c) \tag{S27e}$$

A similar approach gives us the variance in the number of daughters for a female:

$$V_{\mathrm{f}}^{\mathrm{f}} = \mathrm{E}\left[\mathrm{Var}[b_{\mathrm{ff}}|b_{\mathrm{f}}]\right] + \mathrm{Var}\left[E[b_{\mathrm{ff}}|b_{\mathrm{f}}]\right] = (1-c)^2 V_{\mathrm{f}} + \overline{b}_{\mathrm{f}} c(1-c) \tag{S28}$$

And the covariance between the number of sons and daughters:

$$\mathrm{E}[b_{\mathrm{mf}}|b_{\mathrm{f}}] = c b_{\mathrm{f}} \tag{S29}$$

$$\mathrm{E}[b_{\mathrm{ff}}|b_{\mathrm{f}}] = (1-c) b_{\mathrm{f}} \tag{S30}$$

$$\mathrm{Cov}[\mathrm{E}[b_{\mathrm{mf}}|b_{\mathrm{f}}], \mathrm{E}[b_{\mathrm{ff}}|b_{\mathrm{f}}]] = \left(\frac{1}{N_{\mathrm{f}}} \sum_{\mathbb{i}} b_{\mathrm{f}_{\mathbb{i}}}^2 c(1-c)\right) - \overline{b}_{\mathrm{f}}^2 c(1-c) = c(1-c) V_{\mathrm{f}} \tag{S31}$$

$$\mathrm{Cov}[b_{\mathrm{mf}}, b_{\mathrm{ff}}|b_{\mathrm{f}}] = -c(1-c) b_{\mathrm{f}} \tag{S32}$$

$$\mathrm{E}[\mathrm{Cov}[b_{\mathrm{mf}}, b_{\mathrm{ff}}|b_{\mathrm{f}}]] = \frac{1}{N_{\mathrm{f}}} \sum_{\mathbb{i}} -b_{\mathrm{f}_{\mathbb{i}}} c(1-c) = -c(1-c)\overline{b}_{\mathrm{f}} \tag{S33}$$

$$C_{\mathrm{fm}}^{\mathrm{f}} = \mathrm{E}[\mathrm{Cov}[b_{\mathrm{mf}}, b_{\mathrm{ff}}|b_{\mathrm{f}}]] + \mathrm{Cov}[\mathrm{E}[b_{\mathrm{mf}}|b_{\mathrm{f}}], \mathrm{E}[b_{\mathrm{ff}}|b_{\mathrm{f}}]] = (1-c)c\left(V_{\mathrm{f}} - \overline{b}_{\mathrm{f}}\right) \tag{S34}$$

Provided we have a stationary population, the Poisson variance in the total number of offspring produced by females is $\overline{b}_{\mathrm{f}} = 1/(1-c)$. Then we can rewrite these in terms of the deviations from the Poisson as before, where $\Delta V_{\mathrm{i}}^{\mathrm{k}} = V_{\mathrm{i}}^{\mathrm{k}} - \overline{b}_{\mathrm{ik}}$.

$$\Delta V_{\mathrm{m}}^{\mathrm{f}} = V_{\mathrm{m}}^{\mathrm{f}} - \frac{c}{1-c} = \left(c^2 V_{\mathrm{f}} + \frac{1}{1-c} c(1-c)\right) - \frac{c}{1-c} = c^2\left(V_{\mathrm{f}} - \frac{1}{1-c}\right) = c^2 \Delta V_{\mathrm{f}} \tag{S35}$$

$$\Delta V_{\mathrm{f}}^{\mathrm{f}} = V_{\mathrm{f}}^{\mathrm{f}} - \frac{1-c}{1-c} = \left((1-c)^2 V_{\mathrm{f}} + \frac{1}{1-c} c(1-c)\right) - \frac{c}{1-c} = c^2\left(V_{\mathrm{f}} + \frac{1}{1-c} - \frac{1}{c^2}\right) = (1-c)^2 \Delta V_{\mathrm{f}} \tag{S36}$$

$$C_{\mathrm{fm}}^{\mathrm{f}} = (1-c)c\left(\Delta V_{\mathrm{f}} + \frac{1}{1-c} - \frac{1}{1-c}\right) = c(1-c)\Delta V_{\mathrm{f}} \tag{S37}$$

We can follow a similar procedure to calculate the variance for contributions from males. Assembling this together we can then write out the expressions for the coalescence probabilities as so:

$$\tilde{Q}_A = \frac{1}{2N_{e,A}} = \frac{1}{8}(\mathbb{F} + \mathbb{M}) \tag{S38}$$

$$\tilde{Q}_X = \frac{1}{2N_{e,X}} = \frac{1}{9}(2\mathbb{F} + \mathbb{M}) \tag{S39}$$

Where:

$$\mathbb{F} = \frac{1 + (1-c)^2 \Delta V_{\mathrm{f}}}{N_{\mathrm{f}}} \tag{S40}$$

$$\mathbb{M} = \frac{1 + c^2 \Delta V_{\mathrm{m}}}{N_{\mathrm{m}}} \tag{S41}$$

We can see that these are equivalent to the results of Laporte & Charlesworth (2002) for the case of outbreeding. Setting the deviation in the variance from Poisson to zero, recovers the classic results of Wright (1933).

##### 1.4.2 Monogenic scenario with binomial distribution of gynogenic vs androgenic daughters

Now we develop a similar situation to above, but instead we have a binomial distribution of gynogenic and androgenic daughters, with $\delta$ being the probability of a gynogenic daughter. Taking a similar approach to above, the deviations from the Poisson variances are:

$$\Delta V_{\mathrm{f_g}}^{\mathrm{m}} = (1-c)^2\delta^2\Delta V_{\mathrm{m}} \tag{S42a}$$

$$\Delta V_{\mathrm{f_a}}^{\mathrm{m}} = (1-c)^2(1-\delta)^2\Delta V_{\mathrm{m}} \tag{S42b}$$

$$\Delta V_{\mathrm{m}}^{\mathrm{m}} = c^2\Delta V_{\mathrm{m}} \tag{S42c}$$

$$\Delta V_{\mathrm{f_g}}^{\mathrm{f_g}} = \delta^2\Delta V_{\mathrm{f_g}} \tag{S42d}$$

$$\Delta V_{\mathrm{f_a}}^{\mathrm{f_g}} = (1-\delta)^2\Delta V_{\mathrm{f_g}} \tag{S42e}$$

$$\Delta V_{\mathrm{m}}^{\mathrm{f_a}} = \Delta V_{\mathrm{f_a}} \tag{S42f}$$

And covariances:

$$C_{\mathrm{f_g f_a}}^{\mathrm{m}} = \delta(1-\delta)(1-c)^2\Delta V_{\mathrm{m}} \tag{S43a}$$

$$C_{\mathrm{f_g m}}^{\mathrm{m}} = \delta c(1-c)\Delta V_{\mathrm{m}} \tag{S43b}$$

$$C_{\mathrm{f_a m}}^{\mathrm{m}} = (1-\delta)c(1-c)\Delta V_{\mathrm{m}} \tag{S43c}$$

$$C_{\mathrm{f_g f_a}}^{\mathrm{f_g}} = \delta(1-\delta)\Delta V_{\mathrm{m}} \tag{S43d}$$

Again, combining these we can write out expression for the rate of coalescence and effective population size:

$$\tilde{Q}_A = \frac{1}{2N_{e,A}} = \frac{1}{32}(\mathbb{F}_{\mathrm{g}} + \mathbb{F}_{\mathrm{a}} + 4\mathbb{M}) \tag{S43e}$$

$$\tilde{Q}_X = \frac{1}{2N_{e,X}} = \frac{1}{18}(\mathbb{F}_{\mathrm{g}} + \mathbb{F}_{\mathrm{a}} + 2\mathbb{M}) \tag{S43f}$$

$$\tilde{Q}_M = \frac{1}{2N_{e,M}} = (1-\lambda)^4\,\mathbb{F}_{\mathrm{g}} + (1-\lambda)^2\,\lambda^2\mathbb{F}_{\mathrm{a}} + \lambda^2\mathbb{M} \tag{S43g}$$

$$\tilde{Q}_{A_{\mathrm{a}}} = \frac{1}{2N_{e,A_{\mathrm{a}}}} = \frac{1}{49}(\mathbb{F}_{\mathrm{g}} + 2\mathbb{F}_{\mathrm{a}} + 8\mathbb{M}) \tag{S43h}$$

$$\tilde{Q}_{X_{\mathrm{a}}} = \frac{1}{2N_{e,X_{\mathrm{a}}}} = \frac{1}{25}(\mathbb{F}_{\mathrm{g}} + 2\mathbb{F}_{\mathrm{a}} + 4\mathbb{M}) \tag{S43i}$$

$$\tilde{Q}_{A_{\mathrm{g}}} = \tilde{Q}_{X_{\mathrm{g}}} = \frac{1}{2N_{e,A_{\mathrm{g}}}} = \frac{1}{2N_{e,X_{\mathrm{g}}}} = \mathbb{F}_{\mathrm{g}} \tag{S43j}$$

Where:

$$\mathbb{F}_{\mathrm{g}} = \frac{1+\delta^2\Delta V_{\mathrm{f_g}}}{N_{\mathrm{f_g}}} \tag{S44a}$$

$$\mathbb{F}_{\mathrm{a}} = \frac{1+\left(\frac{1-c}{c}\right)^2(1-\delta)^2\Delta V_{\mathrm{f_a}}}{N_{\mathrm{f_a}}} \tag{S44b}$$

$$\mathbb{M} = \frac{1+c^2\Delta V_{\mathrm{m}}}{N_{\mathrm{m}}} \tag{S44c}$$

### 2 Diffusion Equations

#### 2.1 Advection and diffusion terms

Consider a locus with two alleles {0,1}, and symmetric mutation rates between these alleles. We can then write the advection term in our diffusion equation as:

$$A(p) \approx p(1-p)\tilde{a}(p) + \tilde{\mu}\left(1-2p\right) = p(1-p)[\tilde{a}_1 + \tilde{a}_2 p] + \tilde{\mu}\left(1-2p\right) \tag{S45}$$

And our diffusion term:

$$B(p) \approx \frac{p(1-p)}{2N_e} = p(1-p)\tilde{Q} \tag{S46}$$

Where the effective population size, mutation rate, and marginal fitness effects are as described earlier.

#### 2.2 Fixation probabilities and substitution rates

First, we consider the probability of fixation of our mutant allele 1, arising in background fixed for 0, at frequency $p_0$, and we ignore the further effects of mutation. The probability that our mutant allele fixes can be expressed as:

$$\Pi\left(p_0\right) = \frac{\int_0^{p_0} \psi\left(y\right) dy}{\int_0^1 \psi\left(y\right) dy} \tag{S47}$$

$$\psi\left(y\right) = \text{Exp}\left(-2\int_0^y \frac{A\left(x\right)}{B\left(x\right)} dx\right) \tag{S48}$$

We can now consider variations depending on the fitness effects of this mutant allele.

##### 2.2.1 Neutrality

First, consider the neutral case where $\tilde{a}_1 = \tilde{a}_2 = 0$. In this case:

$$\Pi\left(p_0\right) = p_0 \tag{S49}$$

i.e. the probability of fixation is simply the initial frequency of the allele. The initial frequency of the allele will depend on the size and composition of the population. The initial frequency of the allele given it arises in class $i$ will be:

$$p_0^i = \frac{c_i}{N_i \gamma_i} \tag{S50}$$

i.e. will depend on the relative size of that class, and relative share of the future that descend from it. The number of mutants in class $i$ is given by the absolute number of gene copies $N_i\gamma_i$, where $N_i$ is the number of individuals in class $i$, $\gamma_i$ is the ploidy of class $i$, and the mutation rate of that class $\mu_i$. Putting this together, gives us the neutral substitution rate:

$$K_{neu} = \sum_i N_i \gamma_i \mu_i \Pi\left(p_0^i\right) = \sum_{i,j} N_i \gamma_i \mu_i \frac{c_i}{N_i \gamma_i} = \sum_i c_i \mu_i = \tilde{\mu} \tag{S51}$$

And so we recover the result that the neutral substitution rate is simply given by the "effective" mutation rate (Lehmann, 2014). By comparing the substitution rate on portions of the genome which spend different time in males and females, inferences about male mutation bias may be made (Miyata et al., 1987; de Manuel et al., 2022).

##### 2.2.2 Additive

Now let's consider a "genic" model of selection ($\tilde{a}_2 = 0$), i.e. additivity. In this case our fixation probability then becomes:

$$\Pi\left(p_0\right) = \frac{1 - e^{-4\tilde{a}_1 N_e p_0}}{1 - e^{-4\tilde{a}_1 N_e}} = \frac{4\tilde{a}_1 N_e p_0}{1 - e^{-4\tilde{a}_1 N_e}} + O(p_0^2) \tag{S52}$$

Once again, we can incorporate differences in the sizes of classes as before. If we ignore terms of $O(p_0^2)$, which tends to be a highly accurate approximation, then:

$$K_{sel} = \sum_i N_i \gamma_i \mu_i \Pi\left(p_0^i\right) = \sum_i N_i \gamma_i \mu_i \frac{4\tilde{a}_1 N_e \left(c_i / N_i \gamma_i\right)}{1 - e^{-4\tilde{a}_1 N_e}} = \tilde{\mu}\left(\frac{4\tilde{a}_1 N_e}{1 - e^{-4\tilde{a}_1 N_e}}\right) \tag{S53}$$

##### 2.2.3 Non-additive

We can additionally incorporate non-additivity, and may express analytically in terms of error functions (Whitlock, 2003; Mrnjavac et al., 2023):

$$\Pi\left(p_0\right) = \frac{\mathrm{erf}\left(\frac{\sqrt{2}\sqrt{N_e}(\tilde{a}_1 + \tilde{a}_2 p_0)}{\sqrt{\tilde{a}_2}}\right) - \mathrm{erf}\left(\frac{\sqrt{2}\tilde{a}_1\sqrt{N_e}}{\sqrt{\tilde{a}_2}}\right)}{\mathrm{erf}\left(\frac{\sqrt{2}\sqrt{N_e}(\tilde{a}_1 + \tilde{a}_2)}{\sqrt{\tilde{a}_2}}\right) - \mathrm{erf}\left(\frac{\sqrt{2}\tilde{a}_1\sqrt{N_e}}{\sqrt{\tilde{a}_2}}\right)} = \frac{p_0\sqrt{\frac{8\tilde{a}_2 N_e}{\pi}}\, e^{-\frac{2\tilde{a}_1^2 N_e}{\tilde{a}_2}}}{\mathrm{erf}\left(\frac{\sqrt{2}\sqrt{N_e}(\tilde{a}_1 + \tilde{a}_2)}{\sqrt{\tilde{a}_2}}\right) - \mathrm{erf}\left(\frac{\sqrt{2}\tilde{a}_1\sqrt{N_e}}{\sqrt{\tilde{a}_2}}\right)} + O(p_0^2) \tag{S54}$$

As before, ignoring terms of $O(p_0^2)$, then we can write out substitution rate as:

$$K_{sel} = \sum_i N_i \gamma_i \mu_i \Pi\left(p_0^i\right) = \tilde{\mu}\frac{2\sqrt{\frac{2\tilde{a}_2 N_e}{\pi}}\, e^{-2\tilde{a}_1\left(\frac{a_1}{a_2}\right)N_e}}{\mathrm{erf}\left(\frac{\sqrt{2}\sqrt{N_e}(\tilde{a}_1 + \tilde{a}_2)}{\sqrt{\tilde{a}_2}}\right) - \mathrm{erf}\left(\frac{\sqrt{2}\tilde{a}_1\sqrt{N_e}}{\sqrt{\tilde{a}_2}}\right)} \tag{S55}$$

Which, again, will be proportional to $\tilde{\mu}$. Thus, we can see even with various sex-biases in mutation rates (and more complex class structure), $K_{sel}/K_{neu}$ will be essentially independent of any mutation bias. This recovers the finding of Vicoso and Charlesworth (2009).

#### 2.3 Stationary distributions

We may also calculate the stationary distribution for an allele under mutation-selection-drift balance $\phi(p)$. This is given by:

$$\phi\left(p\right) = \frac{C}{B\left(x\right)}\mathrm{Exp}\left(2\int \frac{A(p)}{B(p)}dp\right) \tag{S56}$$

Where the constant of integration, $C$, is computed such that $\int_0^1 \phi(p)dp = 1$. In general, it is not possible to compute an analytically closed form of $C$. For this reason we consider the neutral scenario, with which to compute neutral heterozygosity, and then consider some approximations for strongly deleterious mutations.

##### 2.3.1 Neutrality

If we first assume our allele is neutral ($\tilde{a}_1 = \tilde{a}_2 = 0$), and allowing for different mutation rates to $\mu$ and from $\nu$ our focal allele, then the frequency of that allele is given by:

$$\phi(p) = C(1-p)^{4N_e\nu - 1}p^{4N_e\mu - 1} \tag{S57}$$

$$\int_0^1 \phi(p)dp = 2N_e C\int_0^1 ((1-p)p)^{4\mu N_e - 1}dp = 2N_e C\frac{\Gamma(4N_e\mu)}{\Gamma(8N_e\mu)} = 1 \tag{S58}$$

We express compound parameters of the effective population size and mutation rate as $4N_e\tilde{\mu} = \theta$. Solving for our constant of integration, our stationary distribution becomes:

$$\phi\left(p\right) = \frac{\Gamma\left(2\theta\right)}{\Gamma\left(\theta\right)^2}\left((1-p)p\right)^{\theta-1} \tag{S59}$$

From this we can calculate the expected neutral heterozygosity as:

$$E[H] = \int_0^1 2p(1-p)\phi(p)dp = \frac{\theta}{1+2\theta} \tag{S60}$$

As autosomes and X chromosomes have the same effective population size, and would be expected to have the same mutation rate - as they are effected by sex-specific mutation bias in the same way - then their heterozygosity would expected to be equivalent too.

###### 2.3.2 Approximation for a strongly deleterious mutation

Nei (1968) makes the following approximation for the stationary distribution of a strongly deleterious allele. By first assuming that selection is strong enough such that we can ignore the effects of back mutation from the mutant allele to the wild-type, the stationary distribution becomes:

$$\phi\left(p\right) = 2CN_e(1-p)^2 p^{\theta-1} e^{2N_e ps((1-2h)p+2h)} \tag{S61}$$

Which provided that selection is strong enough such that the terms involving $p^2$ are much smaller than the those involving $p$, then Nei approximates the distribution $\phi(p)$ with by a gamma distribution, which may be further approximated if $\theta << 1$ by making the substitution that $\Gamma(\theta) \approx \theta^{-1}$ (Charlesworth et al., 2018).

$$\phi(p) = \frac{\zeta^\theta}{\Gamma(\theta)} e^{\zeta(-p)} p^{\theta-1} \approx \theta\zeta^\theta e^{-p\zeta} p^{\theta-1} \tag{S62}$$

As Nei (1968) states, the mean and variance of the allele frequency in this case become:

$$E[p] = \frac{\theta}{\zeta} = \frac{4N_e\tilde{\mu}}{4N_e\tilde{a}_1} = \frac{\tilde{\mu}}{\tilde{a}_1} \tag{S63}$$

With variance:

$$Var[p] = \frac{E[p]}{\zeta} \tag{S64}$$

So on the PGE autosomes, we expect both a higher frequency of deleterious alleles, and a higher variance in the deleterious allele frequency across loci.

##### 2.4 Adaptation from standing genetic variation

Using these results, we can also consider the probability that an adaptive variant fixes given its emerges from standing genetic variation. Focusing on variant which has a fitness effect of $\tilde{a}_b$ during the beneficial phase, and a fitness effect of $\tilde{a}_d$ during the deleterious phase. If there are initially $p_0$ copies of the allele in the population, then at least one of them goes to fixation can be approximated by:

$$\Pi_{\text{sgv}} \approx 1 - \text{Exp}\left(-2N_e\tilde{a}_b p_0\right) \tag{S65}$$

Orr and Betancourt (2001) use the deterministic approximation that the frequency of the deleterious allele is initially $p_0 = \mu / \tilde{a}_d$. When substituted in this gives the expression:

$$\Pi_{\text{sgv}} \approx 1 - \text{Exp}\left(-4N_e\tilde{\mu}\frac{\tilde{a}_b}{\tilde{a}_d}\right) \tag{S66}$$

As Hermisson and Pennings point out (2005), this may overestimate the probability of a fixation in such cases. Substituting in an approximation for the stationary distribution of alleles at mutation selection balance instead, they approximate the fixation probability from standing variation as:

$$\Pi_{\text{sgv}} \approx 1 - \text{Exp}\left(-4N_e\tilde{\mu}\text{Log}\left(1+R\right)\right) \tag{S67}$$

Where $R$ is the relative selective advantage during the beneficial phase:

$$R = \frac{\tilde{a}_b}{\tilde{a}_d + \frac{1}{4N_e}} \tag{S68}$$

We can see these expressions plotted for some different dominance values, and under difference genetic systems, in Figures S1 and S2. When dominance is equivalent during the beneficial and deleterious phase then, in general, autosomes will most readily fix beneficial mutations, and PGE autosomes and X chromosomes will be essentially equivalent. The weaker purging of deleterious variants can compensate for the lower probability that each one of them fixes.

These two effects may be uncoupled when reversals of dominance instead are assumed. If deleterious variants are recessive, but beneficial ones dominant, then this will relatively promote fixation rates on the autosomes and PGE autosome relative to the X chromosome, with the converse occurring if deleterious variants are dominant, and beneficial ones recessive.

Of such sweeps from standing genetic variation, one can calculate the probability that the sweep itself will be "hard" (a single copy fixes), or "soft" (multiple copies fix). The conditional probability of a "soft" sweep, given the sweep occurs is given by:

$$P_{\text{mult}} = \frac{1 - \left(1 + 4N_e p_0 \tilde{a}_b\right)\text{Exp}\left(-4N_e p_0 \tilde{a}_b\right)}{1 - \text{Exp}\left(-4N_e p_0 \tilde{a}_b\right)} \tag{S69}$$

for the deterministic mutation-selection balance. And for the stochastic mutation-selection balance by:

$$P_{\text{mult}} = 1 - \frac{\theta R/(1+R)}{(1+R)^{\theta} - 1} \tag{S70}$$

As previously given by Hermisson and Pennings (2005). Plots for the different genetic systems, and different values and assumptions about dominance can be seen in Figures S3 and S4. We can see that under equal dominance, PGE autosomes and X chromosomes are similar, with similar probabilities of a soft vs hard sweep, and with autosomes more likely to undergo a soft sweep (conditional on it occurring). However, with dominance reversals, then again the probability of a soft sweep becomes higher under PGE when deleterious mutations are recessive, and higher on X chromosomes when deleterious mutations are dominant.

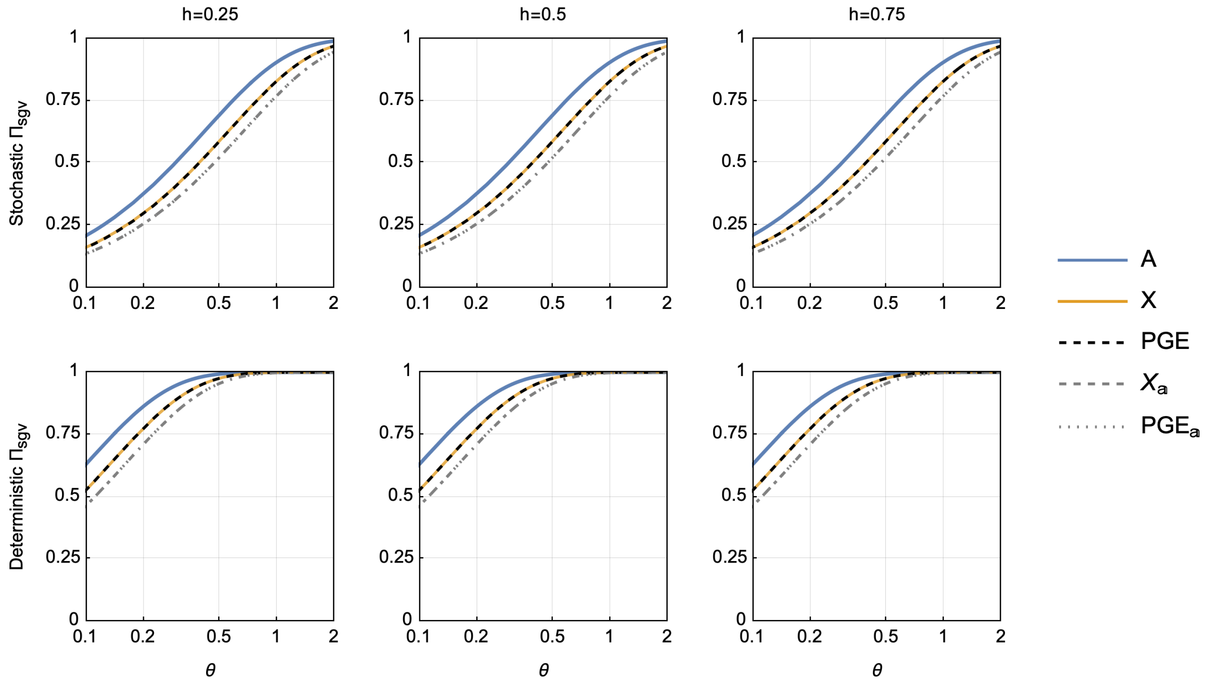


Figure S1: Probability that at least one allele fixes from the current standing genetic variation, and where dominance is assumed to be equal during the deleterious and beneficial phase such that $h_b = h_d = h$. During the deleterious phase the mutant allele imposes a cost of $s_d = 0.01$ and during the beneficial phase confers a benefit of $s_b = 0.1$, in both cases these are the effects in the homo/hemizygous form. The mutation rate is $\mu = 10^{-5}$.

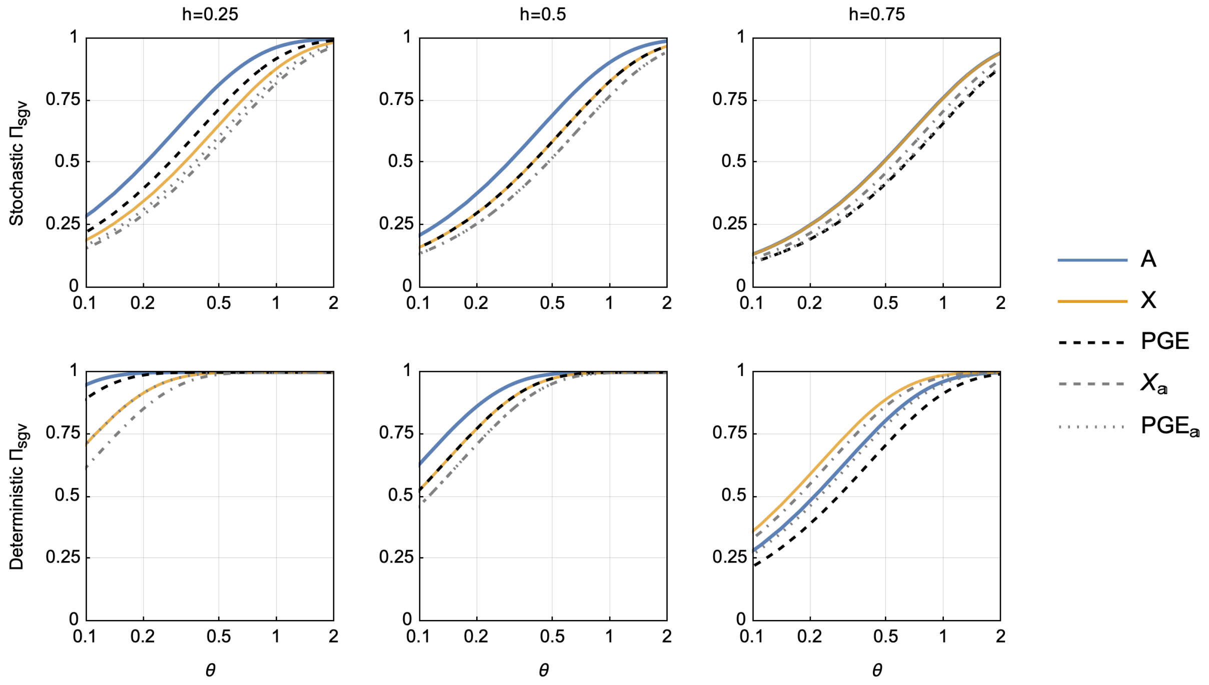


Figure S2: Probability that at least one allele fixes from the current standing genetic variation, and where dominance is assumed to be reversed during the deleterious and beneficial phase such that $1 - h_b = h_d = h$. During the deleterious phase the mutant allele imposes a cost of $s_d = 0.01$ and during the beneficial phase confers a benefit of $s_b = 0.1$, in both cases these are the effects in the homo/hemizygous form. The mutation rate is $\mu = 10^{-5}$.

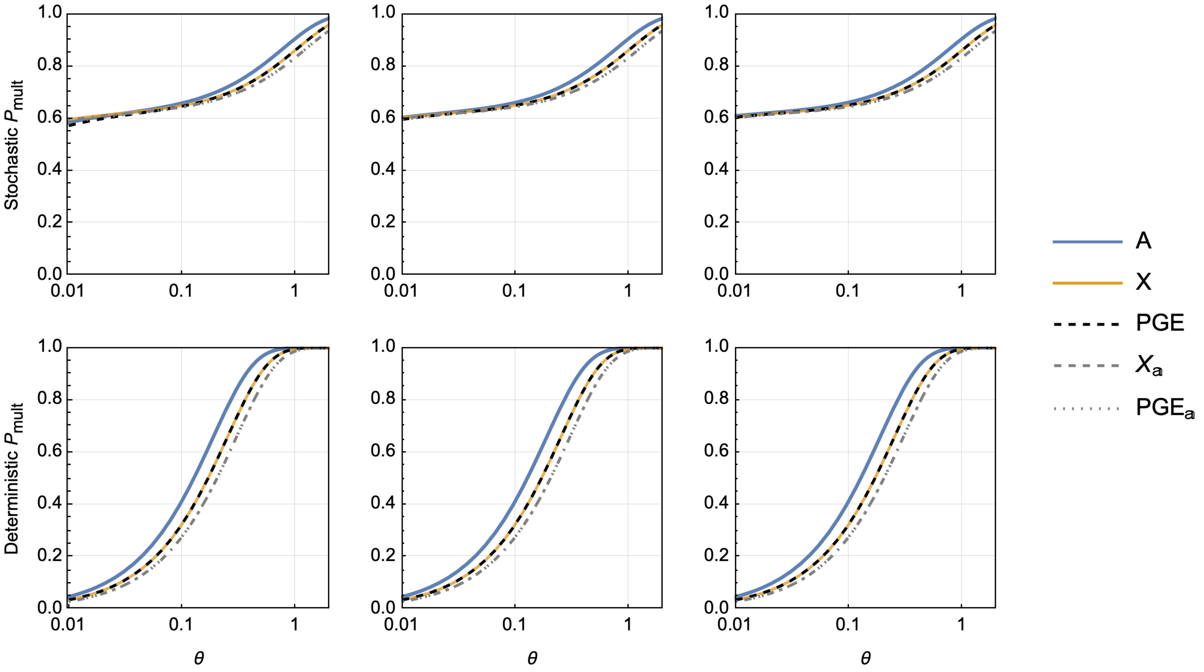


Figure S3: Probability that multiple copies from the standing genetic variation contribute to a substitution, and where dominance is assumed to be reversed during the deleterious and beneficial phase such that $h_b = h_d = h$. During the deleterious phase the mutant allele imposes a cost of $s_d = 0.01$ and during the beneficial phase confers a benefit of $s_b = 0.1$, in both cases these are the effects in the homo/hemizygous form. The mutation rate is $\mu = 10^{-5}$.

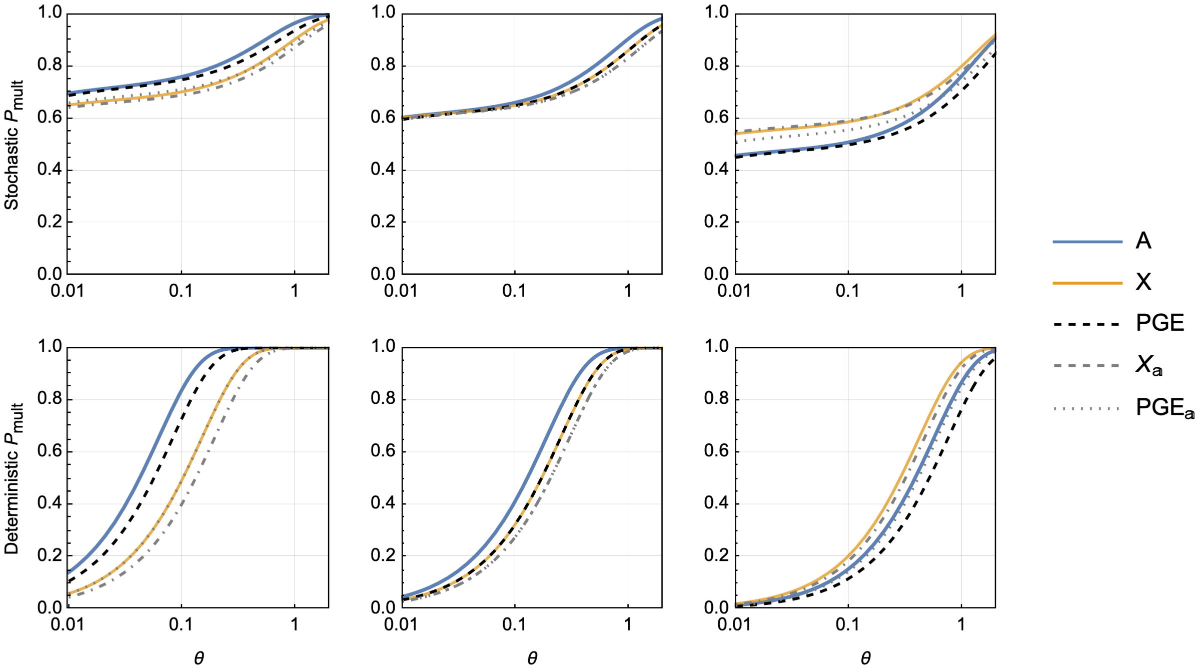


Figure S4: Probability that multiple copies from the standing genetic variation contribute to a substitution, and where dominance is assumed to be reversed during the deleterious and beneficial phase such that $1-h_b = h_d = h$. During the deleterious phase the mutant allele imposes a cost of $s_d = 0.01$ and during the beneficial phase confers a benefit of $s_b = 0.1$, in both cases these are the effects in the homo/hemizygous form. The mutation rate is $\mu = 10^{-5}$.

### 3 Branching process

An alternative approach to calculating the fixation probabilities of beneficial mutations is to describe the initial trajectory of the mutant lineage using a branching process (Fisher, 1923; Haldane, 1927; Patwa and Wahl, 2008). If the process is assumed to be supercritical, then there is a non-zero probability the lineage will go extinct. We can calculate this probability of non-extinction and this is assumed to be equivalent to the probability of fixation. When there are multiple classes or types in a population, then a multitype branching process may be used (Allen, 2010).

If we consider that there are a total of $k$ classes, then for each type $i$ a probability generating function can be written, describing the the offspring distribution $f_i\left(y_1, y_2, ..., y_k\right)$. From this, the expected number of class $j$ offspring that a gene copy in class $i$ produces can be written:

$$m_{\text{ij}} = \frac{\partial f_i(y_1, y_2, ..., y_k)}{\partial y_j}|_{y_1=1, y_2=1, ..., y_k=1} \tag{S71}$$

We can then write the expectation matrix $M = (m_{\text{ij}})_{i,j}$, composed of these elements. Using this matrix, we compute numerical solutions assuming that individuals produce independent, Poisson distributed numbers of offspring through each route. Additionally, we generate analytical approximations for the fixation probabilities, assuming selection is weak.

#### 3.1 Numerical solutions

If individuals produce an independent, Poisson distributed number of offspring through each route in the lifecycle, and we assume that the population is constant in size, then we can write the fixation probability for a new mutation in class $i$ as:

$$-\log(\Pi_i) = \sum_j m_{\text{ij}}\Pi_j \tag{S72}$$

This generates a set of $I$ equations for each of our states, which we can then jointly solve. With these assumptions, the expected number of offspring of class $j$ that an individual of class $i$ produces can be written as:

$$m_{\text{ij}} = t_{\text{ji}}\frac{N_j\gamma_j}{N_i\gamma_i} \tag{S73}$$

Where $N_i$ is the number of individuals of class $i$, $\gamma_i$ is the ploidy of class $i$, and $t_{\text{ji}}$ is the probability a gene copy in class $j$ came from class $i$ in the previous generation. Here we will assume that if the population is monogenic then there are an equal number of androgenic and gynogenic females $N_{f_\text{g}} = N_{f_\text{a}}$, and that the ratio of males to females in the population is given by $\rho = N_m/N_f$. We can then write out the system of equations for our different genetic systems of interest. The expressions and the weak selection approximations for these different systems can be seen below in section 3.2.

We numerically solve these systems of equations, identifying solutions between 0 and 1 using Mathematica's FindRoot function. These results are developed for different dominance scenarios, sex ratios, and strength of selection. This can be seen in Figures S5-S8.

###### 3.1.1 Autosomes

$$-\log(1-\Pi_\mathrm{f}) = (1+hs_\mathrm{f})\left(\frac{1}{2}\Pi_\mathrm{f} + \frac{\rho}{2}\Pi_\mathrm{m}\right) \tag{S74a}$$

$$-\log(1-\Pi_\mathrm{m}) = (1+hs_\mathrm{m})\left(\frac{1}{2\rho}\Pi_\mathrm{f} + \frac{1}{2}\Pi_\mathrm{m}\right) \tag{S74b}$$

###### 3.1.2 X chromosomes

$$-\log(1-\Pi_\mathrm{f}) = (1+hs_\mathrm{f})\left(\frac{1}{2}\Pi_\mathrm{f} + \frac{\rho}{2}\Pi_\mathrm{m}\right) \tag{S75a}$$

$$-\log(1-\Pi_\mathrm{m}) = (1+s_\mathrm{m})\left(\frac{1}{\rho}\Pi_\mathrm{f}\right) \tag{S75b}$$

###### 3.1.3 PGE Autosomes

$$-\log(1-\Pi_\mathrm{f}) = (1+hs_\mathrm{f})\left(\frac{1}{2}\Pi_\mathrm{f} + \frac{\rho}{2}\Pi_\mathrm{m}\right) \tag{S76a}$$

$$-\log(1-\Pi_\mathrm{m}) = (1+hs_\mathrm{m})\left(\frac{1}{\rho}\Pi_\mathrm{f}\right) \tag{S76b}$$

###### 3.1.4 Autosomal androgenic

For the androgenic portion in a standard autosomal system:

$$-\log\left(1-\Pi_{\mathrm{f_g}}\right) = (1+s_\mathrm{f})\left(\Pi_{\mathrm{f_a}}\right) \tag{S77a}$$

$$-\log\left(1-\Pi_{\mathrm{f_a}}\right) = (1+hs_\mathrm{f})\left(\rho\Pi_\mathrm{m}\right) \tag{S77b}$$

$$-\log(1-\Pi_\mathrm{m}) = (1+hs_\mathrm{m})\left(\frac{1}{4\rho}\Pi_{\mathrm{f_g}} + \frac{1}{4\rho}\Pi_{\mathrm{f_a}} + \frac{1}{2}\Pi_\mathrm{m}\right) \tag{S77c}$$

###### 3.1.5 X chromosome androgenic

$$-\log\left(1-\Pi_{\mathrm{f_g}}\right) = (1+s_\mathrm{f})\left(\Pi_{\mathrm{f_a}}\right) \tag{S78a}$$

$$-\log\left(1-\Pi_{\mathrm{f_a}}\right) = (1+hs_\mathrm{f})\left(\rho\Pi_\mathrm{m}\right) \tag{S78b}$$

$$-\log(1-\Pi_\mathrm{m}) = (1+s_\mathrm{m})\left(\frac{1}{2\rho}\Pi_{\mathrm{f_g}} + \frac{1}{2\rho}\Pi_{\mathrm{f_a}}\right) \tag{S78c}$$

###### 3.1.6 PGE androgenic

$$-\log\left(1-\Pi_{\mathrm{f_g}}\right) = (1+s_\mathrm{f})\left(\Pi_{\mathrm{f_a}}\right) \tag{S79a}$$

$$-\log\left(1-\Pi_{\mathrm{f_a}}\right) = (1+hs_\mathrm{f})\left(\rho\Pi_\mathrm{m}\right) \tag{S79b}$$

$$-\log(1-\Pi_\mathrm{m}) = (1+hs_\mathrm{m})\left(\frac{1}{2\rho}\Pi_{\mathrm{f_g}} + \frac{1}{2\rho}\Pi_{\mathrm{f_a}}\right) \tag{S79c}$$

#### 3.2 Weak selection approximation

We now generate analytical approximations under weak selection using a result from Hoppe (1992) and Pollak (1992). First,we can rewrite the expectation matrix in the following way:

$$\mathbf{M} = (\mathbf{I} + \mathbf{S})\mathbf{M}_0 \tag{S80}$$

Where $\mathbf{M}_0$ is expectation matrix is there were no selection, and our mutation was neutral, and then $\mathbf{S}$ is a diagonal matrix of our selective effects of the mutation when rare in the different classes. We can note that transpose of $\mathbf{M}_0$ is equivalent to a projection matrix $P$ (Caswell, 2000). We can then write out the left and right eigenvectors associated with the dominant eigenvalue of this matrix as:

$$Pu = uM_0 = \lambda u \tag{S81a}$$

$$vP = M_0 v = \lambda v \tag{S81b}$$

Where $u$ is the stable class distribution and $v$ are the individual reproductive values. We can then normalise these such that the total reproductive value sums to 1, and the total abundance sums to the number of haploid gene copies in the population.

$$\sum_i u_i v_i = 1 \tag{S82a}$$

$$\sum_i u_i = N_h \tag{S82b}$$

Pollak (1992) uses a different normalisation, we will write these differently normalised quantities as $\mathbb{u}$ and $\mathbb{v}$ respectively.

$$\sum_i \mathbb{u}_i \mathbb{v}_i = 1 \tag{S83a}$$

$$\sum_i \mathbb{u}_i = 1 \tag{S83b}$$

Pollak approximates the fixation probability for a mutation in class $i$ as:

$$\Pi_i \approx \frac{2\mathbb{u}.\mathbf{S}.\mathbb{v}}{\sum_i \mathbb{u}_i \left(\mathrm{Var}(\sum_j \mathbb{v}_j m_{\mathrm{ij}})\right)} \mathbb{v}_i \tag{S84}$$

We can make the following substitutions. First, the numerator can be realised to be equivalent to the weighted marginal fitness effect when the mutant is rare in the population.

$$\mathbb{u}.\mathbf{S}.\mathbb{v} = \tilde{a}(0) \tag{S85}$$

Second, Pollak in a later paper, recognised that the denominator of this expression is an approximation for the effective population size (Pollak, 2000).

$$\frac{N_h}{2N_e} \approx \sum_i \mathbb{u}_i \left(\mathrm{Var}(\sum_j \mathbb{v}_j m_{\mathrm{ij}})\right) \tag{S86}$$

Finally, we can replace Pollak's normalisation of reproductive value with the one described here:

$$\mathbb{v}_i = v_i N_h \tag{S87}$$

Substituting these in gives us:

$$\Pi_i \approx \frac{2\mathbb{u}.\mathbf{S}.\mathbb{v}}{\sum_i \mathbb{u}_i \left(\mathrm{Var}(\sum_j \mathbb{v}_j m_{\mathrm{ij}})\right)} \mathbb{v}_i = 4N_e \tilde{a}(0) v_i \tag{S88}$$

Table S4: Weak selection approximations for the fixation of a beneficial new mutation that arises in a specific class. The ratio of males to females in the population is given by $\rho = N_m/N_f$, and it is assumed that there are an equal number of gynogenic and androgenic females. Mutations are assumed to be equally beneficial in all classes of individual $s_{\mathrm{f_g}} = s_{\mathrm{f_a}} = s_{\mathrm{m}} = s$. In the case of PGE, the fixation probability in males refers to a new mutation on the maternal-origin copy, new mutations on the paternal-origin copy are ignored.

| | $\Pi_{f_{\mathrm{a}}}$ | $\Pi_{f_{\mathrm{g}}}$ | $\Pi_m$ |
|---|---|---|---|
| A | $\frac{4h\rho s}{\rho+1}$ | | $\frac{4hs}{\rho+1}$ |
| X | $\frac{2(2h+1)\rho s}{2\rho+1}$ | | $\frac{2(2h+1)s}{2\rho+1}$ |
| PGE | $\frac{6h\rho s}{2\rho+1}$ | | $\frac{6hs}{2\rho+1}$ |
| $A_{\mathrm{a}}$ | $\frac{2(6h+1)\rho s}{3\rho+4}$ | $\frac{2(6h+1)\rho s}{3\rho+4}$ | $\frac{2(6h+1)s}{3\rho+4}$ |
| $X_{\mathrm{a}}$ | $\frac{2(2h+3)\rho s}{3\rho+2}$ | $\frac{2(2h+3)\rho s}{3\rho+2}$ | $\frac{2(2h+3)s}{3\rho+2}$ |
| $\mathrm{PGE}_{\mathrm{a}}$ | $\frac{2(4h+1)\rho s}{3\rho+2}$ | $\frac{2(4h+1)\rho s}{3\rho+2}$ | $\frac{2(4h+1)s}{3\rho+2}$ |

Fixation probabilities for our different systems can be found in Table S4. We plot the fixation probabilities for some of our different genetic systems, under different dominance coefficients and sex ratios, and under different strengths of selection, and compare these to the numerical results. In general, provided selection is weak, then these approximations are very close to the numerical solutions.

##### 3.3 Substitution rates

With the fixation probabilities, we can then compute the substitution rates. If the absolute number of mutations that arises in class $i$ is $U_i$, which is a product of the size of the class $N_i\gamma_i$, and the mutation rate in that class $\mu_i$, then the the substitution rate will be:

$$K = \sum_i U_i \times \Pi_i = \sum_i N_i\gamma_i\mu_i \times 4N_e\tilde{a}(0)v_i = 4N_e\tilde{a}(0)\sum_i N_i\gamma_i\mu_i \times \frac{c_i}{N_i\gamma_i} = 4N_e\tilde{a}(0)\tilde{\mu} \tag{S89}$$

Where the $\tilde{\mu}$ that emerges is equivalent to the effective mutation rate computed earlier. The expression also agrees with the approximation from the diffusion analysis (Kimura and Ohta, 2020). We can then compute and compare for our different genetic systems. Focusing on Poisson distributed numbers of offspring, these can be seen in Table S4.

Table S5: Weak selection approximations for the adaptive substitution rates on different portions of the genome, for different classes of genes including: genes expressed in both sexes, female limited and male limited.

| | Both | Female-limited | Male-limited |
|---|---|---|---|
| Autosomal | $4nhs$ | $2nhs$ | $2nhs$ |
| X linked | $n(1+2h)s$ | $2nhs$ | $ns$ |
| PGE | $3nhs$ | $2nhs$ | $nhs$ |
| $A_{\mathbf{a}}$ | $\frac{1}{2}n(6h+1)s$ | $\frac{1}{2}n(2h+1)s$ | $2nhs$ |
| $X_{\mathbf{a}}$ | $\frac{1}{2}n(2h+3)s$ | $\frac{1}{2}n(2h+1)s$ | $ns$ |
| PGE$_{\mathbf{a}}$ | $\frac{1}{2}n(4h+1)s$ | $\frac{1}{2}n(2h+1)s$ | $nhs$ |

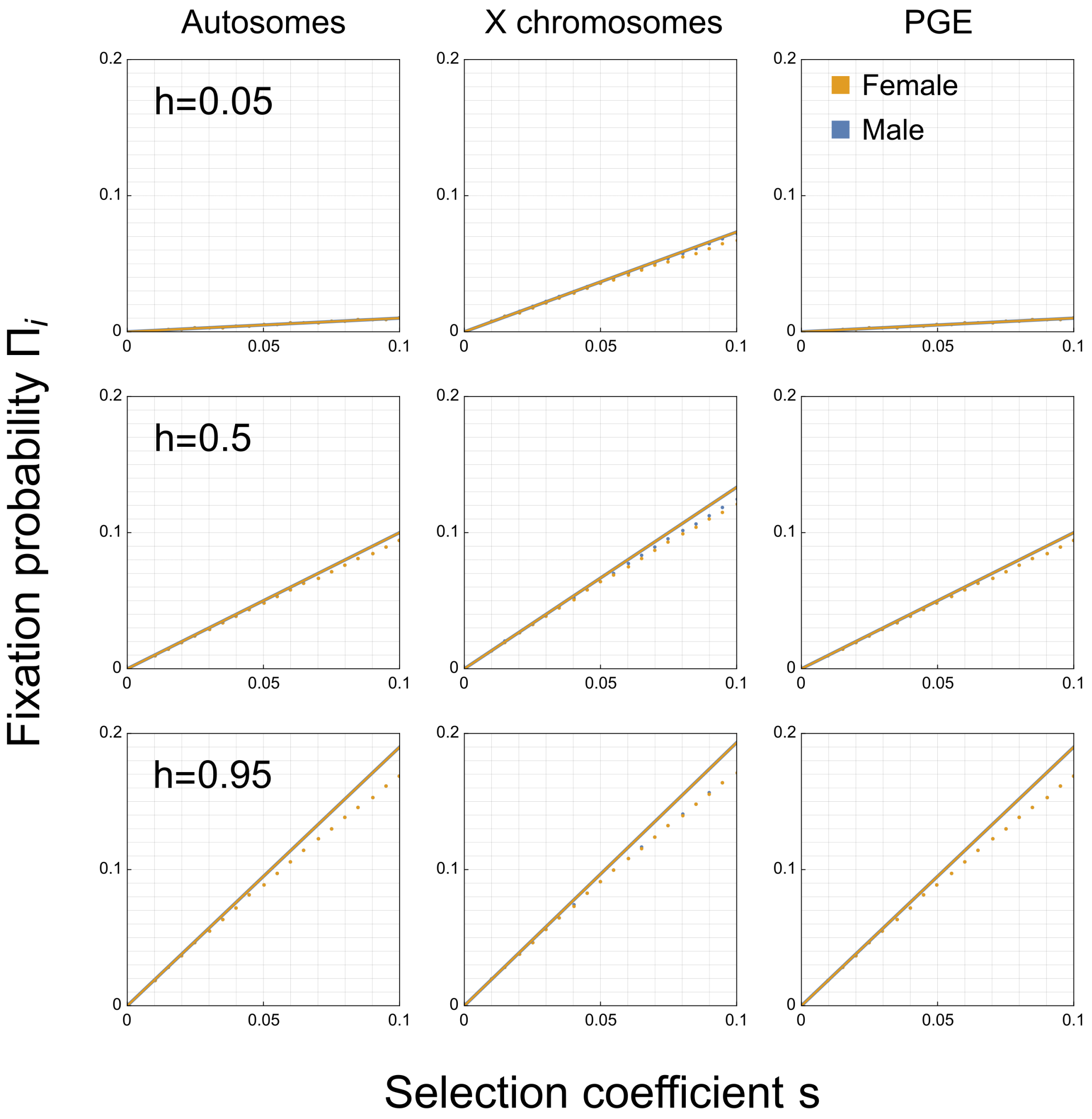


Figure S5: Numerical solutions (points) and analytical approximations (solid lines) for the fixation probability of a new, beneficial mutation arising in males and females under different genetic systems. Selection coefficients are assumed to be equal in males and females, and there are an equal number of males and females in the population. Individuals produce a Poisson distributed number of offspring.

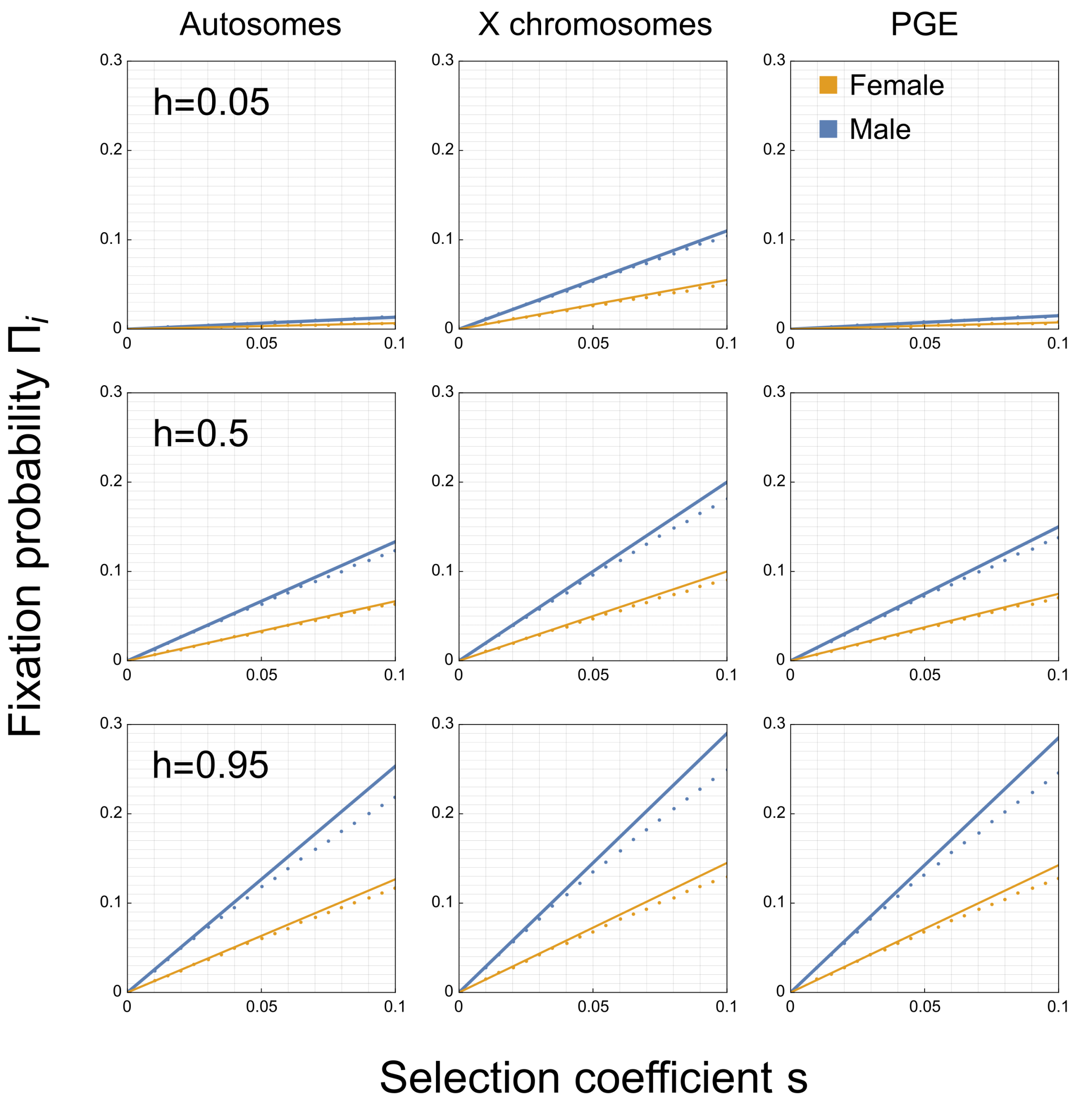


Figure S6: Numerical solutions (points) and analytical approximations (solid lines) for the fixation probability of a new, beneficial mutation arising in males and females under different genetic systems. Selection coefficients are assumed to be equal in males and females, and there are assumed to be twice as many females in the population as males. Individuals produce a Poisson distributed number of offspring.

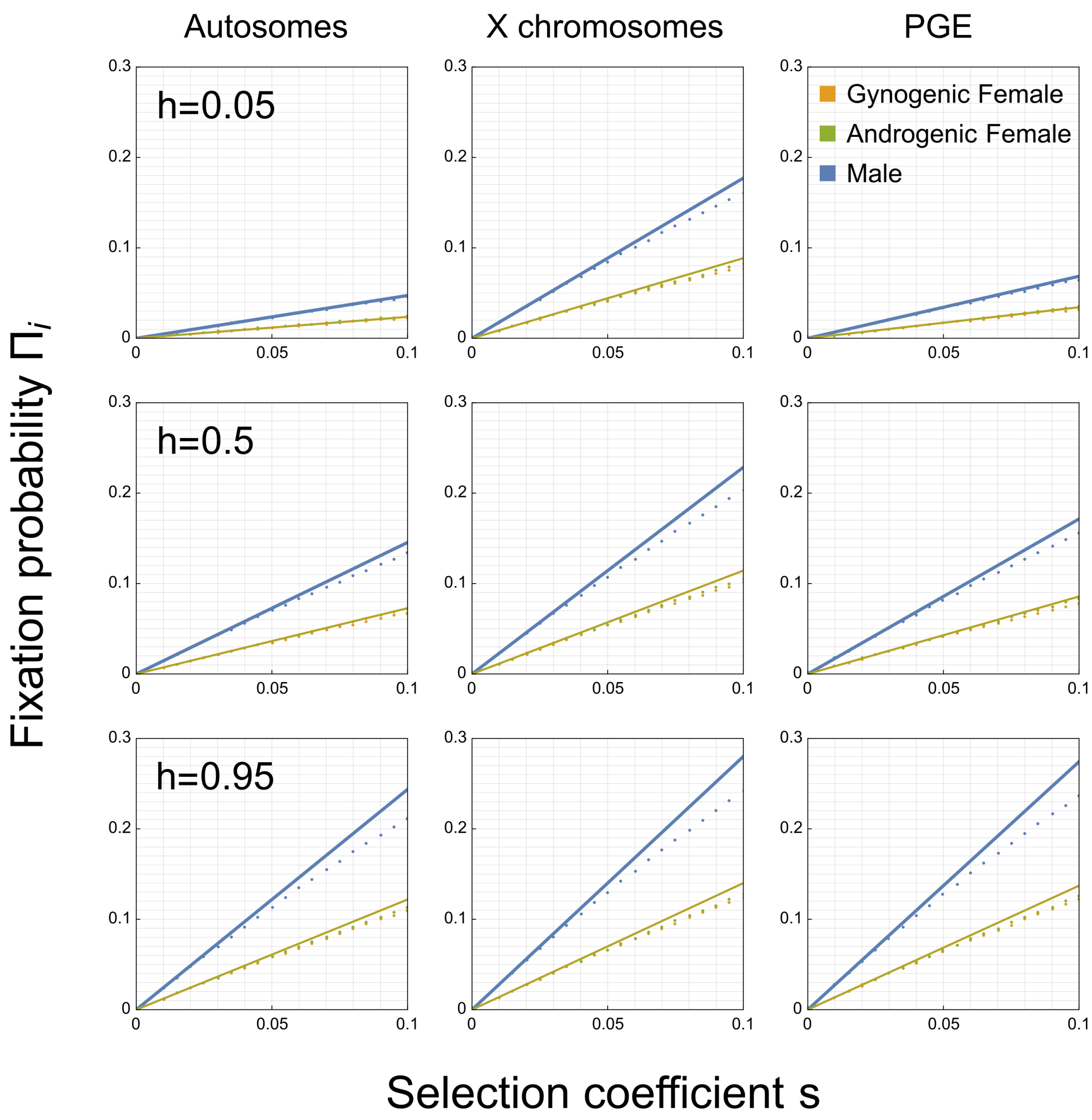


Figure S7: Numerical solutions (points) and analytical approximations (solid lines) for the fixation probability of a new, beneficial mutation arising in males, gynogenic females and androgenic females under different genetic systems. Selection coefficients are assumed to be equal in all classes, and there are an equal number of males and females in the population, and an equal number of gynogenic and androgenic females. Individuals produce a Poisson distributed number of offspring.

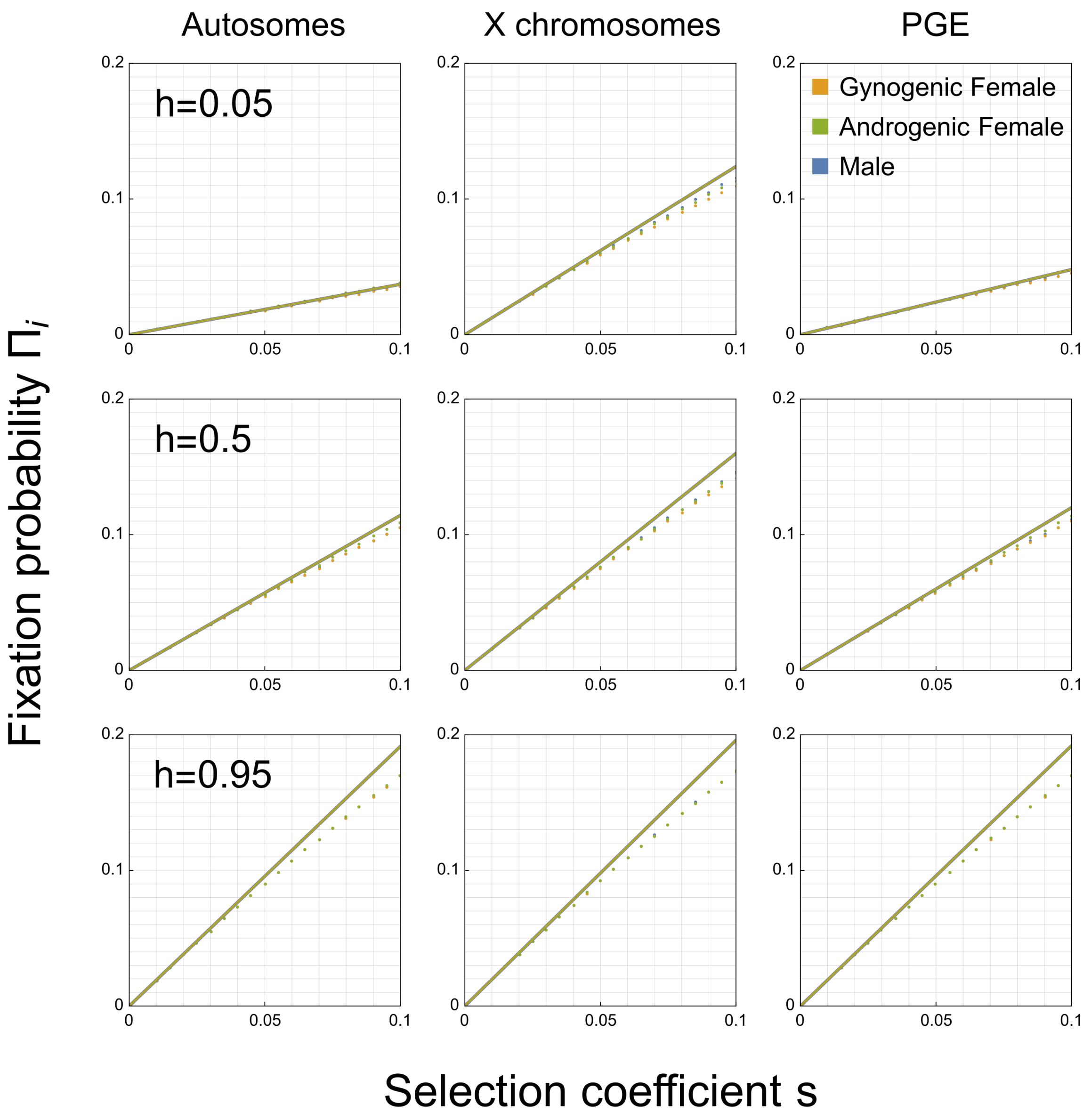


Figure S8: Numerical solutions (points) and analytical approximations (solid lines) for the fixation probability of a new, beneficial mutation arising in males and females under different genetic systems. Selection coefficients are assumed to be equal in all classes, and there are assumed to be twice as many females in the population as males, but an equal number of gynogenic and androgenic females. Individuals produce a Poisson distributed number of offspring.

### 4 Deterministic models

We now consider some results for selection in infinite populations, and thus ignore the effects of genetic drift. First we write out the recursions for the allele frequency in different classes, under different genetic systems. We then use these to calculate the allele frequency at mutation-selection balance, and then use these expressions to calculate the genetic load that the deleterious mutation imposes on different classes. We then we calculate the invasion conditions for sexually antagonistic and maternally antagonistic alleles. In both cases, by assuming selection is weak, we then calculate the equilibrium allele frequency if a polymorphism is maintained, and express the space for polymorphism as the angle between the two invasion boundaries

#### 4.1 Allele frequency recursions

First, we write out recursions for the allele frequency in our different systems, where $p_i$ is the allele frequency in class $i$. The allele frequencies are censused in the gametes of our different classes of individual, so for example $p_{\mathrm{f}}$ corresponds to the frequency in eggs. The life cycle proceeds in the following order:(1) fertilisation, (2) selection, (3) mutation. We notate the allele frequency in the gametes after phase of selection as $\hat{p}_i$.

##### 4.1.1 Autosomes

$$\hat{p}_{\mathrm{f}} = \frac{\frac{1}{2} w^{\mathrm{f}}_{01}\left(p_{\mathrm{f}}\left(1-p_{\mathrm{m}}\right)+\left(1-p_{\mathrm{f}}\right)p_{\mathrm{m}}\right)+p_{\mathrm{f}}p_{\mathrm{m}}w^{\mathrm{f}}_{11}}{w^{\mathrm{f}}_{01}\left(p_{\mathrm{f}}\left(1-p_{\mathrm{m}}\right)+\left(1-p_{\mathrm{f}}\right)p_{\mathrm{m}}\right)+w^{\mathrm{f}}_{11}p_{\mathrm{f}}p_{\mathrm{m}}+w^{\mathrm{f}}_{00}\left(1-p_{\mathrm{f}}\right)\left(1-p_{\mathrm{m}}\right)} \tag{S90a}$$

$$\hat{p}_{\mathrm{m}} = \frac{\frac{1}{2} w^{\mathrm{m}}_{01}\left(p_{\mathrm{f}}\left(1-p_{\mathrm{m}}\right)+\left(1-p_{\mathrm{f}}\right)p_{\mathrm{m}}\right)+p_{\mathrm{f}}p_{\mathrm{m}}w^{\mathrm{m}}_{11}}{w^{\mathrm{m}}_{01}\left(p_{\mathrm{f}}\left(1-p_{\mathrm{m}}\right)+\left(1-p_{\mathrm{f}}\right)p_{\mathrm{m}}\right)+w^{\mathrm{m}}_{11}p_{\mathrm{f}}p_{\mathrm{m}}+w^{\mathrm{m}}_{00}\left(1-p_{\mathrm{f}}\right)\left(1-p_{\mathrm{m}}\right)} \tag{S90b}$$

$$p'_{\mathrm{f}} = \hat{p}_{\mathrm{f}}(1-\mu_{\mathrm{f}})+(1-\hat{p}_{\mathrm{f}})\mu_{\mathrm{f}} \tag{S90c}$$

$$p'_{\mathrm{m}} = \hat{p}_{\mathrm{m}}(1-\mu_{\mathrm{m}})+(1-\hat{p}_{\mathrm{m}})\mu_{\mathrm{m}} \tag{S90d}$$

##### 4.1.2 X chromosomes

$$\hat{p}_{\mathrm{f}} = \frac{\frac{1}{2} w^{\mathrm{f}}_{01}\left(p_{\mathrm{f}}\left(1-p_{\mathrm{m}}\right)+\left(1-p_{\mathrm{f}}\right)p_{\mathrm{m}}\right)+p_{\mathrm{f}}p_{\mathrm{m}}w^{\mathrm{f}}_{11}}{w^{\mathrm{f}}_{01}\left(p_{\mathrm{f}}\left(1-p_{\mathrm{m}}\right)+\left(1-p_{\mathrm{f}}\right)p_{\mathrm{m}}\right)+w^{\mathrm{f}}_{11}p_{\mathrm{f}}p_{\mathrm{m}}+w^{\mathrm{f}}_{00}\left(1-p_{\mathrm{f}}\right)\left(1-p_{\mathrm{m}}\right)} \tag{S91a}$$

$$\hat{p}_{\mathrm{m}} = \frac{w^{\mathrm{m}}_{1}p_{\mathrm{f}}}{w^{\mathrm{m}}_{1}p_{\mathrm{f}}+w^{\mathrm{m}}_{0}\left(1-p_{\mathrm{f}}\right)} \tag{S91b}$$

$$p'_{\mathrm{f}} = \hat{p}_{\mathrm{f}}(1-\mu_{\mathrm{f}})+(1-\hat{p}_{\mathrm{f}})\mu_{\mathrm{f}} \tag{S91c}$$

$$p'_{\mathrm{m}} = \hat{p}_{\mathrm{m}}(1-\mu_{\mathrm{m}})+(1-\hat{p}_{\mathrm{m}})\mu_{\mathrm{m}} \tag{S91d}$$

##### 4.1.3 Paternal genome elimination

$$\hat{p}_{\mathrm{f}} = \frac{\frac{1}{2} w^{\mathrm{f}}_{01}\left(p_{\mathrm{f}}\left(1-p_{\mathrm{m}}\right)+\left(1-p_{\mathrm{f}}\right)p_{\mathrm{m}}\right)+p_{\mathrm{f}}p_{\mathrm{m}}w^{\mathrm{f}}_{11}}{w^{\mathrm{f}}_{01}\left(p_{\mathrm{f}}\left(1-p_{\mathrm{m}}\right)+\left(1-p_{\mathrm{f}}\right)p_{\mathrm{m}}\right)+w^{\mathrm{f}}_{11}p_{\mathrm{f}}p_{\mathrm{m}}+w^{\mathrm{f}}_{00}\left(1-p_{\mathrm{f}}\right)\left(1-p_{\mathrm{m}}\right)} \tag{S92a}$$

$$\hat{p}_{\mathrm{m}} = \frac{w^{\mathrm{m}}_{01}p_{\mathrm{f}}\left(1-p_{\mathrm{m}}\right)+p_{\mathrm{f}}p_{\mathrm{m}}w^{\mathrm{m}}_{11}}{w^{\mathrm{m}}_{01}\left(p_{\mathrm{f}}\left(1-p_{\mathrm{m}}\right)+\left(1-p_{\mathrm{f}}\right)p_{\mathrm{m}}\right)+w^{\mathrm{m}}_{11}p_{\mathrm{f}}p_{\mathrm{m}}+w^{\mathrm{m}}_{00}\left(1-p_{\mathrm{f}}\right)\left(1-p_{\mathrm{m}}\right)} \tag{S92b}$$

$$p'_{\mathrm{f}} = \hat{p}_{\mathrm{f}}(1-\mu_{\mathrm{f}}) + (1-\hat{p}_{\mathrm{f}})\mu_{\mathrm{f}} \tag{S92c}$$

$$p'_{\mathrm{m}} = \hat{p}_{\mathrm{m}}(1-\mu_{\mathrm{m}}) + (1-\hat{p}_{\mathrm{m}})\mu_{\mathrm{m}} \tag{S92d}$$

###### 4.1.4 Autosomal androgenic

$$\hat{p}_{\mathrm{f_g}} = \frac{w_1^{\mathrm{f_g}} p_{\mathrm{m}}}{w_1^{\mathrm{f_g}} + w_0^{\mathrm{f_g}}\left(1-p_{\mathrm{m}}\right)} \tag{S93a}$$

$$\hat{p}_{\mathrm{f_a}} = \frac{\frac{1}{2} w_{01}^{\mathrm{f_a}}\left(p_{\mathrm{f_g}}\left(1-p_{\mathrm{m}}\right) + \left(1-p_{\mathrm{f_g}}\right)p_{\mathrm{m}}\right) + p_{\mathrm{f_g}} p_{\mathrm{m}} w_{11}^{\mathrm{f_a}}}{w_{01}^{\mathrm{f_a}}\left(p_{\mathrm{f_g}}\left(1-p_{\mathrm{m}}\right) + \left(1-p_{\mathrm{f_g}}\right)p_{\mathrm{m}}\right) + w_{11}^{\mathrm{f_a}} p_{\mathrm{f_g}} p_{\mathrm{m}} + w_{00}^{\mathrm{f_a}}\left(1-p_{\mathrm{f_g}}\right)\left(1-p_{\mathrm{m}}\right)} \tag{S93b}$$

$$\hat{p}_{\mathrm{m}} = \frac{\frac{1}{2} w_{01}^{\mathrm{m}}\left(p_{\mathrm{f_a}}\left(1-p_{\mathrm{m}}\right) + \left(1-p_{\mathrm{f_a}}\right)p_{\mathrm{m}}\right) + p_{\mathrm{f_a}} p_{\mathrm{m}} w_{11}^{\mathrm{m}}}{w_{01}^{\mathrm{m}}\left(p_{\mathrm{f_a}}\left(1-p_{\mathrm{m}}\right) + \left(1-p_{\mathrm{f_a}}\right)p_{\mathrm{m}}\right) + w_{11}^{\mathrm{m}} p_{\mathrm{f_a}} p_{\mathrm{m}} + w_{00}^{\mathrm{m}}\left(1-p_{\mathrm{f_a}}\right)\left(1-p_{\mathrm{m}}\right)} \tag{S93c}$$

$$p'_{\mathrm{f_g}} = \hat{p}_{\mathrm{f_g}}(1-\mu_{\mathrm{f_g}}) + (1-\hat{p}_{\mathrm{f_g}})\mu_{\mathrm{f_g}} \approx \hat{p}_{\mathrm{f_g}} + \mu_{\mathrm{f_g}} \tag{S93d}$$

$$p'_{\mathrm{f_a}} = \hat{p}_{\mathrm{f_a}}(1-\mu_{\mathrm{f_a}}) + (1-\hat{p}_{\mathrm{f_a}})\mu_{\mathrm{f_a}} \approx \hat{p}_{\mathrm{f_a}} + \mu_{\mathrm{f_a}} \tag{S93e}$$

$$p'_{\mathrm{m}} = \hat{p}_{\mathrm{m}}(1-\mu_{\mathrm{m}}) + (1-\hat{p}_{\mathrm{m}})\mu_{\mathrm{m}} \approx \hat{p}_{\mathrm{m}} + \mu_{\mathrm{m}} \tag{S93f}$$

###### 4.1.5 X chromosome androgenic portion

$$\hat{p}_{\mathrm{f_g}} = \frac{w_1^{\mathrm{f_g}} p_{\mathrm{m}}}{w_1^{\mathrm{f_g}} + w_0^{\mathrm{f_g}}\left(1-p_{\mathrm{m}}\right)} \tag{S94a}$$

$$\hat{p}_{\mathrm{f_a}} = \frac{\frac{1}{2} w_{01}^{\mathrm{f_a}}\left(p_{\mathrm{f_g}}\left(1-p_{\mathrm{m}}\right) + \left(1-p_{\mathrm{f_g}}\right)p_{\mathrm{m}}\right) + p_{\mathrm{f_g}} p_{\mathrm{m}} w_{11}^{\mathrm{f_a}}}{w_{01}^{\mathrm{f_a}}\left(p_{\mathrm{f_g}}\left(1-p_{\mathrm{m}}\right) + \left(1-p_{\mathrm{f_g}}\right)p_{\mathrm{m}}\right) + w_{11}^{\mathrm{f_a}} p_{\mathrm{f_g}} p_{\mathrm{m}} + w_{00}^{\mathrm{f_a}}\left(1-p_{\mathrm{f_g}}\right)\left(1-p_{\mathrm{m}}\right)} \tag{S94b}$$

$$\hat{p}_{\mathrm{m}} = \frac{w_1^{\mathrm{m}} p_{\mathrm{f_a}}}{w_1^{\mathrm{m}} p_{\mathrm{f_a}} + w_0^{\mathrm{m}}\left(1-p_{\mathrm{f_a}}\right)} \tag{S94c}$$

$$p'_{\mathrm{f_g}} = \hat{p}_{\mathrm{f_g}}(1-\mu_{\mathrm{f_g}}) + (1-\hat{p}_{\mathrm{f_g}})\mu_{\mathrm{f_g}} \approx \hat{p}_{\mathrm{f_g}} + \mu_{\mathrm{f_g}} \tag{S94d}$$

$$p'_{\mathrm{f_a}} = \hat{p}_{\mathrm{f_a}}(1-\mu_{\mathrm{f_a}}) + (1-\hat{p}_{\mathrm{f_a}})\mu_{\mathrm{f_a}} \approx \hat{p}_{\mathrm{f_a}} + \mu_{\mathrm{f_a}} \tag{S94e}$$

$$p'_{\mathrm{m}} = \hat{p}_{\mathrm{m}}(1-\mu_{\mathrm{m}}) + (1-\hat{p}_{\mathrm{m}})\mu_{\mathrm{m}} \approx \hat{p}_{\mathrm{m}} + \mu_{\mathrm{m}} \tag{S94f}$$

###### 4.1.6 Paternal genome elimination androgenic portion

$$\hat{p}_{\mathrm{f_g}} = \frac{w_1^{\mathrm{f_g}} p_{\mathrm{m}}}{w_1^{\mathrm{f_g}} + w_0^{\mathrm{f_g}}\left(1-p_{\mathrm{m}}\right)} \tag{S95a}$$

$$\hat{p}_{\mathrm{f_a}} = \frac{\frac{1}{2} w_{01}^{\mathrm{f_a}}\left(p_{\mathrm{f_g}}\left(1-p_{\mathrm{m}}\right) + \left(1-p_{\mathrm{f_g}}\right)p_{\mathrm{m}}\right) + p_{\mathrm{f_g}} p_{\mathrm{m}} w_{11}^{\mathrm{f_a}}}{w_{01}^{\mathrm{f_a}}\left(p_{\mathrm{f_g}}\left(1-p_{\mathrm{m}}\right) + \left(1-p_{\mathrm{f_g}}\right)p_{\mathrm{m}}\right) + w_{11}^{\mathrm{f_a}} p_{\mathrm{f_g}} p_{\mathrm{m}} + w_{00}^{\mathrm{f_a}}\left(1-p_{\mathrm{f_g}}\right)\left(1-p_{\mathrm{m}}\right)} \tag{S95b}$$

$$\hat{p}_{\mathrm{m}} = \frac{w_{01}^{\mathrm{m}} p_{\mathrm{f_a}}\left(1-p_{\mathrm{m}}\right) + p_{\mathrm{f_a}} p_{\mathrm{m}} w_{11}^{\mathrm{m}}}{w_{01}^{\mathrm{m}}\left(p_{\mathrm{f_a}}\left(1-p_{\mathrm{m}}\right) + \left(1-p_{\mathrm{f_a}}\right)p_{\mathrm{m}}\right) + w_{11}^{\mathrm{m}} p_{\mathrm{f_a}} p_{\mathrm{m}} + w_{00}^{\mathrm{m}}\left(1-p_{\mathrm{f_a}}\right)\left(1-p_{\mathrm{m}}\right)} \tag{S95c}$$

$$p'_{f_g} = \hat{p}_{f_g}(1-\mu_{f_g}) + (1-\hat{p}_{f_g})\mu_{f_g} \approx \hat{p}_{f_g} + \mu_{f_g} \tag{S95d}$$

$$p'_{f_a} = \hat{p}_{f_a}(1-\mu_{f_a}) + (1-\hat{p}_{f_a})\mu_{f_a} \approx \hat{p}_{f_a} + \mu_{f_a} \tag{S95e}$$

$$p'_{m} = \hat{p}_{m}(1-\mu_{m}) + (1-\hat{p}_{m})\mu_{m} \approx \hat{p}_{m} + \mu_{m} \tag{S95f}$$

###### 4.1.7 Gynogenic portion

$$\hat{p}_{f_g} = \frac{w_1^{f_g} p_{f_g}}{w_1^{f_g} + w_0^{f_g}\left(1 - p_{f_g}\right)} \tag{S96a}$$

$$p'_{f_g} = \hat{p}_{f_g}(1-\mu_{f_g}) + (1-\hat{p}_{f_g})\mu_{f_g} \approx \hat{p}_{f_g} + \mu_{f_g} \tag{S96b}$$

##### 4.2 Mutation-selection balance and genetic load

Considering a mutation which exhibits a deleterious effect in class $i$ of $s_i$ when in it's hemi/homozygous state, and has a dominance coefficient of $h$. The mutation rate between the wildtype and the mutant in class $i$ is $\mu_i$. We trace the life-cycle starting in gametes, fertilisation occurs, selection, and then mutation. We will assume that selection occurs in the adult form, and so track the frequency of our mutation in the gametes of class $i$ as $p_i$. The allele frequency after selection is $\hat{p}_i$, and than after mutation is $p'_i = \hat{p}_i(1-\mu_i) + (1-\hat{p}_i)\mu_i \approx \hat{p}_i + \mu_i$, a good approximation provided mutation is weak.

Provided the mutation is not entirely recessive, then we can ignore terms of $O\left(p_i^2\right)$ and then write out our system of equations as:

$$\vec{p} = (\mathbf{I} - \mathbf{S})\mathbf{T}\vec{p} + \vec{u} \tag{S97}$$

Where $\vec{p}$ is the vector of allele frequencies in the gametes of the different classes, $\vec{u}$ is the vector of mutation rates in those different classes, $\mathbf{I}$ is the identity matrix, $\mathbf{S}$ is a diagonal matrix whose elements are the marginal fitness effects of the mutation when rare in the population $-a_i(0)$, and $\mathbf{T}$ is the backwards transition matrix. We can then solve these to get the frequency of deleterious mutations in the gametes of our different classes. These approximations can be seen in Table S6. In the case of X chromosomes and autosomes, these are equivalent to the expressions given by Werren (1993), however whilst those expressions are written in terms of allele frequencies in the zygotes, instead we express them here in terms of allele frequencies in the gametes. We compare these approximations to numerical solutions of the recursion equations. These are plotted alongside the approximations for the different genetic systems considered.

The genetic load imposed upon class $i$, $L_i$, can then be calculated by first calculating the equilibrium genotype frequencies from the equilibrium frequencies in gametes, and then multiplying these by the fitness effects costs experienced in those genotypes.

$$\vec{L} = (\mathbf{S}.\mathbf{G})\,\vec{p} \tag{S98}$$

Where $G$ is a matrix that converts the gamete frequencies to genotype frequencies, which assumes that our mutation is rare enough in the population that the genotypic frequency is simply a sum of the various gamete frequencies which contribute to that gene position. For example, for an autosomal locus:

$$\vec{L}_{\mathrm{A}} = \begin{pmatrix} hs_{\mathrm{f}} & 0 \\ 0 & hs_{\mathrm{m}} \end{pmatrix} \begin{pmatrix} 1 & 1 \\ 1 & 1 \end{pmatrix} \begin{pmatrix} p_{\mathrm{f}} \\ p_{\mathrm{m}} \end{pmatrix} = \begin{pmatrix} hs_{\mathrm{f}}(p_{\mathrm{f}} + p_{\mathrm{m}}) \\ hs_{\mathrm{m}}(p_{\mathrm{f}} + p_{\mathrm{m}}) \end{pmatrix} \tag{S99}$$

Or for an X-linked locus:

$$\vec{L}_{\mathrm{X}} = \begin{pmatrix} hs_{\mathrm{f}} & 0 \\ 0 & s_{\mathrm{m}} \end{pmatrix} \begin{pmatrix} 1 & 1 \\ 1 & 0 \end{pmatrix} \begin{pmatrix} p_{\mathrm{f}} \\ p_{\mathrm{m}} \end{pmatrix} = \begin{pmatrix} hs_{\mathrm{f}}(p_{\mathrm{f}} + p_{\mathrm{m}}) \\ s_{\mathrm{m}} p_{\mathrm{f}} \end{pmatrix} \tag{S100}$$

The values for the genetic load for our different genetic systems when making different assumptions about the expression patterns of these genes can be seen in Table S7. We can see that patterns of genetic load in PGE species distinct from both diploid and haplodiploid systems, in particular showing a much higher load for genes whose expression is male limited.

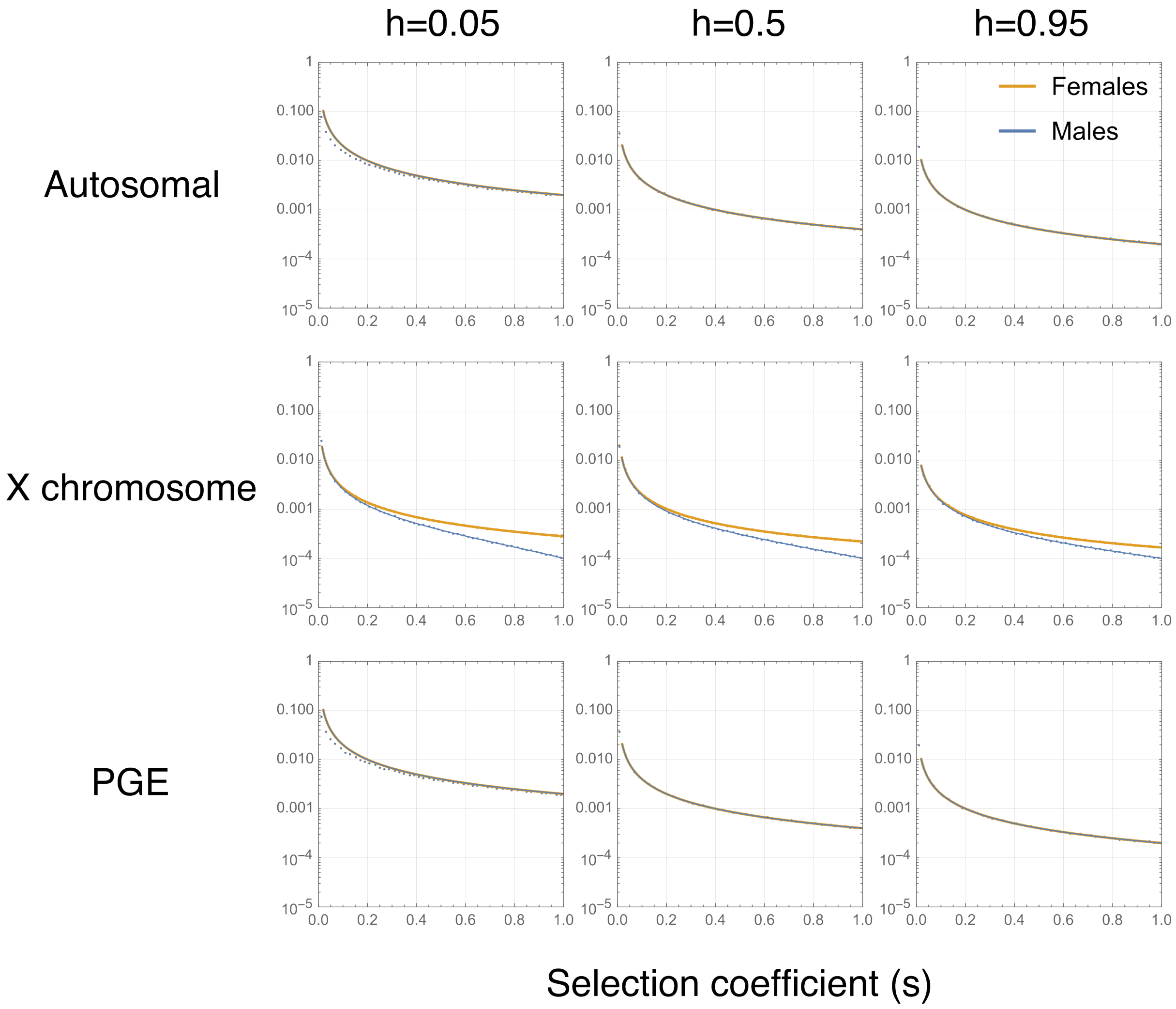


Figure S9: Deleterious allele frequency in different classes of individual, for different genetic systems, with different strengths of selection. The mutation is assumed to have equal fitness effects in males and females. Points are numerical solutions and solid lines are analytical approximations assuming the allele is rare in the population. The mutation rate is $\mu = 10^{-4}$.

Table S6: Allele frequency in different classes under mutation selection balance, where the mutation rate is equal in all classes $\mu$. The allele frequencies here are given in terms of the gametes, rather than in zygotes.

| | Class | All | Female-limited | Male-limited |
|---|---|---|---|---|
| A | $p_{\mathrm{f}}$ | $\mu\left(\frac{1}{hs}\right)$ | $\mu\left(\frac{2-hs}{hs}\right)$ | $\mu\left(\frac{2+hs}{hs}\right)$ |
| | $p_{\mathrm{m}}$ | $\mu\left(\frac{1}{hs}\right)$ | $\mu\left(\frac{2+hs}{hs}\right)$ | $\mu\left(\frac{2-hs}{hs}\right)$ |
| X | $p_{\mathrm{f}}$ | $\mu\left(\frac{3-hs}{s(1-h(2-s))}\right)$ | $\mu\left(\frac{3-hs}{2hs}\right)$ | $\mu\left(\frac{3}{s}\right)$ |
| | $p_{\mathrm{m}}$ | $\mu\left(\frac{(3-(2-h)s)}{s(1-h(2-s))}\right)$ | $\mu\left(\frac{(3+hs)}{2hs}\right)$ | $\mu\left(\frac{3-2s}{s}\right)$ |
| PGE | $p_{\mathrm{f}}$ | $\mu\left(\frac{1}{hs}\right)$ | $\mu\left(\frac{3-hs}{2hs}\right)$ | $\mu\left(\frac{3}{hs}\right)$ |
| | $p_{\mathrm{m}}$ | $\mu\left(\frac{1}{hs}\right)$ | $\mu\left(\frac{(3+hs)}{2hs}\right)$ | $\mu\left(\frac{3-2hs}{hs}\right)$ |
| $A_{\mathbb{a}}$ | $p_{\mathrm{f}_{\mathrm{g}}}$ | $\mu\left(\frac{7-s(6-h(2+(2-h)s))}{s+hs(6-s(2+h(2-s)))}\right)$ | $\mu\left(\frac{7-(6-h)s}{s(1+h(2-s))}\right)$ | $\mu\left(\frac{7+hs}{4hs}\right)$ |
| | $p_{\mathrm{f}_{\mathbb{a}}}$ | $\mu\left(\frac{7-s(2+h(2-(2-h)s))}{s+hs(6-s(2+h(2-s)))}\right)$ | $\mu\left(\frac{(7-s(2+h(5-2s)))}{s(1+h(2-s))}\right)$ | $\mu\left(\frac{7+3hs}{4hs}\right)$ |
| | $p_{\mathrm{m}}$ | $\mu\left(\frac{7-hs(4-hs)}{s+hs(6-s(2+h(2-s)))}\right)$ | $\mu\left(\frac{7-hs}{s(1+h(2-s))}\right)$ | $\mu\left(\frac{7-3hs}{4hs}\right)$ |
| $X_{\mathbb{a}}$ | $p_{\mathrm{f}_{\mathrm{g}}}$ | $\mu\left(\frac{5-s(5-2s-h(1-s))}{s(3-s+h(2-s)(1-s))}\right)$ | $\mu\frac{(5-s(4-h))}{s(1+h(2-s))}$ | $\mu\left(\frac{5-s}{2s}\right)$ |
| | $p_{\mathrm{f}_{\mathbb{a}}}$ | $\mu\left(\frac{5-s(1+h(3-s))}{s(3-s+h(2-s)(1-s))}\right)$ | $\mu\left(\frac{5-s-hs(3-s)}{s(1+h(2-s))}\right)$ | $\mu\left(\frac{5}{2s}\right)$ |
| | $p_{\mathrm{m}}$ | $\mu\left(\frac{5-s(3+h(1-s))}{s(3-s+h(2-s)(1-s))}\right)$ | $\mu\left(\frac{5-hs}{s(1+h(2-s))}\right)$ | $\mu\left(\frac{5-3s}{2s}\right)$ |
| $PGE_{\mathbb{a}}$ | $p_{\mathrm{f}_{\mathrm{g}}}$ | $\mu\left(\frac{(5-s(4-(2-h)hs))}{s+h(2-s)s(2-hs)}\right)$ | $\mu\left(\frac{5-(4-h)s}{(1+h(2-s))s}\right)$ | $\mu\left(\frac{(5-\mathrm{hs})}{2hs}\right)$ |
| | $p_{\mathrm{f}_{\mathbb{a}}}$ | $\mu\left(\frac{(5-(1+h(3-s))s)}{s+h(2-s)s(2-hs)}\right)$ | $\mu\left(\frac{(5-s-hs(3-s))}{(1+h(2-s))s}\right)$ | $\mu\left(\frac{5}{2hs}\right)$ |
| | $p_{\mathrm{m}}$ | $\mu\left(\frac{(5-hs(4-hs))}{s+h(2-s)s(2-hs)}\right)$ | $\mu\left(\frac{5-hs}{(1+h(2-s))s}\right)$ | $\mu\left(\frac{5-3hs}{2hs}\right)$ |

Table S7: The genetic load imposed upon different classes and under different genetic systems, and patterns of gene expression.

| | Class | Equal Expression | Female-limited | Male-limited |
|---|---|---|---|---|
| Autosomal | $L_f$ | $2\mu$ | $4\mu$ | 0 |
| | $L_m$ | $2\mu$ | 0 | $4\mu$ |
| X | $L_f$ | $\frac{2h\mu(s-3)}{h(s-2)-1}$ | $3\mu$ | 0 |
| | $L_m$ | $\frac{\mu(hs-3)}{h(s-2)-1}$ | 0 | $3\mu$ |
| PGE | $L_f$ | $2\mu$ | $3\mu$ | 0 |
| | $L_m$ | $2\mu$ | 0 | $-2\mu(hs-3)$ |
| $A_{\mathbb{a}}$ | $L_{f_{\mathbf{g}}}$ | $\frac{\mu(h^2s^2-4hs+7)}{h^2(s-2)s-2h(s-3)+1}$ | $\frac{\mu(hs-7)}{h(s-2)-1}$ | 0 |
| | $L_{f_{\mathbb{a}}}$ | $\frac{2h\mu(hs^2-(h+3)s+7)}{h^2(s-2)s-2h(s-3)+1}$ | $\frac{2h\mu(3s-7)}{h(s-2)-1}$ | 0 |
| | $L_m$ | $\frac{2h\mu(hs^2-(3h+1)s+7)}{h^2(s-2)s-2h(s-3)+1}$ | 0 | $\frac{7\mu}{2}$ |
| $X_{\mathbb{a}}$ | $L_{f_{\mathrm{g}}}$ | $\frac{\mu(hs^2-(h+3)s+5)}{h(s^2-3s+2)-s+3}$ | $\frac{\mu(hs-5)}{h(s-2)-1}$ | 0 |
| | $L_{f_{\mathbb{a}}}$ | $\frac{2h\mu(s^2-4s+5)}{h(s^2-3s+2)-s+3}$ | $\frac{2h\mu(2s-5)}{h(s-2)-1}$ | 0 |
| | $L_m$ | $\frac{\mu(hs^2-(3h+1)s+5)}{h(s^2-3s+2)-s+3}$ | 0 | $\frac{5\mu}{2}$ |
| $X_{\mathbb{a}}$ | $L_{f_{\mathrm{g}}}$ | $\frac{\mu(h^2s^2-4hs+5)}{h^2(s-2)s-2h(s-2)+1}$ | $\frac{\mu(hs-5)}{h(s-2)-1}$ | 0 |
| | $L_{f_{\mathbb{a}}}$ | $\frac{2h\mu(hs^2-2(h+1)s+5)}{h^2(s-2)s-2h(s-2)+1}$ | $\frac{2h\mu(2s-5)}{h(s-2)-1}$ | 0 |
| | $L_m$ | $\frac{h\mu(h(h+1)s^2-(7h+1)s+10)}{h^2(s-2)s-2h(s-2)+1}$ | 0 | $\mu\left(5-\frac{3hs}{2}\right)$ |

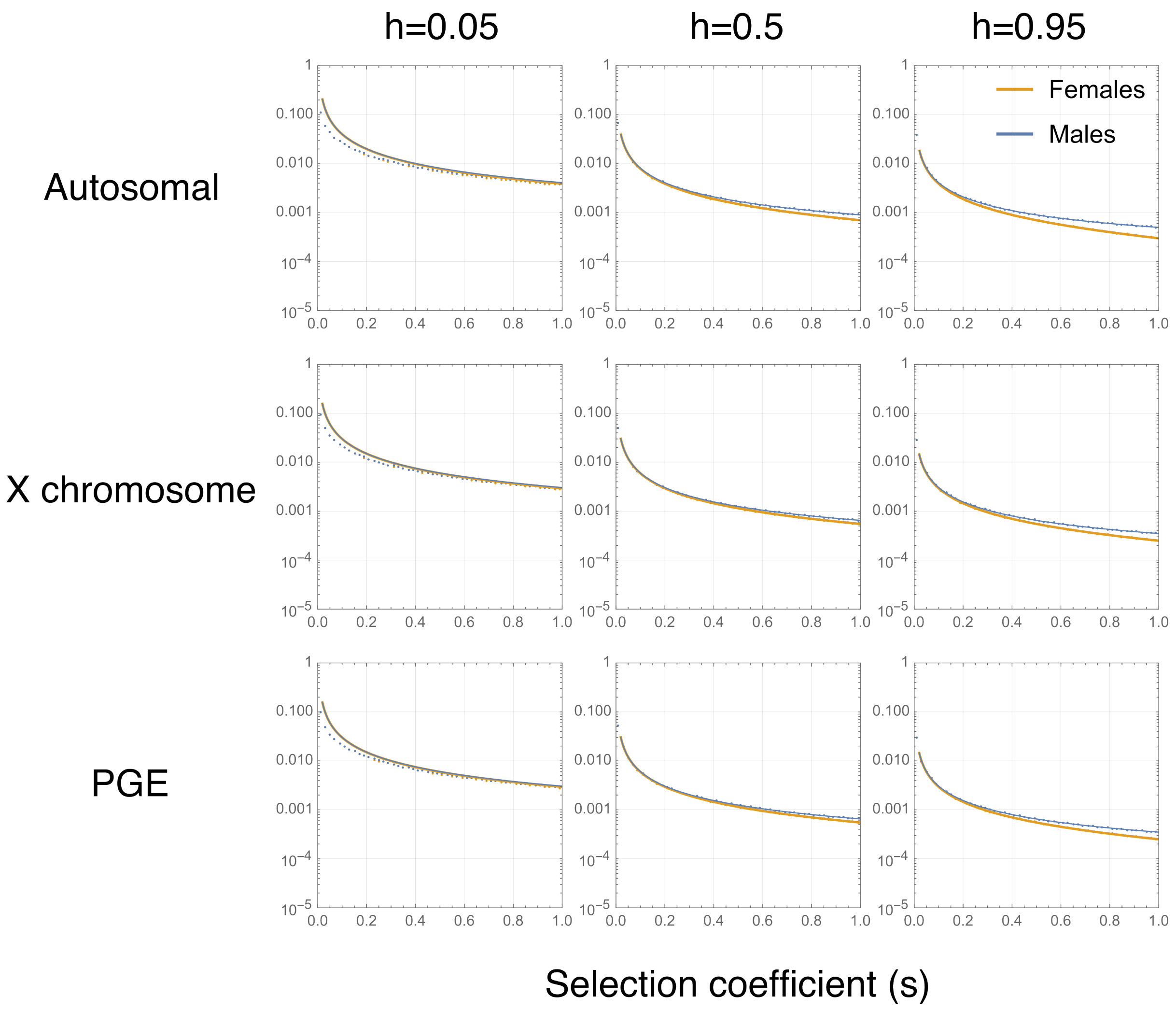


Figure S10: Deleterious allele frequency in different classes of individual, for different genetic systems, with different strengths of selection. The mutation is assumed to have fitness effects only in females. Points are numerical solutions and solid lines are analytical approximations assuming the allele is rare in the population. The mutation rate is $\mu = 10^{-4}$.

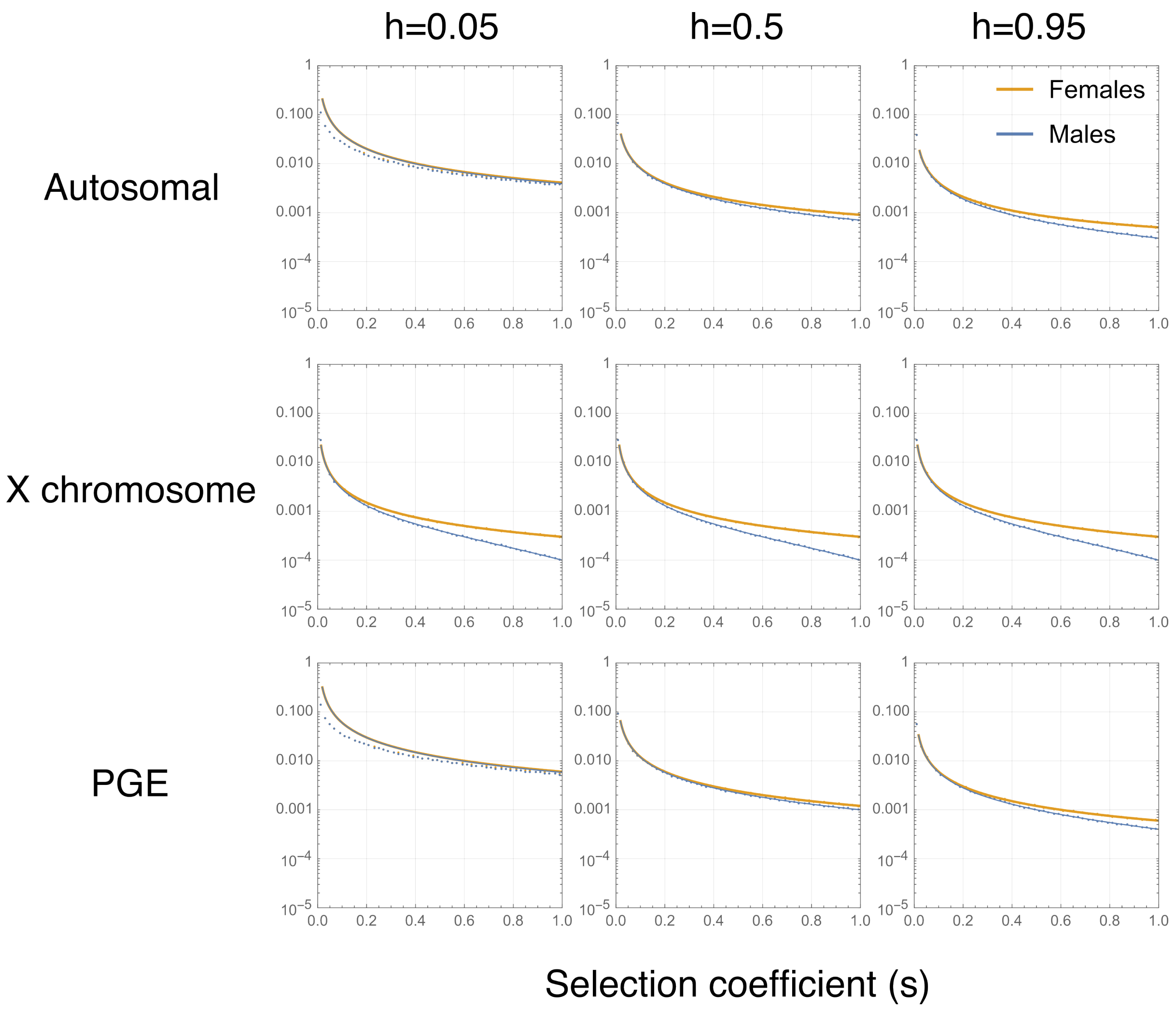


Figure S11: Deleterious allele frequency in different classes of individual, for different genetic systems, with different strengths of selection. The mutation is assumed to have fitness effects only in males. Points are numerical solutions and solid lines are analytical approximations assuming the allele is rare in the population. The mutation rate is $\mu = 10^{-4}$.

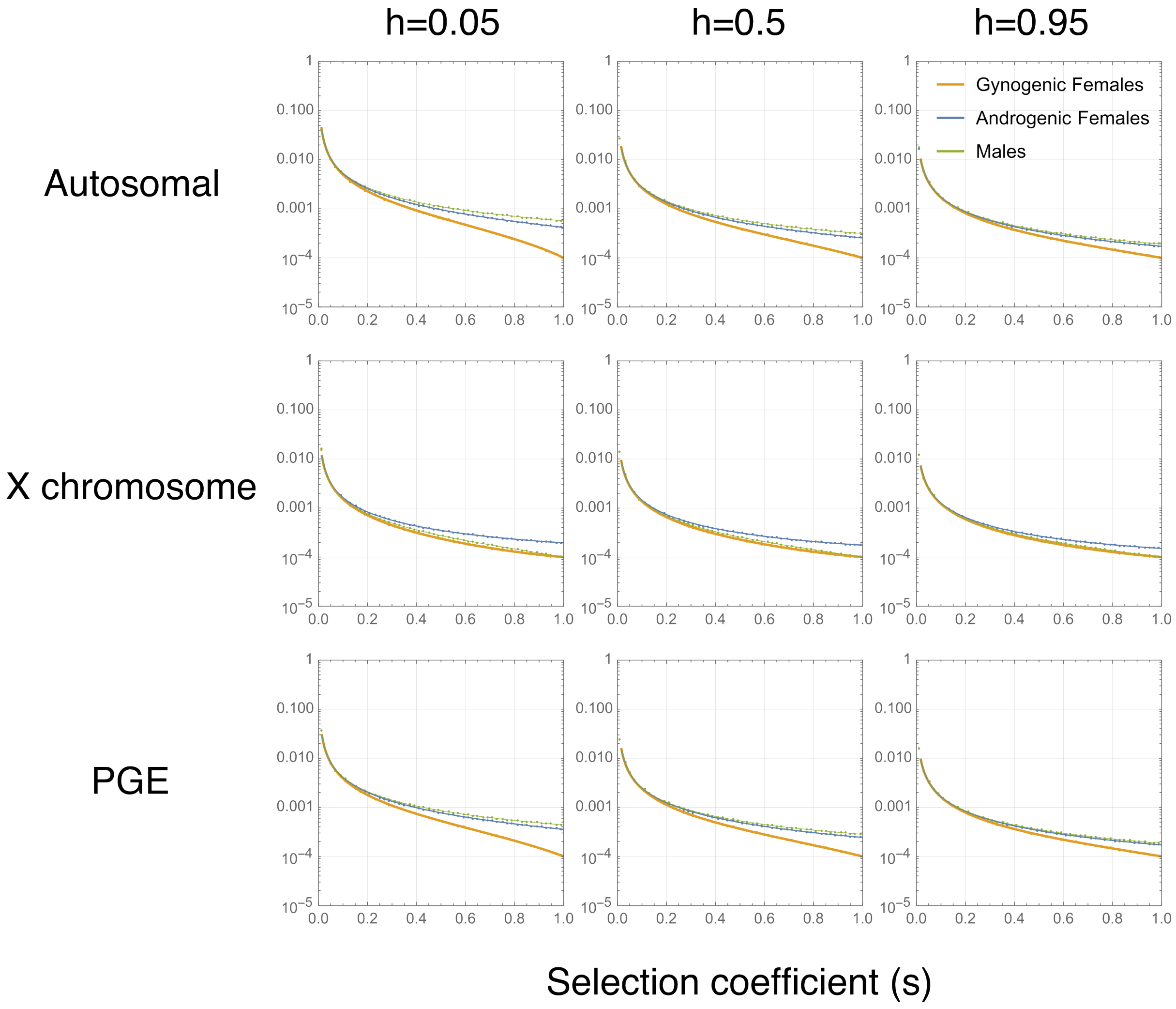


Figure S12: Deleterious allele frequency in different classes of individual, for different genetic systems, with different strengths of selection. Here the plotted genetic systems are the androgenic portion of either the autosome, X chromosome, or PGE autosome. The mutation is assumed to have equal fitness effects in all classes of individual. Points are numerical solutions and solid lines are analytical approximations assuming the allele is rare in the population. The mutation rate is $\mu = 10^{-4}$.

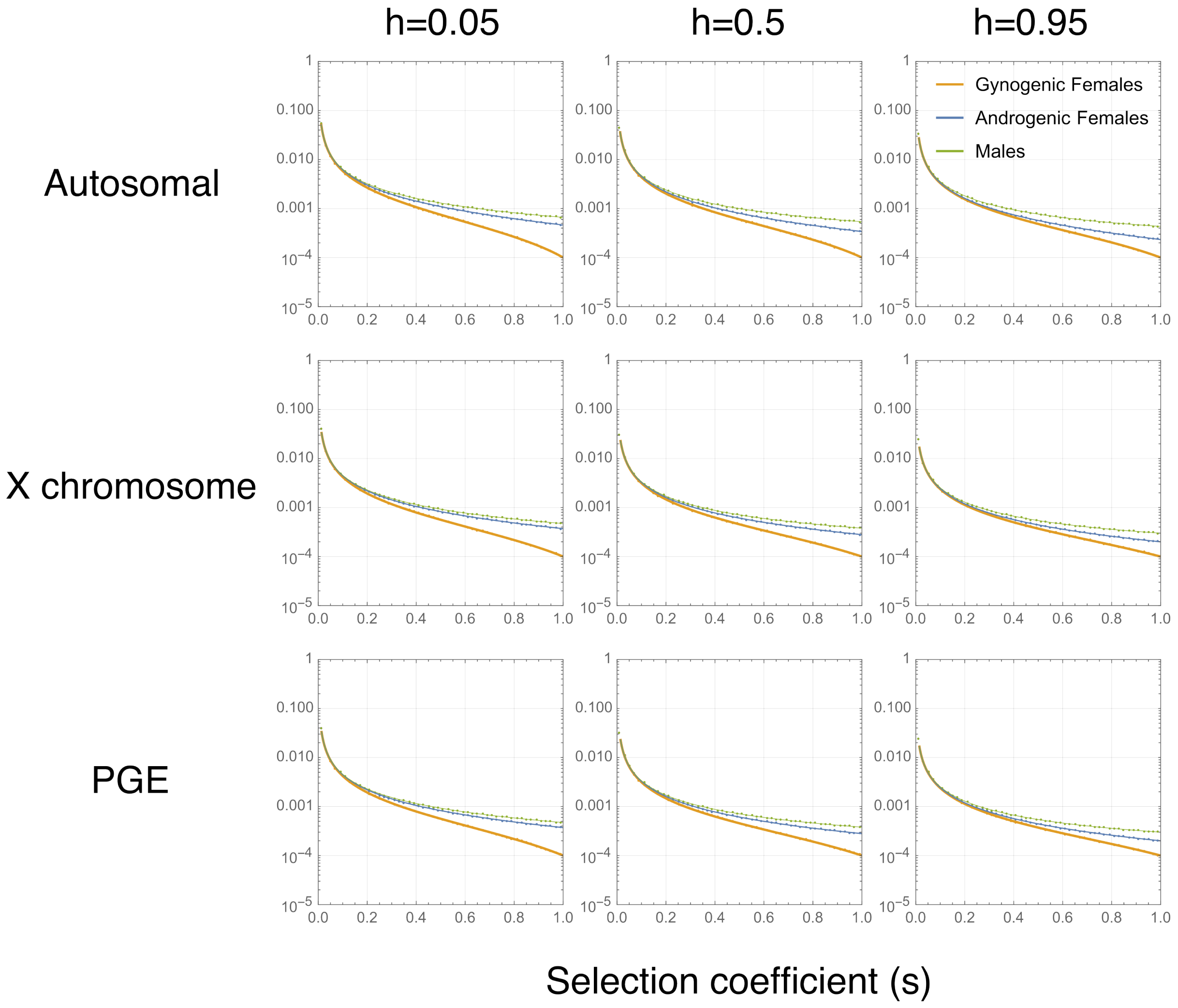


Figure S13: Deleterious allele frequency in different classes of individual, for different genetic systems, with different strengths of selection. Here the plotted genetic systems are the androgenic portion of either the autosome, X chromosome, or PGE autosome. The mutation is assumed to have fitness effects only in the female classes. Points are numerical solutions and solid lines are analytical approximations assuming the allele is rare in the population. The mutation rate is $\mu = 10^{-4}$.

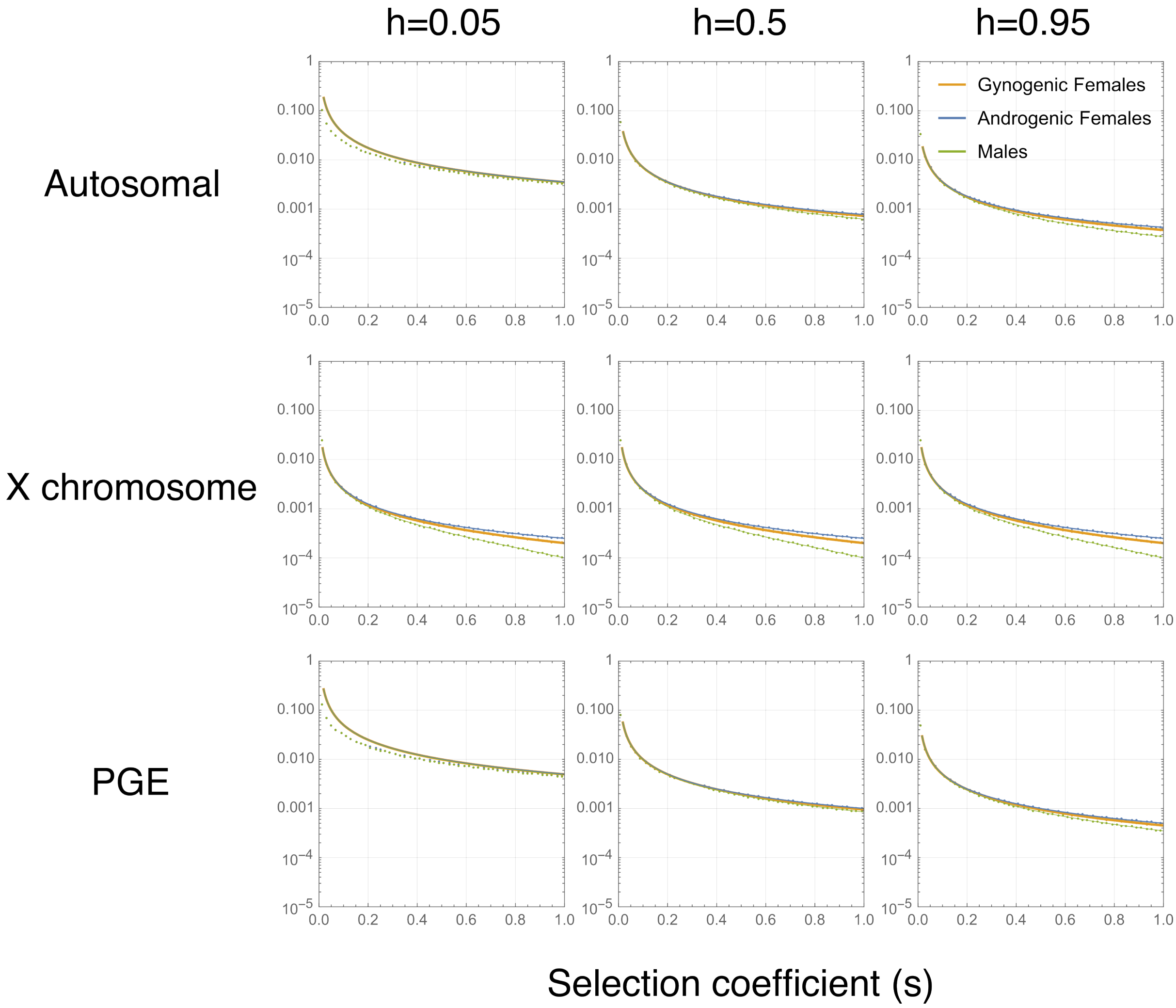


Figure S14: Deleterious allele frequency in different classes of individual, for different genetic systems, with different strengths of selection. Here the plotted genetic systems are the androgenic portion of either the autosome, X chromosome, or PGE autosome. The mutation is assumed to have fitness effects only in males. Points are numerical solutions and solid lines are analytical approximations assuming the allele is rare in the population. The mutation rate is $\mu = 10^{-4}$.

#### 4.3 Balancing selection

Finally, we consider how balancing selection may maintain genetic variation within populations. We consider two types, first sexual antagonism, whereby alleles that are beneficial when expressed in one sex are deleterious in the others. Second, we consider "maternal"-antagonism, where alleles have opposing fitness effects on gynogenic and androgenic females. The fitness schemes for these scenarios can be seen in Tables S8 and S9.

With the recursion equations written in section 4.1 and ignoring the effects of mutation, we consider whether a mutant allele will be able to invade from rarity. To do this we consider the stability of the two monomorphic equilibria, at $p = 0$ and $p = 1$, by calculating the Jacobian matrix, analysed at these equilbrium points.

$$\mathbf{J}_{ij} = \left.\frac{\partial p'_i}{\partial p'_j}\right|_{p=0} \tag{S101}$$

If the leading eigenvalue of this matrix is greater than one, $\lambda_{max} > 1$, then the mutant will increase in frequency, and thus be able to invade. If both of the monomorphic equilibria are unstable, then there will be a stable polymorphism. The boundary conditions for sexually antagonistic alleles can be seen in Table S10, and for maternally antagonistic alleles can be seen in Table S12. Additionally, these boundaries are plotted for some of the different systems, under different assumptions about dominance.

In addition, if we assume that selection is weak, then we can express the conditions for polymorphism as:

$$I_0 = \tilde{a}(0) > 1 \tag{S102}$$

$$I_1 = -(\tilde{a}(1)) > 1 \tag{S103}$$

i.e. the allele can invade, but it cannot fix in the population. As shown before, we can use these invasion boundaries to compute the equilibrium allele frequency, if such a polymorphism is maintained.

$$p* = \frac{I_0}{I_0 + I_1} \tag{S104}$$

These polymorphic conditions can be see in Table S11 and S13. Finally, we can consider the two invasion boundaries as two vectors in the plane of either $s_f$ and $s_m$ if sexually antagonistic, or $s_{\mathbb{g}}$ and $s_{\mathbb{a}}$ if maternally antagonistic. Calculating the angle between these two vectors provides a measure of the space for polymorphism. The angle $\theta$, between the vectors $a$ and $b$ will satisfy:

$$Cos(\theta) = \frac{a.b}{||a||\,||b||} \tag{S105}$$

These expressions for out different systems are plotted in the main text.

Table S8: Fitness scheme for sexually antagonistic selection.

| | 00/0 | 01/10 | 11/1 |
|---|---|---|---|
| $w_{f_\mathbb{g}}$ | 1 | $1-h_f s_f$ | $1-s_f$ |
| $w_{f_\mathbb{a}}$ | 1 | $1-h_f s_f$ | $1-s_f$ |
| $w_m$ | $1-s_m$ | $1-h_m s_m$ | 1 |

Table S9: Fitness scheme for maternally antagonistic selection.

| | 00/0 | 01/10 | 11/1 |
|---|---|---|---|
| $w_{f_\mathbb{g}}$ | $1-s_\mathbb{g}$ | $1-h_\mathbb{g} s_\mathbb{g}$ | 1 |
| $w_{f_\mathbb{a}}$ | 1 | $1-h_\mathbb{a} s_\mathbb{a}$ | $1-s_\mathbb{a}$ |
| $w_m$ | 1 | 1 | 1 |

Table S10: Invasion conditions for sexually antagonistic alleles on different portions of the genome.

| | m+/f- invades | f+/m- invades |
|---|---|---|
| Autosome | $s_m > \frac{h_f s_f}{h_f s_f - h_m + 1}$ | $s_m < \frac{(h_f-1) s_f}{(s_f-1) h_m}$ |
| X chromosome | $s_m > \frac{2 h_f s_f}{h_f s_f + 1}$ | $s_m < \frac{2(h_f-1) s_f}{h_f s_f - 1}$ |
| PGE | $s_m > \frac{2 h_f s_f}{h_f s_f h_m + h_f s_f - h_m + 1}$ | $s_m < \frac{2(h_f-1) s_f}{h_m (h_f s_f - 1)}$ |
| $A_\mathbb{a}$ | $s_m > \frac{s_f (h_f (s_f-2)-1)}{h_m \left(h_f s_f^2 - (2h_f+1) s_f + 4\right) - 4}$ | $s_m < \frac{s_f (h_f (s_f-2) - 2s_f + 3)}{h_m \left((h_f+2) s_f^2 - (2h_f+5) s_f + 4\right)}$ |
| $X_\mathbb{a}$ | $s_m > \frac{1}{2}\left(s_f - h_f (s_f - 2) s_f\right)$ | $s_m < \frac{s_f (h_f (s_f-2) - 2s_f + 3)}{(s_f-2)(h_f s_f - 1)}$ |
| PGE$_\mathbb{a}$ | $s_m > \frac{s_f (h_f (s_f-2)-1)}{(s_f-2) h_m (h_f s_f - 1) - 2}$ | $s_m < \frac{s_f (h_f (s_f-2) - 2s_f + 3)}{(s_f-2) h_m (h_f s_f - 1)}$ |

Table S11: Invasion conditions for sexually antagonistic alleles assuming weak selection on different portions of the genome, and the frequency of the male-beneficial allele if a stable polymorphism is obtained.

| | m+/f- invades | m+/f- fixes | Polymorphic Equilibrium |
|---|---|---|---|
| Autosome | $s_m > -\frac{h_f s_f}{h_m - 1}$ | $s_m > -\frac{(h_f-1) s_f}{h_m}$ | $\frac{h_f s_f + (h_m-1) s_m}{2\left(\frac{1}{2}(h_f s_f + (h_m-1) s_m) + \frac{1}{2}((h_f-1) s_f + h_m s_m)\right)}$ |
| X chromosome | $s_m > 2 h_f s_f$ | $s_m > -2\left(h_f - 1\right) s_f$ | $\frac{2 h_f s_f - s_m}{3\left(\frac{1}{3}(2 h_f s_f - s_m) + \frac{1}{3}(2(h_f-1) s_f + s_m)\right)}$ |
| PGE | $s_m > -\frac{2 h_f s_f}{h_m - 1}$ | $s_m > -\frac{2(h_f-1) s_f}{h_m}$ | $\frac{2 h_f s_f + (h_m-1) s_m}{3\left(\frac{1}{3}(2 h_f s_f + (h_m-1) s_m) + \frac{1}{3}(2(h_f-1) s_f + h_m s_m)\right)}$ |
| $A_\mathbf{a}$ | $s_m > \frac{-2 h_f s_f - s_f}{4(h_m-1)}$ | $s_m > -\frac{(2h_f-3) s_f}{4 h_m}$ | $\frac{2 h_f s_f + s_f + 4(h_m-1) s_m}{7\left(\frac{1}{7}(2 h_f s_f + s_f + 4(h_m-1) s_m) + \frac{1}{7}((2h_f-3) s_f + 4 h_m s_m)\right)}$ |
| $X_\mathbf{a}$ | $s_m > \frac{1}{2}\left(2 h_f s_f + s_f\right)$ | $s_m > -\frac{1}{2}\left(2 h_f - 3\right) s_f$ | $\frac{2 h_f s_f + s_f - 2 s_m}{5\left(\frac{1}{5}(2 h_f s_f + s_f - 2 s_m) + \frac{2}{5}((h_f - \frac{3}{2}) s_f + s_m)\right)}$ |
| PGE$_\mathbf{a}$ | $s_m > \frac{-2 h_f s_f - s_f}{2(h_m-1)}$ | $s_m > -\frac{(2h_f-3) s_f}{2 h_m}$ | $\frac{2 h_f s_f + s_f + 2(h_m-1) s_m}{5\left(\frac{1}{5}(2 h_f s_f + s_f + 2(h_m-1) s_m) + \frac{1}{5}((2h_f-3) s_f + 2 h_m s_m)\right)}$ |

Table S12: Invasion conditions for maternally antagonistic alleles on different portions of the genome.

| | $f_g$+/ $f_a$- invades | $f_a$+/ $f_g$- invades |
|---|---|---|
| Autosome | $s_g > \frac{h_a s_a}{-h_g + h_a s_a + 1}$ | $s_g < \frac{(h_a - 1) s_a}{h_g (s_a - 1)}$ |
| X chromosome | $s_g > \frac{h_a s_a}{-h_g + h_a s_a + 1}$ | $s_g < \frac{(h_a - 1) s_a}{h_g (s_a - 1)}$ |
| PGE | $s_g > \frac{h_a s_a}{-h_g + h_a s_a + 1}$ | $s_g < \frac{(h_a - 1) s_a}{h_g (s_a - 1)}$ |
| $A_a$ | $s_g > \frac{2 h_a s_a}{h_a s_a + 1}$ | $s_g < \frac{2(h_a - 1) s_a}{h_a s_a - 1}$ |
| $X_a$ | $s_g > \frac{2 h_a s_a}{h_a s_a + 1}$ | $s_g < \frac{2(h_a - 1) s_a}{h_a s_a - 1}$ |
| $PGE_a$ | $s_g > \frac{2 h_a s_a}{h_a s_a + 1}$ | $s_g < \frac{2(h_a - 1) s_a}{h_a s_a - 1}$ |

Table S13: Invasion conditions for maternally antagonistic alleles assuming weak selection on different portions of the genome, and the frequency of the gynogenic female-beneficial allele if a stable polymorphism is obtained,

| | $f_{\mathbf{g}}$+/ $f_{\mathbf{a}}$- invades | $f_{\mathbf{g}}$+/ $f_{\mathbf{a}}$- fixes | Polymorphic Equilibrium |
|---|---|---|---|
| Autosome | $s_g > -\frac{h_a s_a}{h_g - 1}$ | $s_g > -\frac{(h_a - 1) s_a}{h_g}$ | $\frac{h_a s_a + (h_g - 1) s_g}{4\left(\frac{1}{4}(h_a s_a + (h_g - 1) s_g) + \frac{1}{4}((h_a - 1) s_a + h_g s_g)\right)}$ |
| X chromosome | $s_g > -\frac{h_a s_a}{h_g - 1}$ | $s_g > -\frac{(h_a - 1) s_a}{h_g}$ | $\frac{h_a s_a + (h_g - 1) s_g}{3\left(\frac{1}{3}(h_a s_a + (h_g - 1) s_g) + \frac{1}{3}((h_a - 1) s_a + h_g s_g)\right)}$ |
| PGE | $s_g > -\frac{h_a s_a}{h_g - 1}$ | $s_g > -\frac{(h_a - 1) s_a}{h_g}$ | $\frac{h_a s_a + (h_g - 1) s_g}{3\left(\frac{1}{3}(h_a s_a + (h_g - 1) s_g) + \frac{1}{3}((h_a - 1) s_a + h_g s_g)\right)}$ |
| $A_a$ | $s_g > 2 h_a s_a$ | $s_g > -2 (h_a - 1) s_a$ | $\frac{2 h_a s_a - s_g}{7\left(\frac{1}{7}(2 h_a s_a - s_g) + \frac{1}{7}(2(h_a - 1) s_a + s_g)\right)}$ |
| $X_a$ | $s_g > 2 h_a s_a$ | $s_g > -2 (h_a - 1) s_a$ | $\frac{2 h_a s_a - s_g}{5\left(\frac{1}{5}(2 h_a s_a - s_g) + \frac{1}{5}(2(h_a - 1) s_a + s_g)\right)}$ |
| $PGE_a$ | $s_g > 2 h_a s_a$ | $s_g > -2 (h_a - 1) s_a$ | $\frac{2 h_a s_a - s_g}{5\left(\frac{1}{5}(2 h_a s_a - s_g) + \frac{1}{5}(2(h_a - 1) s_a + s_g)\right)}$ |

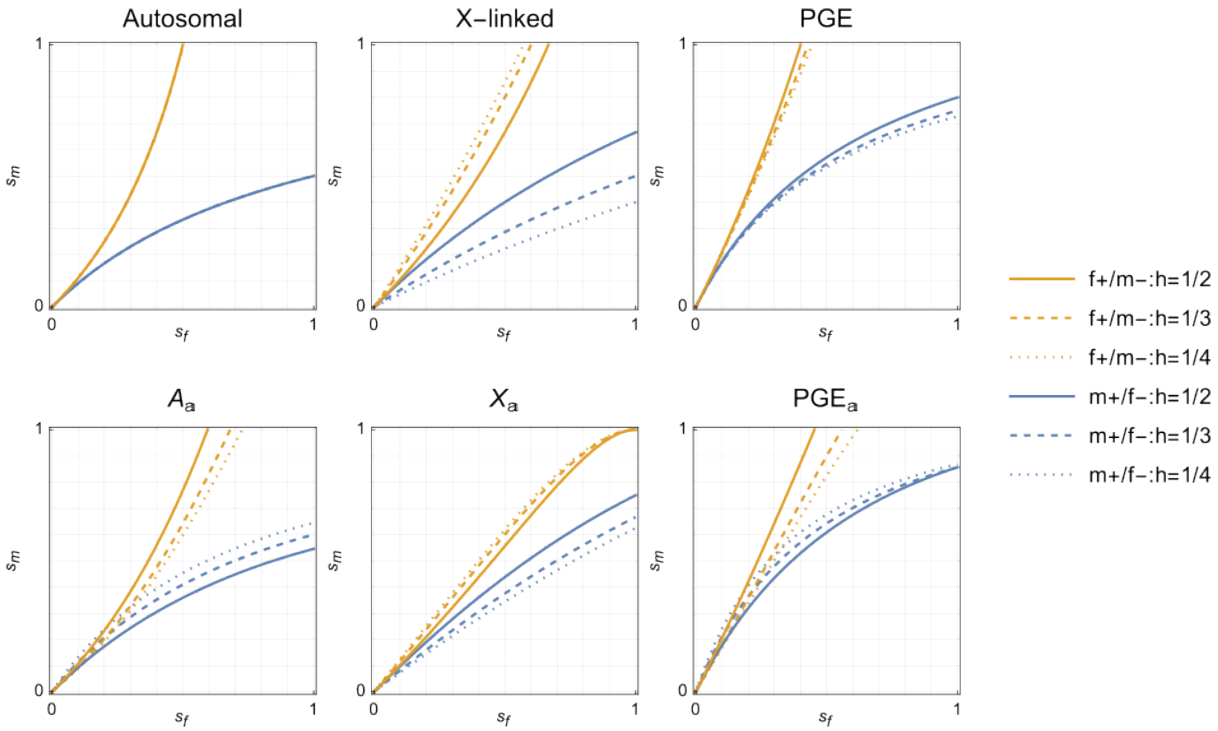


Figure S15: Boundary conditions for sexually antagonistic alleles under different genetic systems where dominance is assumed to be equal between males and females, such that $h_\mathrm{f} = h$ and $h_\mathrm{m} = 1 - h$.

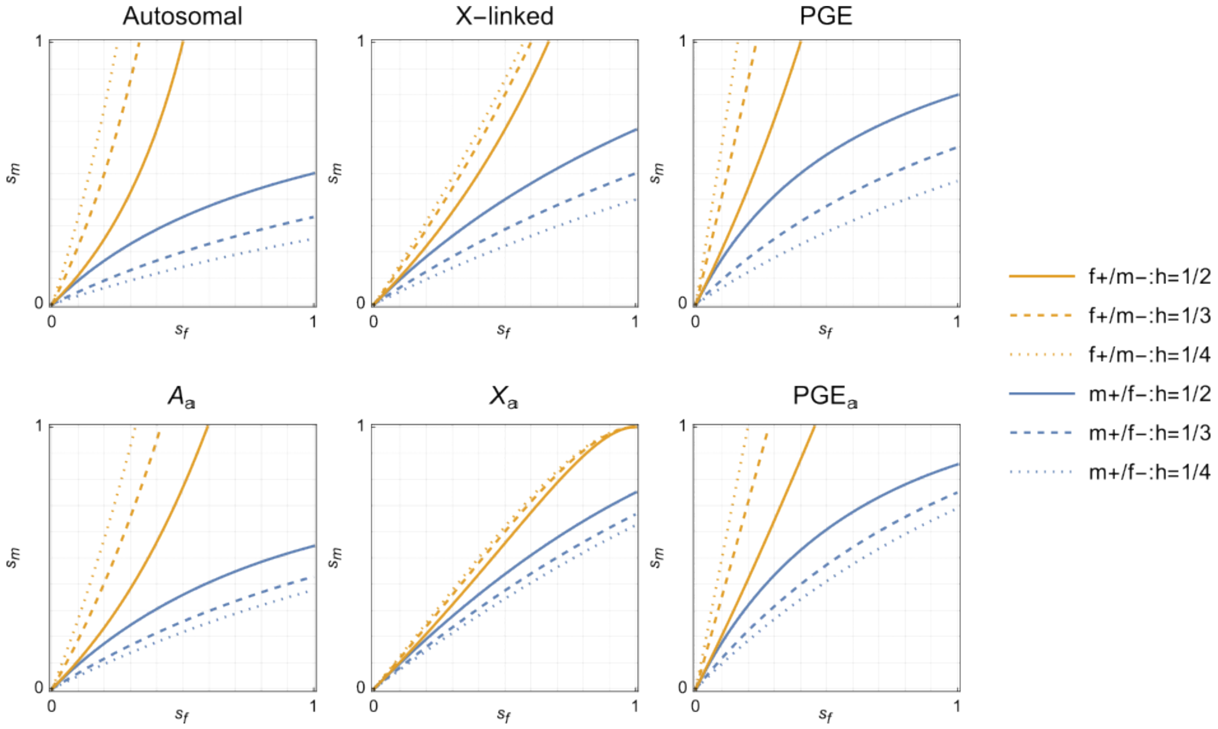


Figure S16: Boundary conditions for sexually antagonistic alleles under different genetic systems where reversals of dominance are assumed between males and females, such that $h_\mathrm{f} = h$ and $h_\mathrm{m} = h$.

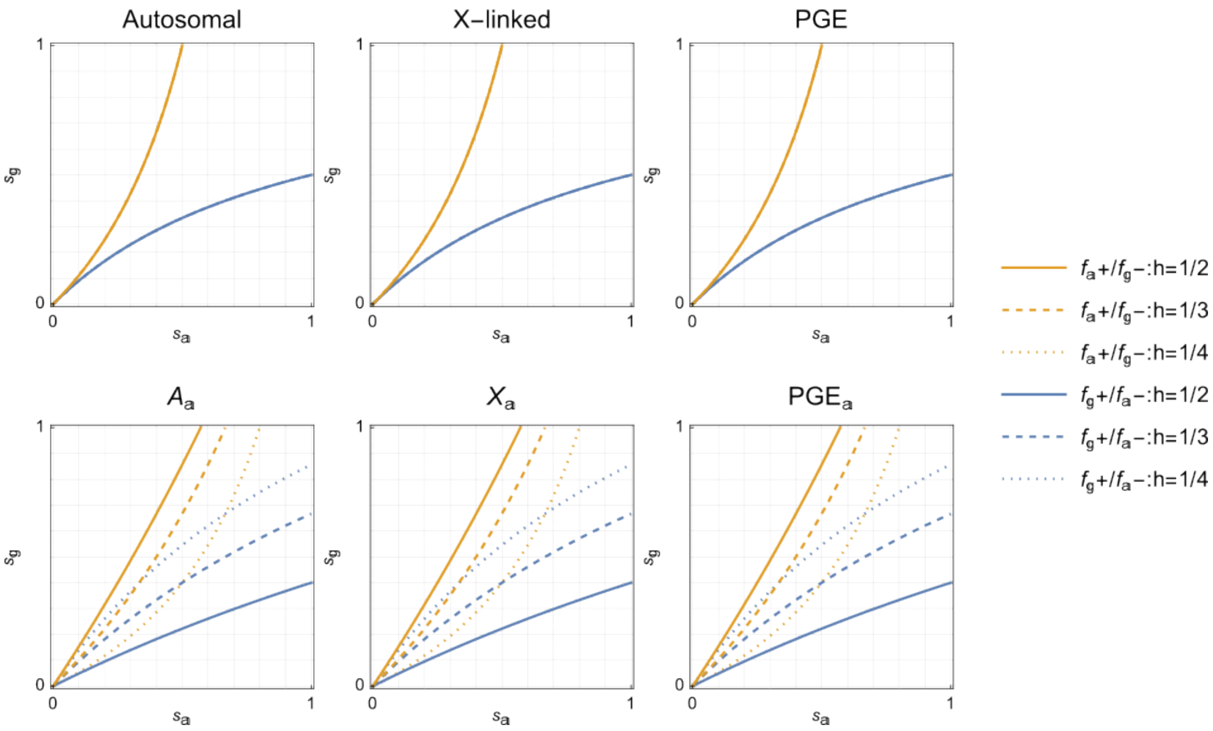


Figure S17: Boundary conditions for maternally antagonistic alleles under different genetic systems where dominance is assumed to be equal between gynogenic and androgenic females, such that $h_{f_a} = h$ and $h_{f_g} = 1 - h$.

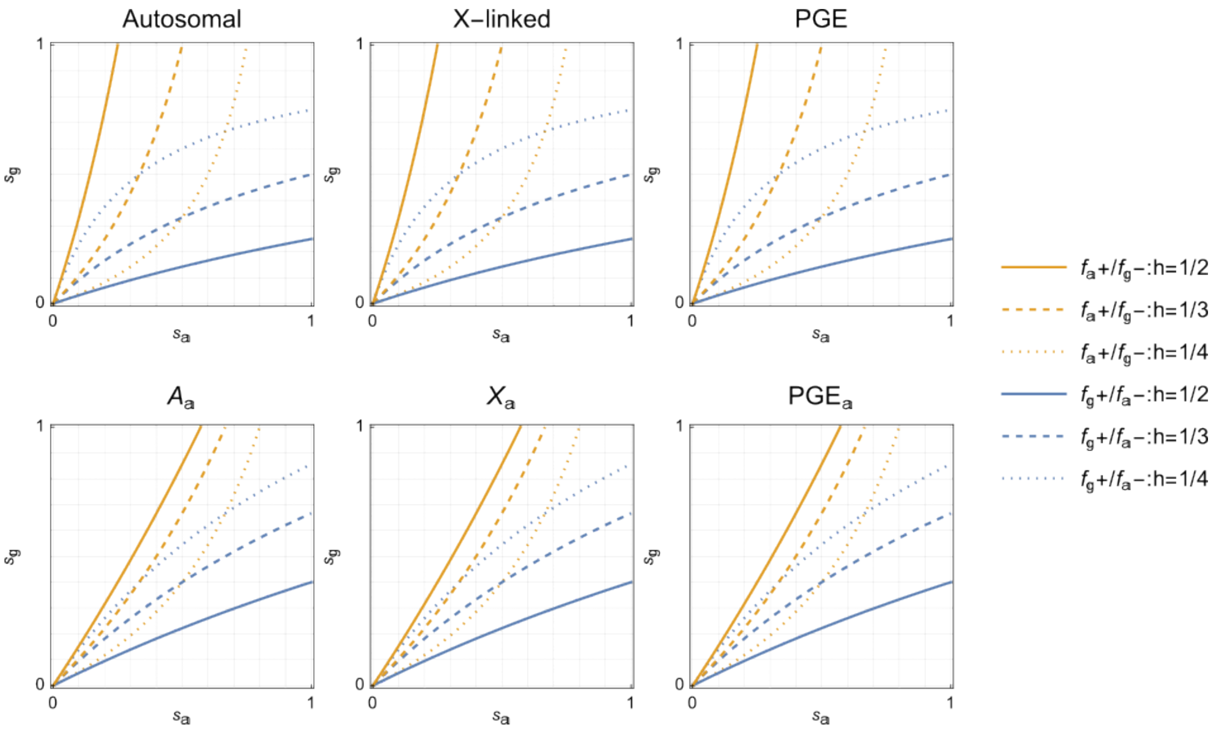


Figure S18: Boundary conditions for maternally antagonistic alleles under different genetic systems where reversals of dominance are between gynogenic and androgenic females, such that $h_{f_a} = h$ and $h_{f_g} = h$.
